# Two axolotl-adapted cell-ablation platforms reveal macrophage-dependent processes essential for spinal-cord and skeletal regeneration

**DOI:** 10.64898/2026.02.10.705144

**Authors:** Gabriela Johnson, Andrew Hart, Markus Sujansky, Joel H. Graber, James W. Godwin

## Abstract

The axolotl (*Ambystoma mexicanum*) has emerged as the premier model organism for studying scarless repair and adult tissue regeneration, supported by an expanding collection of tissue-specific transgenic lines and translucent skin that enables high-quality live imaging and cell tracking. However, functional characterization of specific cell types during regeneration has been limited by the absence of validated cell-specific ablation systems. Here, we developed and rigorously compared two independent inducible genetic cell-ablation platforms — bacterial nitroreductase (NTR 2.0) and mammalian inducible caspase-9 (iCasp9), across developmental stages, animal sizes, and administration routes using various transgenic lines and grafting approaches. The NTR 2.0 platform showed limited applicability due to drug toxicity and solubility constraints, restricting its use primarily to larval stages via immersion. In contrast, the iCasp9 system demonstrated superior efficacy across all life stages, including large adults, with multiple viable administration routes. We further validated these platforms by systematically ablating CD68^+^ macrophages and examined functional consequences during tail regeneration. Sustained depletion revealed essential macrophage-dependent processes despite continuous macrophage repopulation from hematopoietic reservoirs: skeletal-element regeneration was completely abolished, spinal-cord axons degenerated without recovery, and neural crest-derived cells exhibited severe disorganization. These findings establish macrophages as critical orchestrators of central and peripheral nervous-system regeneration and skeletogenesis in axolotls, while providing validated tools for cell-type-specific functional studies across the axolotl lifespan.

## Introduction

Mammalian tissue injury typically culminates in fibrotic scarring and permanent functional deficit, a clinical challenge observed across organ systems from myocardial infarction to spinal-cord injury^1,2^. In stark contrast, salamanders including the Mexican axolotl (*Ambystoma mexicanum*) regenerate complex anatomical structures with complete functional restoration in the absence of scar formation^3, 4^. This remarkable regenerative capacity extends to clinically relevant tissues, including the central nervous system, cardiac muscle, lung, reproductive tissues, and entire limbs, positioning the axolotl as an invaluable model for discovering pro-regenerative mechanisms translatable to human medicine^5,6, 7,8,9,10^. Despite significant advances in axolotl genomics and the development of tissue-specific transgenic lines^11,12^, functional interrogation of cell-type-specific contributions to regeneration has been constrained by the absence of validated inducible ablation systems in axolotl.

We investigated whether two cell-ablation systems established in other model organisms could be effectively implemented in axolotl. In zebrafish, bacterial nitroreductase (NTR) systems have enabled cell ablation, though predominantly in larval stages due to limited efficacy in adults^13,14^. Compared to the first generation NTR system, the rationally engineered enhanced NTR 2.0 variant exhibits >100-fold increased sensitivity to the prodrug metronidazole (MTZ) or ronidazole (RON), which NTR converts to DNA-damaging free radicals to genetically induce cellular death^15,16^, thus enhancing ablation efficiency in larval zebrafish and feasibility in adult applications^17^. Comment. Although axolotls are thought to be resistant to oxidative stress^18, 19^, we anticipated that NTR 2.0, with its enhanced sensitivity, could potentially be an effective tool in mediating targeted cell ablation in axolotls.

In mammalian models, the iCasp9 system—comprising an FKBP12-F36V dimerization domain fused to caspase-9 lacking the caspase recruitment domain (ΔCARD-Casp9)—enables efficient conditional apoptosis of a range of cell types (e.g. T cells, iPSC-derived cells) upon administration of a synthetic homodimerizer ligand (AP20187/BB1070)^20,21,22^. This system has proven particularly effective for depleting difficult-to-ablate cell types such as macrophages in mice when iCasp9 is driven by colony-stimulating factor 1 receptor (CSF1R) regulatory elements^23,24^.

Here, we rigorously developed and compared both the NTR 2.0 and iCasp9 ablation platforms in axolotls across multiple dimensions: *in vitro* cytotoxicity in human, mouse, and axolotl cell lines; *in vivo* efficacy across larval, juvenile, and adult life stages; and optimization of drug-delivery routes. Our systematic cross-platform comparison revealed that while NTR 2.0 showed limited utility beyond larval immersion protocols, the iCasp9 system demonstrated robust efficacy across all life stages in axolotl via multiple administration routes. This included functional validation by allografting transgenically sensitized skin or blastemal tissue onto axolotls to demonstrate that the iCasp9 platform could efficiently ablate either tissue at various stages of limb regeneration in large adult animals, highlighting the effectiveness of ablating non-renewing cell types and representing a significant advance for studying adult regenerative processes in axolotl.

To test the effectiveness of these platforms in axolotl for ablating more challenging cellular targets, i.e., cells that undergo replacement after ablation, we developed several macrophage-specific transgenic ablation-reporter lines under control of the axolotl-specific CD68 pan macrophage promoter. Macrophages are established regulators of tissue repair across phylogeny^25,26^. We previously demonstrated the requirement for macrophages in axolotl limb and heart regeneration using clodronate liposomes, showing that systemic macrophage depletion results in regenerative failure and fibrotic tissue formation in both limb and heart^7,27^. These discoveries establish macrophages as master regulators of salamander regeneration, but their specific functions in nervous-system regeneration and skeletogenesis during tail regeneration remain undefined. Using our new macrophage-specific NTR 2.0 and iCasp9 ablation/reporter axolotl lines, we achieved sustained CD68^+^ macrophage depletion despite continuous hematopoietic repopulation, enabling us to discover that macrophages are required for: (1) regeneration and maintenance of spinal-cord axonal architecture, (2) organization of HNK1^+^ and GFAP^+^ neural crest-derived cells, and (3) *de novo* cartilage formation and skeletal outgrowth. These findings reveal previously unrecognized macrophage-dependent mechanisms controlling central and peripheral nervous-system regeneration and skeletogenesis in tail regeneration and establish validated ablation platforms that will enable the axolotl research community to perform rigorous cell-type-specific functional studies throughout development and regeneration.

## RESULTS

### Construction and *in vitro* evaluation of iCasp9 and NTR 2.0 ablation platform vectors

To enable pharmacologically controlled elimination of engineered cells, we generated ubiquitously expressed cassettes for Nitroreductase 2.0 (NTR 2.0)^16^ and humanized inducible caspase-9 (ihCasp9)^20^, each coupled to fluorescent reporters (CMV:NTR 2.0-P2A-tdTomato, CMV:ihCasp9-T2A-tdTomato). We assessed *in vitro* ablation efficiency in human (HEK293), mouse (L929), and axolotl (AL1 mesenchymal blastemal) cell lines. Cells were transiently transfected, cultured to peak expression, then exposed to dose gradients of the appropriate small molecule: metronidazole (MTZ) or ronidazole (RON) for NTR 2.0 and AP20187 (“B/B” homodimerizer) for ihCasp9 (**Fig. 1a).** Cell death modalities were quantified using a dual live-cell apoptosis staining approach: Annexin V (phosphatidylserine externalization, early apoptosis), caspase 3 activation (committed apoptosis), and dual labeling to capture progression or mixed modalities or Annexin V^+^ necrosis (**Fig. S1a, b**). Non-sensitized CMV:tdTomato controls were included to evaluate off-target drug toxicity at the highest concentrations tested.

**Figure 1.**
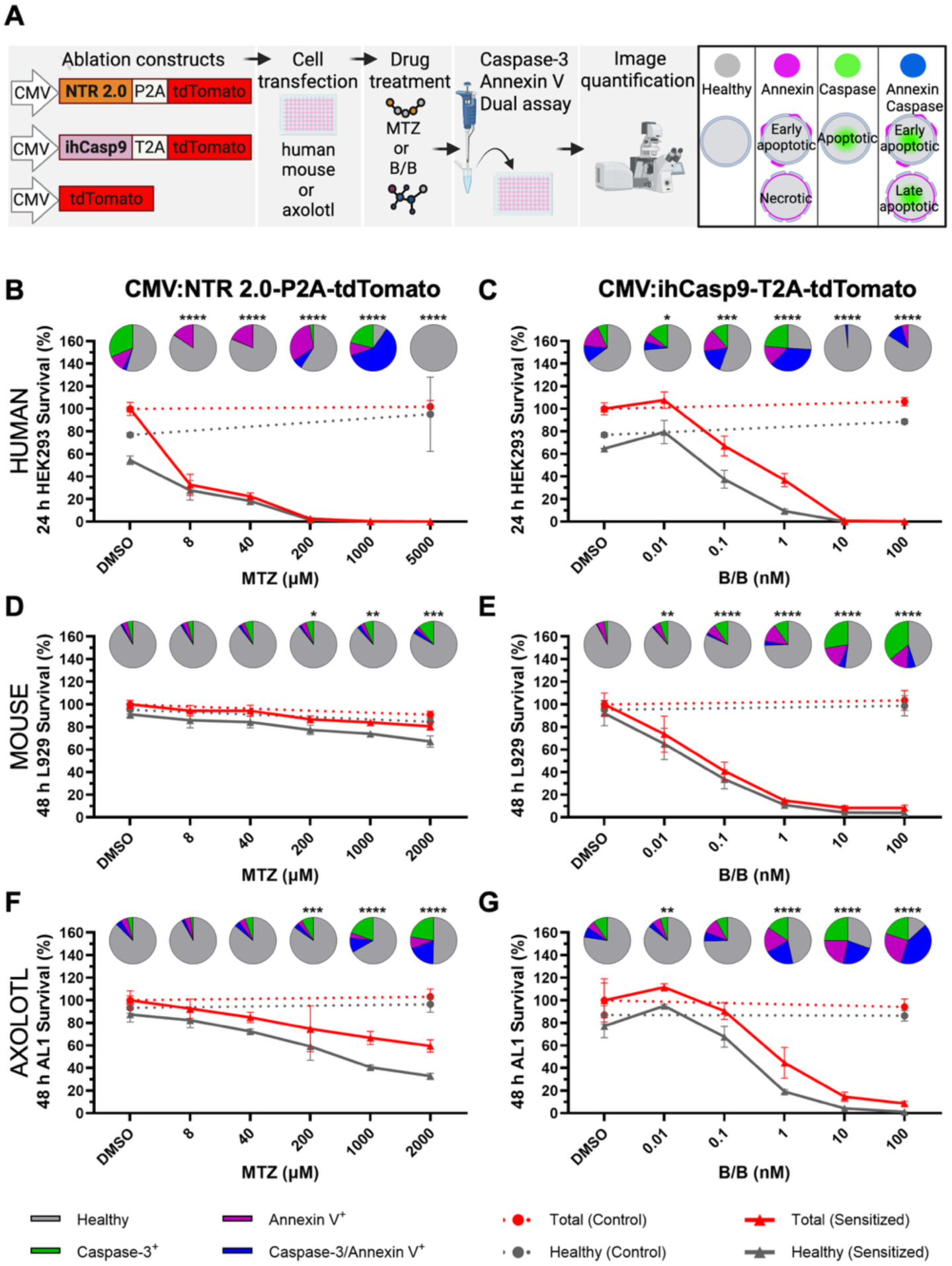
Cross-species comparison reveals differential ablation efficiency of CMV-driven NTR 2.0 and ihCasp9 systems *in vitro*. Established cell lines from human (HEK293; embryonic kidney), mouse (L929; dermal fibroblasts), and axolotl (AL1; limb fibroblasts) were transiently transfected with CMV:NTR 2.0-P2A-tdTomato (sensitized), CMV:ihCasp9-T2A-tdTomato (sensitized), or CMV:tdTomato (non-sensitized control) constructs and cultured to peak expression. Sensitized cells were exposed to dose gradients of metronidazole (MTZ; NTR 2.0 prodrug) or AP20187 (B/B; ihCasp9 homodimerizer). Non-sensitized tdTomato controls received maximum concentrations of both drugs to assess off-target toxicity. Cell death was quantified using live Caspase-3/Annexin V dual-apoptosis staining assay. **(A)** Experimental workflow schematic showing transfection timeline, drug exposure windows, and live apoptosis assay methodology. **(B-G)** Dose-response curves showing survival of sensitized (solid lines) versus control (dotted lines) cells relative to DMSO vehicle. Red lines: total transfected cell counts; gray lines: healthy transfected cell counts (Caspase-3/Annexin V⁻). Pie charts indicate cell fate distribution: healthy (gray), Caspase-3⁺ (green), Annexin V⁺ (magenta), or Caspase-3/Annexin V⁺ (blue). **(B)** NTR 2.0-sensitized HEK293 cells after 24 h MTZ exposure. **(C)** ihCasp9-sensitized HEK293 cells after 24 h B/B exposure. **(D)** NTR 2.0-sensitized L929 cells after 48 h MTZ exposure. **(E)** ihCasp9-sensitized L929 cells after 48 h B/B exposure. **(F)** NTR 2.0-sensitized AL1 cells after 48 h MTZ exposure. **(G)** ihCasp9-sensitized AL1 cells after 48 h B/B exposure. Error bars represent mean ± SD (N=3 wells per treatment). No statistically significant difference was observed between the DMSO vehicle and the maximum MTZ or B/B concentration in healthy non-sensitized tdTomato control cells across all species (unpaired t-test, significance threshold of *p*<0.05). Statistical significance of each inducible drug concentration relative to the DMSO vehicle was determined for healthy sensitized cells for all species by one-way ANOVA with Dunnett’s multiple comparisons test (\**p*<0.05, \*\**p*<0.01, \*\*\**p*<0.001, \*\*\*\**p*<0.0001). Representative of 3 independent experiments.

In human HEK293 cells, peak expression is detectable within 24 hours and NTR 2.0 showed excellent sensitivity to MTZ (∼49% at 8 μM and ∼97% at 200 μM) (**Fig. 1b, Fig S2a**). ihCasp9-sensitized cells are robustly ablated through subsequent 24 hours B/B exposure: ∼42% at 0.1 nM and ∼99% at 10 nM (**Fig. 1c**). Testing of two alternative expression constructs for ihCasp9 demonstrates that tdTomato-T2A-ihCasp9 is approximately 1.5 times more sensitive than tdTomato-IRES-ihCasp9 in HEK293 cells (**Fig. S3a, b**). Non-sensitized CMV:tdTomato controls showed no detectable off-target toxicity at 5000 μM MTZ or 100 nM B/B (**Fig. 1b,c**).

Mouse L929 fibroblasts, which are more challenging to transfect than HEK293 cells, reached peak expression at ∼48 hours post-transfection. NTR 2.0 demonstrated limited ablation under MTZ treatment: ∼20% at 2000 μM (18 h), rising modestly to ∼26% (48 h) (**Fig. 1d**, **Fig. S4a, c**). RON showed higher ablation efficiency than MTZ in L929 cells but still was still relatively inefficient compared to other cell lines tested (**Fig. S4b, d**). Concurrently under these conditions, caspase 3 labeling increased between 18 and 48 hours of prodrug exposure, indicating slower apoptotic progression in NTR 2.0-sensitized cells (**Fig. 1d, Fig. S4**). In contrast, ihCasp9 exhibited high ablation efficiency with B/B in both 18 and 48 hours of inducible drug exposure (**Fig. 1e, Fig. S5a, b**). At 18 hours, ablation efficiency was ∼66% at 0.1 nM and ∼93% at 10 nM, and cell-fate distribution was predominantly Annexin V^+^ across all concentrations, which is consistent with early activation of apoptosis (**Fig. S5a**). At 48 hours, ablation efficiency was ∼63% at 0.1 nM and ∼96% at 10 nM, with an evident shift towards increased caspase 3 labeling that is consistent with the progression to late apoptosis (**Fig. 1e, Fig. S5b**).

Axolotl AL1 (mesenchymal) cells derived from dermal limb blastema reached peak transgene expression at ∼96 hours post-transfection, with transfection efficiencies that are comparable to L929 cells (∼40%). NTR 2.0-sensitized AL1 cells showed low sensitivity to MTZ relative to HEK293 cells, but higher sensitivity than L929 cells: ∼35% at 2000 μM (18 hours), rising to ∼62% (48 hours) (**Fig. 1f**, **Fig. S6a, c**). Annexin V remained relatively stable across timepoints, whereas caspase 3 and dual labeling increased from 18 to 48 hours, indicating cumulative apoptotic engagement (**Fig. 1f**, **Fig. S6a, c**). Given reports that RON is more soluble than MTZ and outperformed MTZ in zebrafish NTR 1.0 systems (Lai et al., 2021)^28^, we evaluated RON against MTZ as an alternative prodrug for triggering NTR 2.0 cell death across species. In HEK293 cells, RON showed reduced sensitivity relative to MTZ at 8 μM, but similar efficiency at 40 μM and higher concentrations (**Fig. S2a, b**). In L929 cells, 2000 μM RON increased NTR 2.0 ablation ∼2 fold compared to MTZ at the same concentration for both 18-and 48-hour timepoints; caspase 3 labeling increased with time, mirroring MTZ kinetics (**Fig. S4a-d**). In AL1 cells, RON achieved ∼60% ablation at 2000 μM (18 hours), subtly increasing to ∼69% at the same concentration (48 hours), and RON outperformed MTZ ∼2 fold at 18 and 48 hours (**Fig. S6a-d**). Notably, Annexin V, caspase 3, and dual caspase 3/Annexin V populations in NTR 2.0 sensitized AL1 cells showed a slight increase with RON over time and with increasing concentrations in addition to the rise in caspase 3 and dual labeling in AL1 cells. This contrasts with the MTZ advantage over RON at low concentrations in HEK293 cells (**Fig. S2a, b**). The failure to ablate all NTR 2.0^+^ AL1 cells derived from a heterogenous population of mesenchymal limb progenitors may represent slower death kinetics or oxidative resistance in different blastemal cell types, an issue that could potentially be addressed by the growing list of alternative NTR prodrugs.

AL1 ihCasp9 ablation was highly effective in a dose-dependent manner with similar levels of B/B sensitivity at higher concentrations compared to L929 and HEK293 mammalian cell lines (**Fig. 1g**, **Fig. S7a, c**). At 48 hours, ihCasp9-sensitized AL1 cells achieved ∼30% ablation at 0.1 nM, ∼80% at 1nM and >95% at 10 nM (**Fig. 1g**, **Fig. S7c**). For comparison, at 48 hours L929 ihCasp9-sensitized cells achieved around 60% at 0.1 nM, ∼90% at 1 nM, and ∼95% at 10 nM (**Fig. 1e**, **Fig. S5b**). HEK293 ihCasp9-sensitized cells at 24hrs achieved around 60% at 0.1 nM, ∼95% at 1nM and >95% at 10 nM (**Fig. 1c**, **Fig. S3b**). At 18 hours, ihCasp9 ablation was not significantly different from controls between 0.1–1 nM B/B, but showed >90% ablation 10 nM B/B (**Fig. S7a**), which was similar to ihCasp9-sensitized L929 cells (**Fig. S5a**). Although AL1 cells required higher B/B doses for robust ablation, final kill rates of AL1 cells at 10 nM B/B were comparable to mammalian lines at all timepoints. For comparative purposes, we also constructed an axolotl-specific version of the iCasp9 module (iaxCasp9) (fusion of DmrA dimerization domain fused to truncated axolotl Caspase 9 lacking pro-CARD domain) and directly compared efficacy against ihCasp9. The iaxCasp9 modestly improved ablation sensitivity relative to ihCasp9 at 18 hours (**Fig. S7a, b**) but showed no detectable change at 48 hours (**Fig. S7c, d**).

Collectively, these data indicate that ihCasp9 delivers rapid, high-efficiency ablation across diverse cell types and species, at lower effective doses than are required for NTR 2.0-mediated ablation. Conversely, NTR 2.0 exhibits species-dependent ablation efficiency, achieving better efficacy in axolotl mesenchymal cells than in mouse fibroblasts. RON enhances NTR 2.0 ablation efficiency in both AL1 and L929 cell lines relative to MTZ, with acceleration of early apoptotic Annexin V activation restricted to AL1 cells. The time-resolved Annexin-V-to-caspase 3 shift is prominent in L929 cells and consistent with canonical apoptosis kinetics^20^. This process is more asynchronized in AL1 cells, potentially reflecting heterogeneous subpopulations and variable death trajectories.

### Effective *in vivo* cell ablation of mosaic CMV-driven transgenic NTR 2.0 or ihCasp9 genetically sensitized cells in larval limbs

The overarching goal for developing an inducible cell-ablation platform *in vivo* is to genetically sensitize a specific cell type and assess its function during developmental or regenerative processes. Encouraged by the potential of both systems, we generated transgenic axolotls driving ubiquitous expression of NTR 2.0 or ihCasp9 tagged with a fluorescent reporter under control of the CMV promoter to test these systems *in vivo (*CMV:NTR 2.0-P2A-tdTomato or CMV:ihCasp9-T2A-mCherry). Using the MTZ waterborne (WB) immersion protocol previously described for the NTR 2.0 system in zebrafish^16^ and the B/B homodimerizer intraperitoneal (*i.p.*) injection delivery method used in the mouse for iCasp9^29^, we sought to compare these systems in larval F0 transgenic founders. We selected animals with low levels of mosaicism predominantly localized to the appendages to mitigate the risk of ablating cells within vital organ systems (3- to 4-month-old, 3-4 cm in length). After 2 days of 10 mM MTZ exposure, ∼97% of the NTR 2.0^+^ cells were ablated relative to the untreated condition (**Fig. 2a, c**). The animal was imaged at 3 months post-treatment and the ablated cell population was not replaced. In the ihCasp9 animal, only ∼40% of the sensitized cells had ablated by 2 days following a single B/B injection via IP cavity (**Fig. 2b, d**). A maximum ablation efficiency of ∼90% was not observed until 6 days post-injection. This suggests that NTR 2.0 is more suitable for larval/juvenile studies than ihCasp9. These experiments revealed that both the NTR 2.0 and ihCasp9 systems can achieve sufficient levels of *in vivo* targeted cell ablation (**Fig. 2a-d)**.

**Figure 2.**
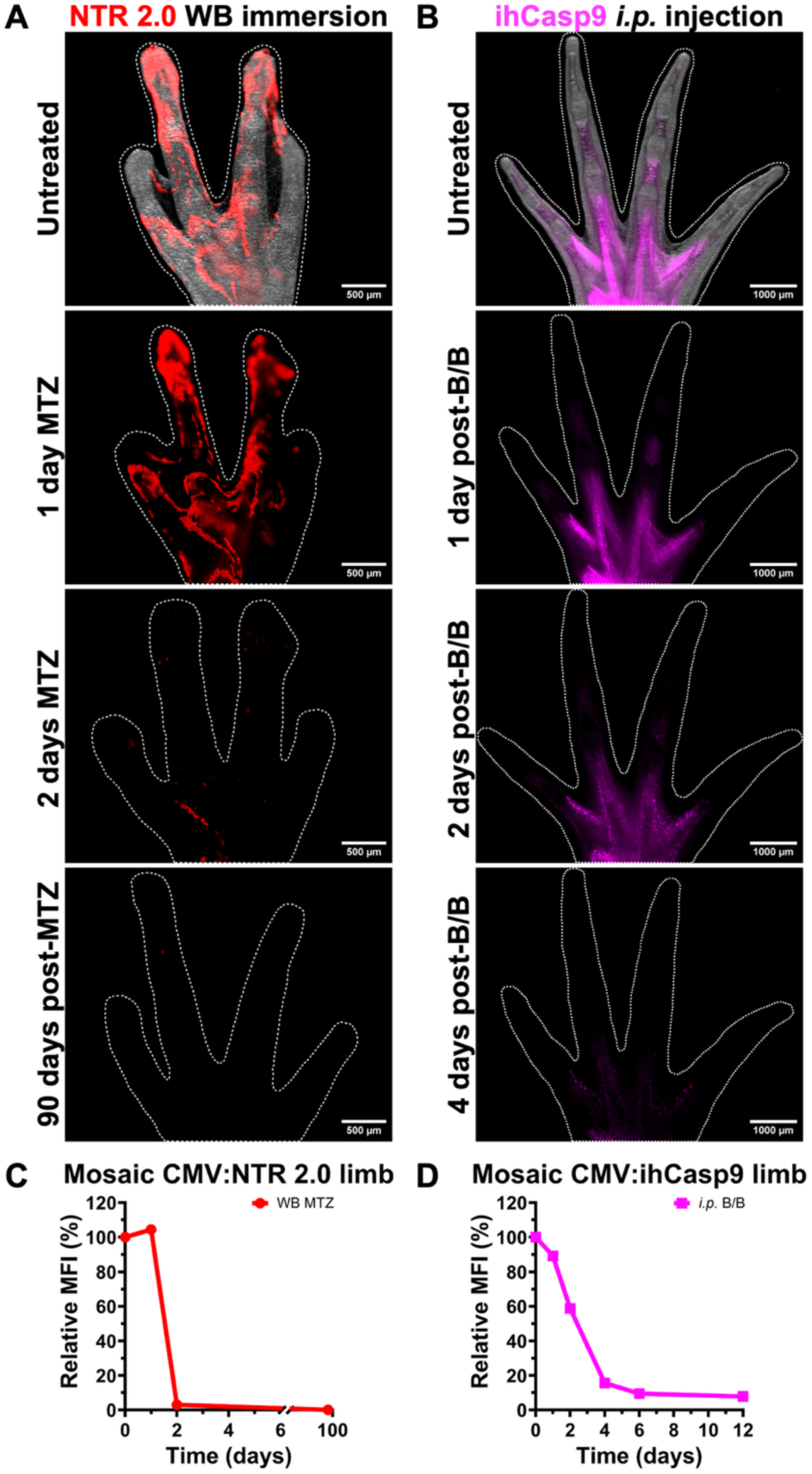
Mosaic CMV:NTR 2.0 and CMV:ihCasp9 transgenic juveniles demonstrate effective *in vivo* cell ablation. Mosaic (F0) transgenic juvenile axolotls expressing CMV:NTR 2.0-P2A-tdTomato (CMV:NTR 2.0) or CMV:ihCasp9-T2A-mCherry (CMV:ihCasp9) predominantly in appendages were subjected to either metronidazole (MTZ; NTR 2.0 prodrug) or AP20187 (B/B; ihCasp9 homodimerizer) drug treatments to assess *in vivo* ablation kinetics. Animals were selected for low-level expression to minimize off-target effects on critical organs. **(A)** Representative fluorescence images used to measure (C) relative mean fluorescence intensity (MFI) demonstrate efficient NTR 2.0 ablation in the limb after 2 days of continuous 10 mM MTZ waterborne (WB) immersion *in vivo* without repopulation following treatment (N=1 animal shown, representative of 2 independent experiments, 3-month-old, 3 cm length). Limb outline (white dotted line) was created at each timepoint to define the region for MFI quantification. Scale Bar: 500 µm. **(B)** Representative fluorescence images used to measure (D) relative MFI show ihCasp9 limb ablation efficiency within 2 to 4 days following a single 5 mg/kg B/B dose via intraperitoneal (*i.p.)* injection (N=1 animal shown, representative of 2 independent experiments, 4-month-old, 4 cm length). Limb outline (white dotted line) was created at each timepoint to define the region for MFI quantification. Scale Bar: 1000 µm. **(C)** Quantification of MFI in the limb of one representative mosaic CMV:NTR 2.0 juvenile axolotl relative to untreated control at pre-treatment (day 0), during treatment (1 and 2 days), and post-treatment (190 days) measured in Fiji (3-month-old, ∼3 cm length). Data are presented descriptively without statistical analysis. **(D)** Quantification of MFI in the limb of one mosaic CMV:ihCasp9 juvenile axolotl relative to untreated control at pre-treatment (day 0) and post-treatment (1, 2, 4, 6, and 12 days) measured in Fiji (4-month-old, 4 cm length). Data are presented descriptively without statistical analysis.

### *In vivo* ihCasp9 allograft ablation is highly effective through different administration routes where NTR 2.0 is more limited in adults

Cell-ablation tools have proven extremely valuable for studying development^30^, but our ultimate goal was to develop an effective ablation platform for the study of regeneration in large adult axolotls. To test the effectiveness of these systems in adult animals, we employed a transgenically encoded skin-allografting approach and evaluated different routes of drug delivery in both systems for optimal efficacy (**Fig. 3a**). Tissue grafting is an established and frequently utilized technique in axolotls for tracking the migration and contribution of labelled cells to regenerating structures. This is facilitated by the absence of acute rejection in axolotls, whereby grafts can survive up to 60 days before chronic rejection^31^.

**Figure 3.**
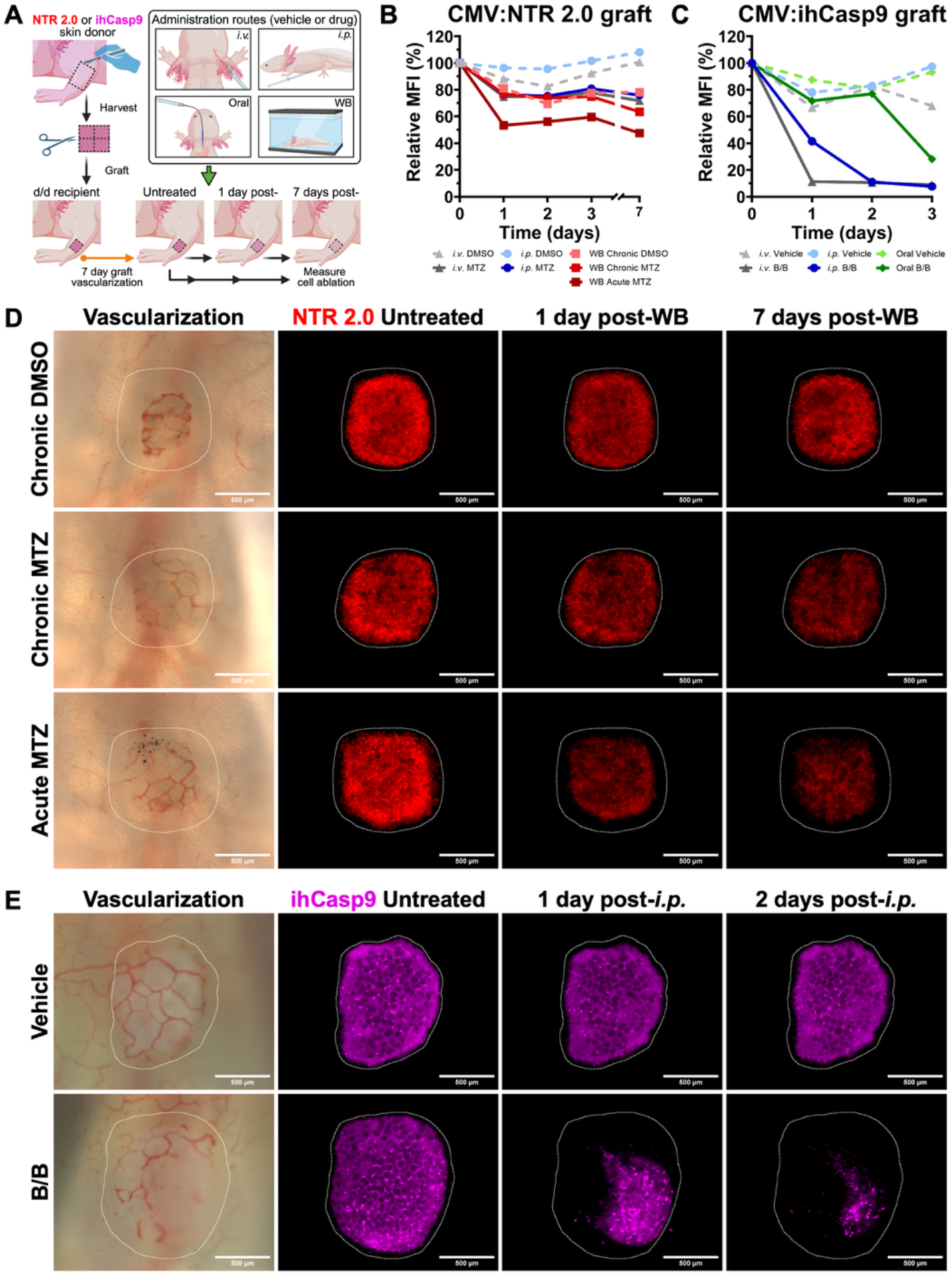
Allograft ablation model reveals route-dependent drug efficacy with superior performance of systemically delivered B/B. Full-thickness transgenic skin grafts from CMV:NTR 2.0-P2A-tdTomato (CMV:NTR 2.0) or CMV:ihCasp9-T2A-mCherry (CMV:ihCasp9) donors were transplanted onto the limbs of non-transgenic d/d recipients and allowed to vascularize for 7 days prior to metronidazole (MTZ; NTR 2.0 prodrug) or AP20187 (B/B; ihCasp9 homodimerizer) drug delivery. Evaluated administration routes include intravenous injection (*i.v.*), intraperitoneal injection (*i.p.*), oral gavage, and waterborne immersion (WB). **(A)** Experimental schematic depicting grafting timeline, drug administration routes, and imaging schedule. **(B)** Quantification of mean fluorescence intensity (MFI) in NTR 2.0 skin grafts treated with DMSO vehicle (dotted lines) or MTZ (solid lines) relative to untreated control via *i.v.* (845 mg/kg), *i.p.* (845 mg/kg), chronic WB (1 mM for 7 days), and acute WB (20 mM for 18 hours) (N=1 animal per treatment, 8- to 10-month-old, ∼10 cm length). Measured pre-treatment (day 0) and post-treatment (1, 2, 3, and 7 days) MFI in Fiji. Data are presented descriptively without statistical analysis. **(C)** Quantification of MFI in ihCasp9 skin grafts treated with vehicle (dotted lines) or 10 mg/kg B/B (solid lines) through *i.v.*, *i.p.*, and oral routes of administration (N=1 animal per treatment, 8- to 10-month-old, ∼10 cm length). Measured pre-treatment (day 0) and post-treatment (1, 2, and 3 days) MFI in Fiji. Data are presented descriptively without statistical analysis. **(D)** Representative extended depth of focus projections show skin allograft vascularization for CMV:NTR 2.0 expression (red) prior to treatment (day 0) and ablation efficiency post-treatment (1 and 7 days) with chronic (1 mM for 7 days) or Acute (20 mM for 18 hours) MTZ WB immersion relative to the DMSO control (N=1 animal per treatment, representative of 3 independent experiments, 8- to 10- month-old, ∼10 cm length). Transgenic skin allograft boundary (white dotted line) was created at pre-treatment (day 0) then applied to all timepoints as the region for MFI quantification. Scale Bar: 500 µm. **(E)** Representative extended depth of focus projections show skin allograft vascularization for CMV:ihCasp9 expression (magenta) prior to treatment (day 0) and ablation efficiency post-treatment (1 and 2 days) with a single 10 mg/kg B/B *i.p.* injection relative to the vehicle (N=1 animal per treatment, representative of 3 independent experiments, 8- to 10-month-old, ∼10 cm length). Transgenic skin allograft boundary (white dotted line) was created at pre-treatment (day 0) then applied to all timepoints as the region for MFI quantification. Scale Bar: 500 µm.

To test the effectiveness of both ablation tools in adult animals, full-thickness skin was harvested from either CMV:NTR 2.0-P2A-tdTomato or CMV:ihCasp9-T2A-mCherry adult donors and divided into at least 10 individual 0.8-1.0 mm squares (N=1 donor animal per transgenic line, 10- to 12-month-old, ∼14 cm length). Donor skin grafts were subsequently transplanted onto at least 10 non-transgenic leucistic (d/d) recipients within a corresponding size-matched region of excised skin while maintaining proximal-distal identity (8- to 10-month-old, ∼10 cm length). After 7 days, successful allograft integration was confirmed by restoration of blood flow (Data in supplemental movie – (**Fig. S8)**. Drug efficacy was then evaluated by supplying a single dose through different routes of administration (**Fig. 3a**). Because intravenous (*i.v.)* injection is more technically challenging in the axolotl than it is in the mouse, we evaluated the potential for *i.p.* injection as a less burdensome and higher-throughput route of administration. To provide additional non-invasive routes of administration suitable for large animal cohorts, we refined and tested the oral gavage method, which is rarely used in salamander research.

The NTR 2.0 system was established in zebrafish, where MTZ is supplied through the immersion protocol^16^. We first performed toxicity testing for chronic MTZ exposure in animals that were 2 cm, 4 cm and 7 cm in length (**Fig. S9)**. These experiments showed less visible toxicity in larger animals than smaller animals at the same concentrations but established 1 mM as a safe dose for chronic exposure in all sizes. In larger animals, an acute dose of 20 mM is tolerated for only a brief period, i.e., ∼18 h, but is not tolerated in smaller animals. To test the efficacy of waterborne (WB) immersion dosing in this model, we performed a chronic 1 mM MTZ WB immersion spanning a 7-day period or a single 20 mM acute MTZ WB immersion for 18 hours, using our transgenic skin-grafting model **(Fig. 3b, d, Fig. S10).** Under chronic MTZ exposure conditions, 10 cm animals showed no real ablation above the levels we observed in DMSO treated control animals **(Fig. 3b, d, Fig. S10).** In contrast, acute MTZ exposure in 10 cm animals resulted in ∼43% cell-ablation efficiency beginning immediately following dosing and continuing at a relatively constant fluorescence level over the 7-day period **(Fig. 3b, d, Fig. S10)**.

Attempts to test alternative routes of MTZ administration are complicated by the poor solubility of this compound in aqueous solutions at ambient temperature and neutral pH (around 5 mg/ml). MTZ requires high DMSO percentages to keep the drug solubilized for *in vivo* injections or oral gavage administration. In our experiments, delivery of MTZ via oral gavage was not tolerated at DMSO percentages required for this system. Delivery of extremely high MTZ concentrations (845 mg/kg is predicted *in vivo*; the theoretical maximum molarity is 4.9 mM) via *i.p.* and *i.v.* injections showed poor ablation efficiency **(Fig. 3b, Fig. S11)** even though we used doses double that of the effective dose *in vitro*. Furthermore, following MTZ administration, these animals were lethargic and anemic with signs of acute toxicity, limiting the utility of these delivery modalities. In an attempt to reduce the amount of DMSO and improve the solubility of MTZ, we tested the inclusion of 10% niacinamide (a form of vitamin B3), previously reported to improve MTZ solubility^32^. Niacinamide dramatically improved solubility, and although it did seem to enhance the ablation efficiency of both MTZ (**Fig. S12a, a’**) and an alternative prodrug nifurpirinol (NFP) (also activated by the NTR enzyme) (**Fig. S12b, b’),** we were concerned that niacinamide could cause non-specific vehicle toxicity in the graft, and thus we did not pursue it further. In the absence of niacinamide, NFP lost efficacy in all routes of administration at 6 mg/kg (*in vivo* theoretical maximum molarity is 24.4 μM) (**Fig. S12c,c’)**. We then investigated RON^28^ as a more soluble NTR prodrug. Like MTZ, RON was well tolerated at 1 mM chronic exposure, but like MTZ, it was found to be toxic at 2 mM and 5 mM in animals ranging from 2 to 7 cm in length (**Fig. S9)**.

We then tested the toxicity vs. efficacy of MTZ, RON, and the B/B homodimerizer drug in a reciprocal grafting model where the fluorescence of either CMV:NTR 2.0-P2A-tdTomato or CMV:ihCasp9-T2A-mCherry grafts were exposed to each of the three drugs (**Fig. S13)**. These experiments showed that exposure of ihCasp9 grafts to either MTZ or RON showed no toxicity, and that NTR 2.0 graft exposure to the B/B homodimer was also shown to be non-toxic (**Fig. S13)**. Intraperitoneal injection of 10 mg/kg B/B into axolotls exhibited dramatic cell ablation in ihCasp9-sensitized grafts (**Fig. S13)**. Exposure to 20 mM acute MTZ showed some ablation of NTR 2.0-sensitized grafts, and chronic 1 mM RON exposure was more effective than chronic 1 mM MTZ exposure (**Fig. S13)**.

Evaluation of the ihCasp9 system in our transgenic grafting model revealed high ablation efficiency via all routes of B/B administration (*i.v.*, *i.p.*, oral gavage) (**Fig. 3c, e, Fig. S14**). At 10 mg/kg B/B (a predicted maximum *in vivo* molarity of 6.47 μM, which is 1,000 times the effective B/B concentration *in vitro*), *i.v.* injection produced rapid ablation of ∼90% of the graft within 1 day of administration (**Fig. 3c, Fig. S14**). We found that ablation following *i.p.* delivery of B/B was slightly slower, with ∼90% ablation by 2 days post-injection (**Fig. 3c, e, Fig. S14**). After 3 days, ∼70% ablation was observed with the oral gavage route of administration (**Fig. 3c, Fig. S14**). Due to the high cost of dosing via immersion, this method was not practical.

### Effective *in vivo* ablation of ihCasp9-sensitized blastema grafts in large animals

To test the effectiveness of the ihCasp9 platform in mediating *in vivo* cell ablation of heterogeneous cell types participating in regeneration in large animals, we took advantage of the blastemal grafting model. In this model, immediately following axolotl limb amputation, the wound is sealed by rapidly migrating epithelial cells and the underlying tissue undergoes a regenerative response that produces a mound of undifferentiated cells known as the blastema, which ultimately gives rise to all limb structures distal to the amputation plane. We harvested 7-day post-amputation (DPA) blastemal tissue from CMV:ihCasp9-T2A-mCherry genetically sensitized animals and then grafted the tissue onto the freshly amputated forelimbs of d/d control recipients (8- to 10-month-old, ∼10 cm length) (**Fig. 4a**). We found that in the absence of blastema ablation, these grafts are the main source of the regenerated limb (**Fig. 4b-h**). However, when animals are treated with B/B at three distinctly characterized regenerative stages (late bud, paddle, and digit stage), full ablation is observed, with regression of the regenerated tissue (**Fig. 4b-h**). We also performed grafting of 13 DPA blastemal tissue from CMV:ihCasp9-T2A-mCherry animals onto CAGGS:GFP transgenic recipients to show that effective ablation within 2 days of a single 10 mg/kg B/B injection (**Fig. S15a-c**). This highlights the potential of the ihCasp9 platform to efficiently ablate many different cell types involved in the regenerative process at defined stages of regeneration and in animals of different sizes.

**Figure 4.**
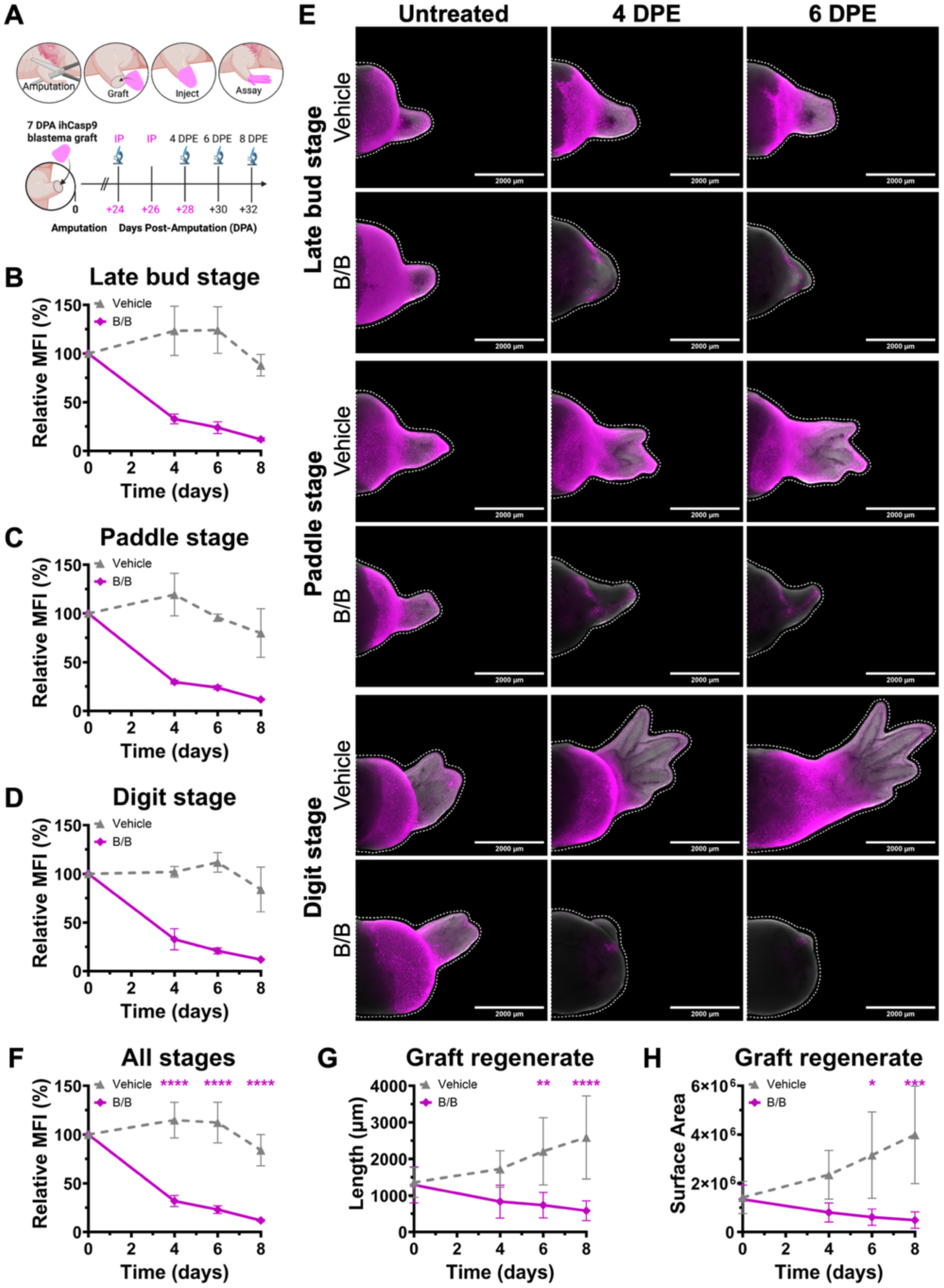
Stage-specific blastema ablation in adult axolotls demonstrates the versatility of the ihCasp9 system. CMV:ihCasp9-T2A-mCherry (CMV:ihCasp9) blastemal tissue was harvested at 7 days post-amputation (DPA) and grafted onto the stumps of freshly amputated d/d recipient limbs. Several intraperitoneal injections (*i.p.*) of vehicle or 10 mg/kg AP20187 (B/B; ihCasp9 homodimerizer) were administered to assess ablation efficacy at defined stages of limb regeneration. **(A)** Schematic of experimental design depicts blastema graft placement and integration following amputation of d/d recipient limbs. Timeline includes *i.p.* delivery of vehicle or B/B (magenta), MFI measurement timepoints (microscope icon), and days post-exposure (DPE) from the initial injection. **(B-D)** Quantification of mean fluorescence intensity (MFI) in CMV:ihCasp9 grafts at three distinct regenerative stages: **(B)** late bud, **(C)** paddle, and **(D)** digit (N=4 animals per stage, N=2 animals per treatment, 8- to 10-month-old, ∼10 cm length). Measured pre-treatment (day 0) and post-treatment (4, 6, and 8 days) MFI in Fiji. Error bars represent mean ± SD. Data are presented descriptively without statistical analysis. **(E)** Representative extended depth of focus projections comparing vehicle-versus B/B-treated CMV:ihCasp9 grafts prior to treatment and days after the first *i.p.* injection was administered within each regenerative stage (N=4 animals per stage, N=2 animals per treatment, representative of 3 independent experiments, 8- to 10-month-old, ∼10 cm length). Graft boundary (white dotted line) was used as the region for MFI quantification. Scale Bar: 2000 µm. **(F-H)** Longitudinal analysis of CMV:ihCasp9 graft morphometrics: **(F)** MFI, **(G)** regenerate length, **(H)** regenerate surface area across all stages of limb regeneration measured in Fiji (N=6 animals per treatment group, 8- to 10-month-old, ∼10 cm length). Error bars represent mean ± SD. CMV:ihCasp9 grafts treated with B/B showed a significant decrease in relative MFI and regression of regenerate outgrowth relative to the vehicle (one-way ANOVA with Tukey’s multiple comparisons test, \**p*<0.05, \*\**p*<0.01, \*\*\**p*<0.001, \*\*\*\**p*<0.0001).

### Development of axolotl macrophage-specific labeling and ablation tools

To test the utility of each system in cell types known to be difficult to ablate, we developed new macrophage-specific transgenic reporter/ablation axolotls for each system using the CD68 pan-macrophage promoter. Using existing single-cell datasets, we identified several genes that are expressed in the monocyte/macrophage cell linages with high specificity (**Fig. 5a**). *Mpeg1* is commonly used as a macrophage-specific marker in zebrafish but is also expressed in B cells and NK cells in zebrafish^33^ ^34^. In the axolotl, we found that *Mpeg1* has several paralogs with expression in macrophage and non-macrophage cell types, and we therefore excluded *Mpeg1* from our ablation tool development (**Fig. 5a**). Instead, we focused on *CD68*, which shows high expression levels and macrophage specificity in the axolotl, similar to what has been reported in *CD68* mouse transgenic lines^35^ (**Fig. 5a**). We found that *CSF1R* was highly restricted to monocyte/macrophages, but due to the complex long-range enhancer elements required for expression in all mononuclear phagocytes, we focused on isolation of the simpler *CD68* promoter (**Fig. 5a**). In mice, a ∼2.9kb human *CD68* promoter can direct GFP expression specifically to both monocytes or macrophages^35^. The murine *CD68* promoter region contains several regulatory elements located between − 7.0 and − 2.5 kb from the transcriptional start site, and these regulatory elements have strong enhancer activity in macrophages and repressor activity in non-myeloid cells^35, 36^. The proximal -150-bp sequence of the mouse *CD68* promoter exhibits high-level promoter activity in macrophages^36^. Based on these lines of evidence in mice, we predicted that the axolotl *CD68* proximal promoter could be used to drive macrophage-specific expression. After isolation of ∼ 3.7kb of the proximal promoter region of the axolotl *CD68* gene, we cloned this 3.7-kb segment upstream of both the NTR 2.0 and ihCasp9 ablation and reporter coding sequences in separate plasmids and produced transgenic animals. Larval animals were confirmed to express fluorescent reporters in both tissue-resident and circulating immune cells as early as 8 days post-fertilization (data not shown). We raised animals to ∼10-15 cm in length before performing limb amputations and flow cytometric analysis on dissociated limb blastemas harvested at 4 DPA. We were able to clearly identify CD68:NTR 2.0-P2A-tdTomato^+^ cells in transgenic animals **(Fig. 5b)**. These cells exhibited monocyte/macrophage morphology after cytospin analysis (**Fig. 5c**) and were identified as monocyte/macrophages via previously validated flow cytometric surface-marker staining (IB4/CD18 staining)^37,38,39^ (**Fig. S16**). CD68^+^ and CD68^-^ FACS-sorted cells were harvested for RNA and qRT-PCR analysis (**Fig. 5d**). This analysis confirmed CD68 and CSF1R enrichment in tdTomato^+^ cells and enrichment of a B-cell marker (IGHM)^40^ ^41^ and a neutrophil marker (CSF3R)^42^ in tdTomato^-^ cells, confirming specificity of the CD68 promoter (**Fig. 5d**). Immunostaining of limb blastemas at 7 DPA shows co-expression of the CD68^+^ cell population with the canonical macrophage marker Iba1 (**Fig. 5e**, **Fig. S17a, b**) Both staining and flow cytometry indicated that CD68:tdTomato expression appeared weaker in monocytes than mature macrophages **(Fig. 5e**). It was also noted that macrophages were also closely associated with acetylated tubulin^+^ axons in the regenerating limb (**Fig. S17a, b**).

**Figure 5.**
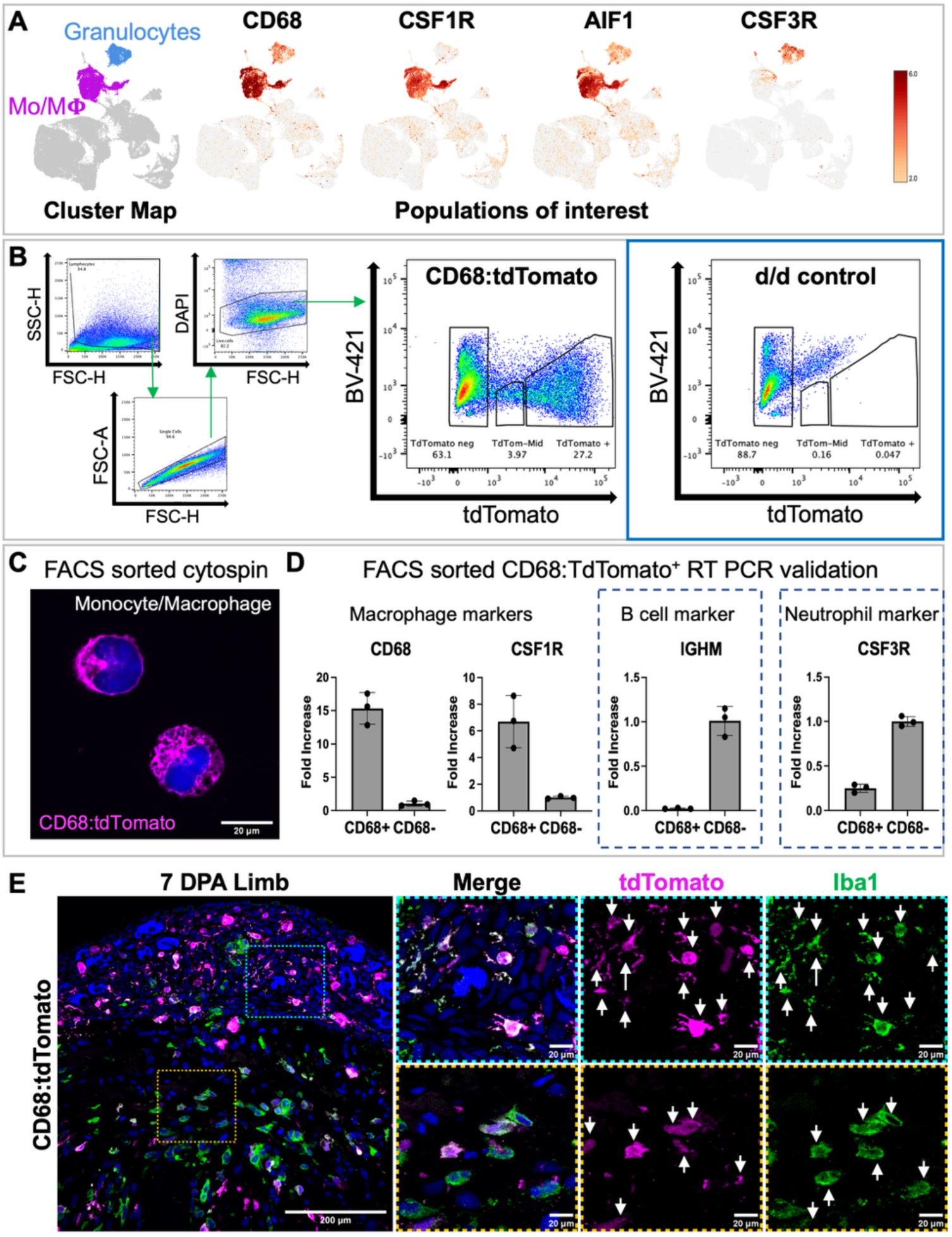
CD68 promoter-driven transgenic reporters specifically label monocyte/macrophage populations in axolotls. Validation of CD68:NTR 2.0-P2A-tdTomato transgenic line as a macrophage-specific reporter through multi-modal characterization combining transcriptomics, flow cytometry, cell morphology, and immunofluorescence. **(A)** Single-cell RNA sequencing analysis of regenerating limb tissue across a temporal time course showing cell-type-specific expression patterns. Heatmap demonstrates enrichment of CD68, CSF1R, and AIF1 in monocyte/macrophage clusters and CSF3R in granulocyte/neutrophil populations. UMAP showing clustering of 59128 cells over 8 timepoints. Li et al. dataset.**(B)** Flow cytometry gating strategy for isolation of CD68:tdTomato⁺ cells from 4 DPA regenerating limbs. tdTomato+ cells were around 30% of the dissociated total population. **(C)** Representative image of FACS-sorted CD68:tdTomato⁺ cytospin preparations stained with DAPI demonstrates characteristic monocyte/macrophage morphology (magenta) including kidney-shaped nuclei (blue) (N=3 independent sorts). Scale Bar: 20 µm. **(D)** Quantitative RT-PCR analysis of FACS-sorted CD68:tdTomato⁺ versus tdTomato⁻ populations. Bar graphs show enrichment of macrophage markers (CD68, CSF1R) in tdTomato⁺ cells and enrichment of B-cell (IGHM) and neutrophil (CSF3R) markers in tdTomato⁻ fractions. Cells were isolated by FACS from three transgenic animals and pooled per condition prior to RNA extraction to obtain sufficient material (N=3 animals, ∼1-year-old, ∼12 cm length). Each sample was analyzed in triplicate technical qPCR reactions; dots represent individual technical replicates and bars represent the mean +/- SD. Gene expression is shown as fold change, calculated using the ΔΔCt method with normalization to RBL27 and Beta-actin reference genes. Data are shown from a representative experiment; the experiment was repeated independently with similar results. **(E)** Confocal maximum intensity projections of 7 DPA CD68:tdTomato transgenic limb section shows coexpression of tdTomato (Magenta) with the canonical macrophage marker Iba1(AIF1; in green), shown with white arrows. Merged image with DAPI (blue) confirms nuclear localization. Tissue-resident macrophages found in the wound epithelium (cyan inset) and circulating monocytes recruited to the blastema (orange inset) are both shown to be CD68/Iba1^+^. Scale Bar: 200 µm. Inset Scale Bar: 20 µm. DPA=Days post-amputation. Antibodies: Rabbit anti-RFP (1:200, Rockland cat. 600-401-379); Chicken anti-Iba1 (1:50, Antibodies Inc. cat. IBA1-0020).

### Macrophage ablation reveals essential roles in cartilage and spinal-cord repair during larval tail regeneration

To demonstrate the capacity of these systems to specifically ablate macrophages, we used the tail regeneration model to evaluate the effect of macrophages in larval tail regeneration. Macrophages have been shown to be essential for adult limb and heart regeneration in salamander regeneration^7, 27^ and larval zebrafish regeneration^43,44^. To determine if axolotl tail regeneration is a macrophage-dependent process in larval axolotls, we used germline F1 CD68:NTR 2.0-P2A-tdTomato and CD68:mScarlet-IRES-ihCasp9 transgenic lines with corresponding non-sensitized d/d control animals (2-month-old, 2-3 cm in length). Tails were imaged prior to injury then amputated ∼0.5 mm from the distal portion of the notochord. We captured the time course of larval tail regeneration at 2, 7, 14, and 21 DPA in animals treated with MTZ/RON (NTR 2.0 prodrug) (**Fig. 6a**) or the B/B (ihCasp9 homodimerizer) (**Fig. 6b**). We compared *in vivo* ablation as well as regenerative outcomes by measuring the regenerate cartilage length, spinal-cord length, and surface area at previously described timepoints (**Fig. 6c-h**). The ihCasp9 system (**Fig. 6e, f, h-j, Fig. S18-20**) was far more effective than the NTR 2.0 system at inhibiting regeneration, and RON was found to be more effective than MTZ under chronic exposure (**Fig. 6c, d, g, Fig. S21-S23).** It should also be noted that overnight applications of 2 mM RON acute immersion every 3 days from 0 to 9 DPA (a total of four overnight applications) enhanced macrophage recruitment and regeneration relative to the chronic 1 mM RON exposure from 0 to 14 DPA (**Fig. S22**), highlighting the challenges in ablating self-renewing populations. Importantly, while B/B administered to d/d control (non-genetically-sensitized) animals had no effect on regeneration (**Fig. 6h, Fig. S20**), nor did chronic exposure to 1 mM MTZ or RON from 0 to 14 DPA, exposure of d/d animals to acute overnight applications of 10 mM MTZ WB immersion every 3 days from 0 to 9 DPA (total of four applications) inhibited tail regeneration (**Fig. 6g, Fig. S21-23**), highlighting the toxicity risks associated with the NTR 2.0 system.

**Figure 6.**
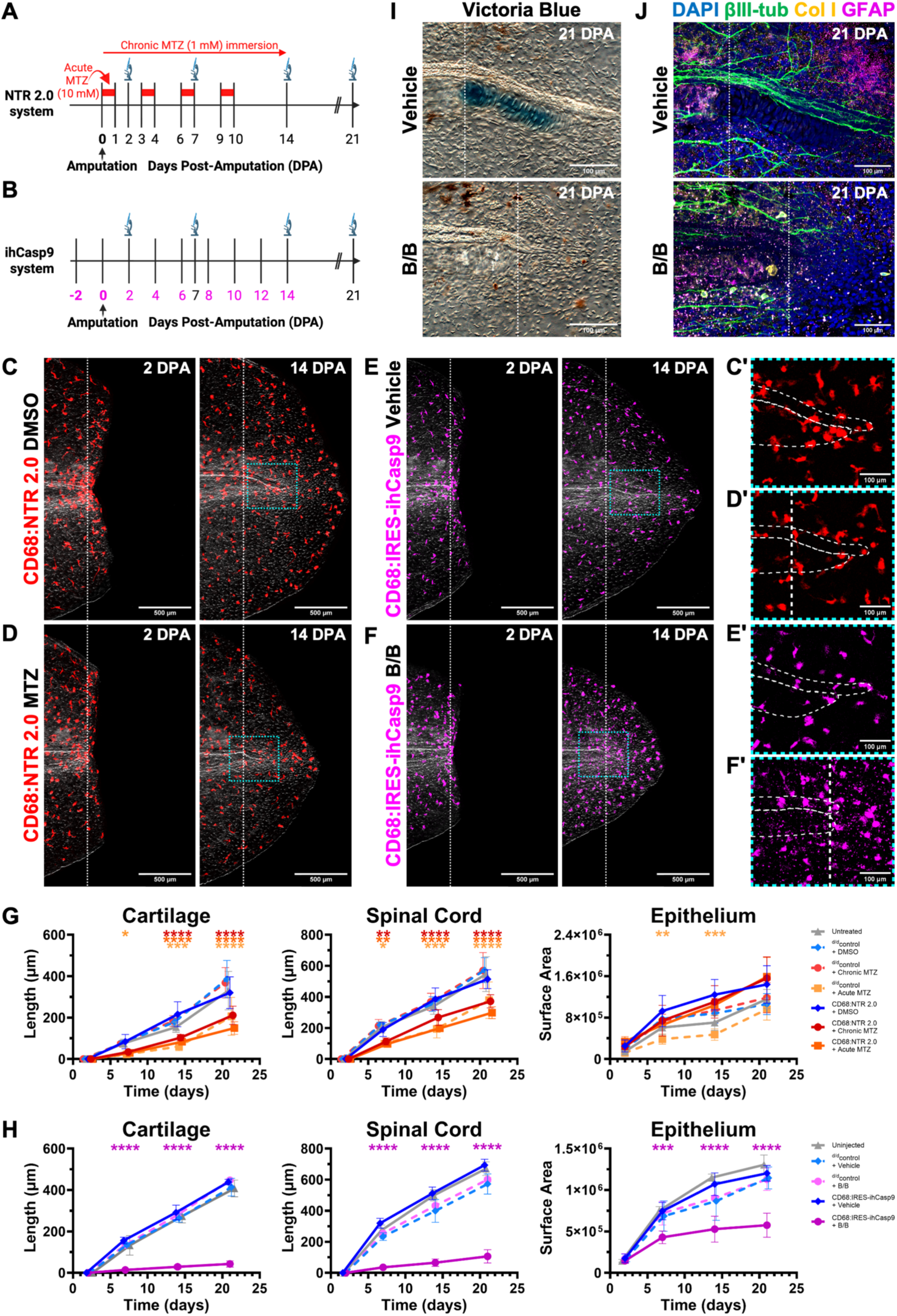
Sustained macrophage depletion via NTR 2.0 or ihCasp9 systems reveals essential roles in tail regeneration. Tail amputations were performed on germline F1 CD68:NTR 2.0-P2A-tdTomato (CD68:NTR 2.0) or CD68:mScarlet-IRES-ihCasp9 (CD68:IRES-ihCasp9) transgenic larvae alongside age and size matched wildtype d/d then subjected to repeated inducible drug treatments to maintain macrophage depletion throughout the regenerative process. **(A)** Experimental timeline of NTR 2.0 macrophage ablation and non-sensitized d/d toxicity evaluation shows chronic (14 day – long red arrow) versus four acute (18 hours every 3 days - red solid blocks) doses of MTZ administered through waterborne immersion (WB) with imaging schedule. Imaging timepoints shown with microscope icon and time relative to amputation. **(B)** Experimental timeline of ihCasp9 macrophage ablation and non-sensitized d/d toxicity evaluation shows 10 mg/kg B/B intraperitoneal injection (*i.p.*) regimen (every 2 days for 23 days) with imagining schedule. Timepoints with *i.p.* drug delivery of B/B shown in magenta. Imaging timepoints shown with microscope icon and time relative to amputation. **(C, D)** Representative extended depth of focus projections show CD68:NTR 2.0 transgenic tails at 2 and 14 DPA: **(C)** 1% DMSO vehicle control, **(D)** chronic 1 mM MTZ-treated (N=5 animals per treatment, 2-month-old, 2-3 cm length). Amputation plane indicated by white dotted line and cyan boxed regions mark the inset location. **(C’, D’)** Insets show decreased CD68:NTR 2.0 recruitment and rounded macrophage morphology within the regenerative zone when treated with MTZ relative to DMSO control. Thin white dotted lines outline the spinal cord (dorsal) and cartilage (ventral). Scale Bar: 500 µm. Inset Scale Bar: 100 µm. **(E, F)** Representative extended depth of focus projections show CD68:IRES-ihCasp9 transgenic tails at 2 and 14 DPA: **(E)** vehicle control, **(F)** 10 mg/kg B/B-treated (N=3-5 animals per treatment, 2-month-old, 2-3 cm length). Amputation plane indicated by white dotted line and cyan boxed regions mark the inset location. **(E’, F’)** Insets show fragmented CD68:IRES-ihCasp9 debris with complete inhibition of both cartilage and spinal cord regeneration relative to the vehicle control. Thin white dotted lines outline the spinal cord (dorsal) and cartilage (ventral). Scale Bar: 500 µm. Inset Scale Bar: 100 µm. **(G)** Quantitative morphometric analysis of tail regenerates in CD68:NTR 2.0 transgenics (solid lines) and non-sensitized d/d controls (dashed lines) treated with MTZ. Measurements of cartilage length, spinal cord length, and epithelial surface area that extend beyond the amputation plane were collected at 2, 7, 14, and 21 DPA in Fiji (N=5 animals per treatment, 2-month-old, 2-3 cm length). **(H)** Quantitative morphometric analysis of tail regenerates in CD68:IRES-ihCasp9 transgenics (solid lines) and non-sensitized d/d controls (dashed lines) shows cartilage length, spinal cord length, and epithelial surface area relative to amputation plane measured at 2, 7, 14, and 21 DPA in Fiji (N=3-5 animals per treatment, 2-month-old, 2-3 cm length). **(I)** Representative maximum intensity projections of Victoria blue stained CD68:IRES-ihCasp9 transgenic tails at 21 DPA demonstrate regenerative failure following amputation of the cartilaginous rod in B/B-treated animals compared to the vehicle-treated controls (N=3 animals per treatment, 2-month-old, 2-3 cm length). Amputation plane indicated by white dotted line. Scale Bar: 100 µm. **(J)** Representative confocal maximum intensity projections of vehicle-versus B/B-treated CD68:IRES-ihCasp9 transgenic tails stained with DAPI (blue), βIII-tubulin^+^ axons (green), GFAP⁺ radial glial cells (magenta), and collagen I (yellow) at 21 DPA (N=3 animals per treatment, 2-month-old, 2-3 cm length). Amputation plane indicated by white dotted line. Scale Bar: 100 µm. Error bars represent mean ± SD. Statistical significance of inducible drug treatments relative to the respective vehicle controls at each regenerative timepoint was determined by two-way ANOVA with Tukey’s multiple comparisons test (\**p*<0.05, \*\**p*<0.01, \*\*\**p*<0.001, \*\*\*\**p*<0.0001). MTZ=Metronidazole (NTR 2.0 prodrug), B/B=AP20187 (ihCasp9 homodimerizer), DPA=Days post-amputation.

Through sustained macrophage depletion using either CD68:NTR 2.0-P2A-tdTomato or CD68:mScarlet-IRES-ihCasp9 transgenic animals, we discovered that, despite continuous hematopoietic repopulation, both platforms inhibited regeneration of spinal-cord and cartilage components after amputation (**Fig. 6c-j**). However, the ihCasp9 system showed more dramatic and complete inhibition (**Fig. 6e, f, h-j).** Tail amputations in germline F1 CD68:mScarlet-IRES-ihCasp9 transgenic axolotls, followed by repeated B/B *i.p.* injections every 2 days from -2 to 21 DPA revealed four critical macrophage-dependent processes **(Fig. 6 e, f, h-j, Fig. 7a,b**): **(1) loss of distal extension of the ependymal tube; (2) complete abolishment of skeletal-element regeneration**, with failure to form new cartilage after tail amputation; **(3) spinal-cord axon degeneration without recovery**, demonstrating that macrophages are essential for both regeneration and maintenance of CNS axonal architecture; and **(4) severe disorganization of HNK1^+^ neural crest-derived cells**, which fail to repopulate the peripheral nervous system after amputation^45^ ^46^. Of note, in axolotl tail regeneration, the ependymal tube containing the CNS leads the notochord tip (cartilage) in outgrowth. Staining revealed only a partial overlap between CNS-associated GFAP^+^ (glial fibrillary acidic protein) cells^47^ and neural crest-associated HNK1^+^ cells (**Fig. S24**) dorsal to the ependymal tube. HNK1^+^ cells were profoundly disrupted by macrophage depletion **(Fig. 7a, b, Fig. S25e**). Whole-mount staining suggests that, compared to control animals, macrophage-depleted animals do not show any excessive collagen deposition, but exhibit decreased numbers of GFAP^+^ cells inside the ependymal tube with no obvious differences to the variable GFAP+ populations outside the ependymal tube (**Fig. S25a-d**). We reason that GFAP^+^ cells inside the neural tube may be required for axonal growth, while those outside the neural tube may disrupt normal ependymal tube extension. These findings establish macrophages as critical orchestrators of central and peripheral nervous system regeneration and skeletogenesis during axolotl tail regeneration, extending previous observations of macrophage requirements in salamander limb and heart regeneration^7, 27^.

**Figure 7.**
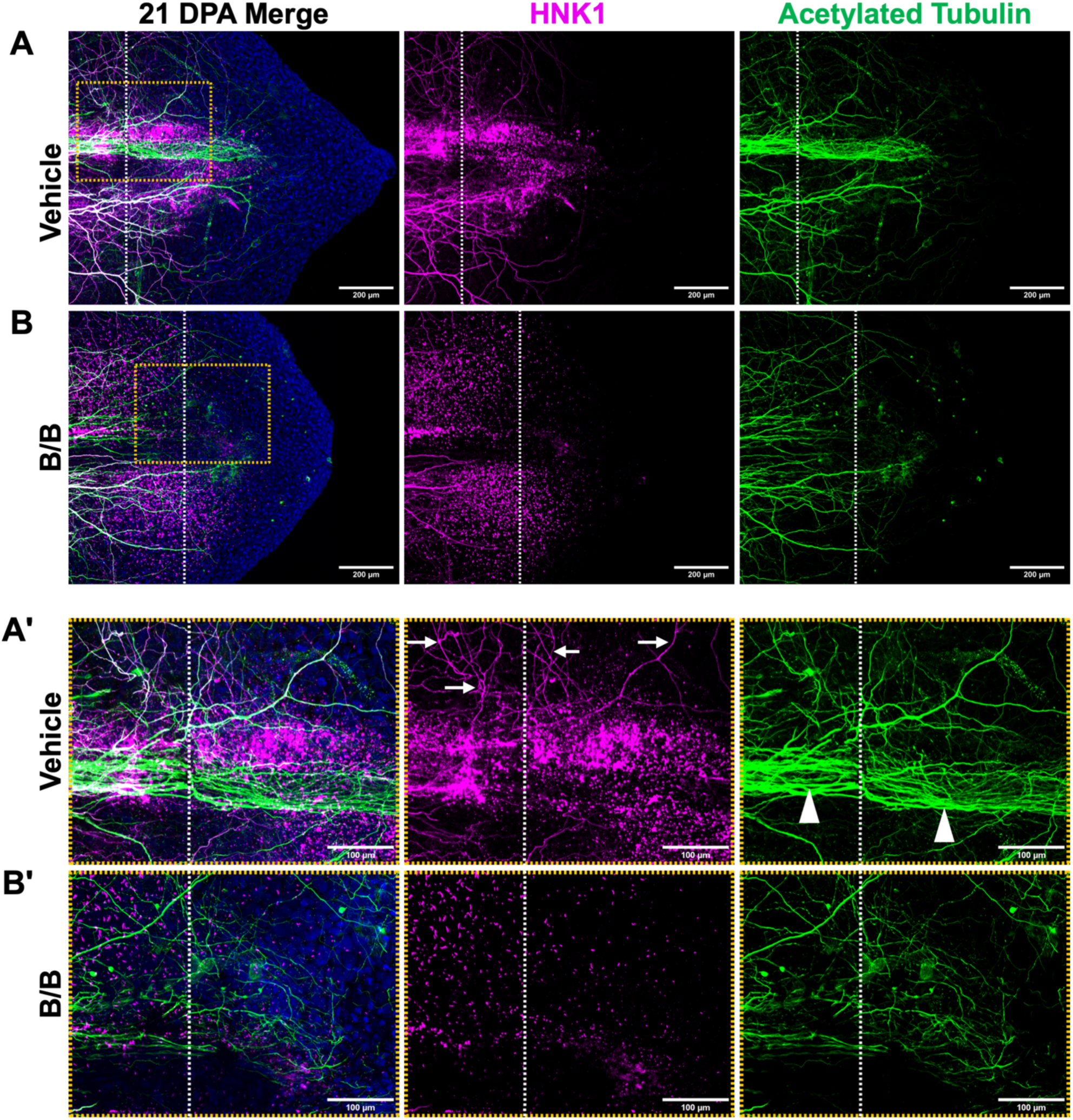
Macrophage depletion causes axonal degeneration, displacement of HNK1⁺ neural crest-derived cells, and failure to generate myelinated axons in peripheral nervous system during tail regeneration. Whole-mount immunofluorescence analysis of CD68:mScarlet-IRES-ihCasp9 transgenic tails at 21 days post-amputation reveals profound disruption of nervous system regeneration following sustained macrophage ablation with 10 mg/kg AP20187 (B/B; ihCasp9 homodimerizer) administered via intraperitoneal injection. **(A, B)** Representative confocal maximum intensity projections of whole-mounted CD68:mScarlet-IRES-ihCasp9 tails at 21 DPA (vehicle-versus B/B-treated) stained for acetylated tubulin⁺ axons (green), HNK1⁺ neural crest-derived Schwann cells (magenta), and DAPI nuclear counterstain (blue)(N=3 animals per treatment, 2-month-old, 2-3 cm length). Amputation plane indicated by white dotted line. Cyan boxed regions indicate the dorsal inset location while orange boxed regions indicate the midline inset location. Scale Bar: 200 µm **(A’, B’)** High-magnification insets (cyan boxes) of dorsal regions at the amputation plane. Normal regenerative axon outgrowth beyond amputation plane shown with white arrowhead (acetylated tubulin:green) absent in Mϕ-depleted (B/B-treated tails). Normal HNK1+ (Magenta) neural-crest-derived Schwann cell myelinated axons in peripheral nerves present in control (white arrows) missing in Mϕ-depleted (B/B-treated tails) showing displacement of HNK1+ cells away from the midline. Scale Bar: 100 µm. **(A’’, B’’)** High-magnification insets (orange boxes) of midline regions show spinal cord axonal architecture. Large hollow arrowhead highlights axonal degeneration proximal to amputation plane in B/B-treated animals **(B’’)**, contrasting with robust axonal extension beyond the amputation plane in vehicle controls **(A’’)**. Scale Bar: 100 µm. CD68:tdTomato^+^ cells stained with anti-tdTomato antibody (magenta). Macrophages stained with anti-Iba1/AIF1 (green). Axons stained with Acetylated tubulin (yellow).

Morphometric analysis of fluorescently tagged CD68^+^ macrophages confirms that the ablation of wound macrophages is dependent on drug access. We quantified the morphological changes in CD68^+^ macrophages in three locations (rostral zone adjacent to amputation, caudal zone beyond the amputation plane, and in a remote peripheral location from amputation site) at three timepoints during the regeneration process (2, 7, and 14 DPA). In the periphery, morphological rounding of ihCasp9-sensitized macrophages is limited (**Fig. S26a, d**), but is prevalent in NTR 2.0-sensitized macrophages (**Fig. S26b, c, e**). Due to B/B delivery via *i.p.* injection, the ihCasp9 system displays an inside-to-outside mode of cell ablation that relies on drug delivery via the lymphatic and vascular network (**Fig. S26a, d**). In contrast, NTR 2.0 prodrugs are toxic when injected, but can be delivered to thin tissues via absorption from the tank water, dictating an outside-to-inside mode of delivery that fails to ablate internal organs or deep structures and limits its use to very small animals or peripheral cellular compartments in larger animals (**Fig. S26b, c, e**). As indicated in **Fig. S26**, CD68:NTR 2.0 animals and CD68:ihCasp9 animals both show labelling of tissue-resident macrophage populations and patrolling monocyte populations recruited to wounds. The NTR 2.0 system induces morphological changes in macrophages in a gradient, with less ablation of macrophages in internal tissues and more ablation of those closer to the skin, where drug concentrations are higher using immersion drug delivery (**Fig. S26b, c, e**). In contrast, the *i.p* route of administration of B/B used with the ihCasp9 system predominantly ablates monocytes and tissue-resident macrophages that are close to endothelial vessels carrying the drug (**Fig. S26a,d**). We observed evidence of mScarlet^+^ puncta (apoptotic macrophage debris) in B/B-treated ihCasp9^+^ macrophages starting at 7 DPA (**Fig. S18**). However, puncta were not observed in NTR 2.0^+^ macrophages with MTZ or RON at the timepoints we analyzed (**Fig. S21-22**).

The efficiency of the ihCasp9 system can be further enhanced with T2A-peptide-optimized expression. Our aforementioned macrophage-depletion studies in tail regeneration used germline CD68:mScarlet-IRES-ihCasp9 transgenic animals. However, it is known that the second transgene under control of the IRES element typically has 20-50% of the expression level of the first gene, depending on cell type^48,49^. To improve the sensitivity of ihCasp9 in macrophages, we created a CD68:ihCasp9-T2A-tdTomato line that will maximize ablation efficiency by producing a 1:1 molecular ratio between tdTomato and ihCasp9. Because we saw less B/B-mediated ablation in cells more distant from vessels, we wanted to know if use of the T2A peptide linker arrangement^50^ would improve ablation in tissues with less vascularization. We generated transgenic animals with the CD68:iCasp9-T2A-tdTomato plasmid and raised them as F0 mosaic animals to the larval stage (2-month-old, 2-3 cm length). To confirm labeling of the monocyte population, we exclusively removed epithelial tissue from the distal portion of the tail (termed epithelial priming) and screened the animals at 2 days post-epithelial removal (**Fig. S27a, c, d**). Animals that showed CD68:ihCasp9-T2A-tdTomato^+^ recruitment to the injured epithelium were selected to repeat the tail amputation assay previously described (**Fig. 6b**) with a shortened B/B *i.p.* injection regimen occurring every 2-3 days from -2 to 13 DPA (**Fig. S27a**). These experiments revealed dramatic inhibition of tail regeneration (**Fig. S27b-d**) and enhanced ablation of the tissue resident population with the T2A peptide linker arrangement relative to the previous IRES configuration. This featured extensive puncta formation (fluorescent cell fragments) across the amputated tissue despite the expected mosaicism, with a predicted maximum of 50% of macrophages being genetically sensitized/labelled (**Fig. S27b-f**). These new ablation technologies will not only facilitate the investigation of macrophage-specific contributions in various developmental and regenerative contexts but can be modified to enable the characterization of other cell types at any developmental stage in axolotl.

## DISCUSSION

### Comparative evaluation of ablation platforms: NTR 2.0 limitations and ihCasp9 advantages in adult axolotls

Our systematic comparison reveals critical differences between the NTR 2.0 and ihCasp9 ablation platforms that have important implications for regeneration research. While NTR 2.0 represents a substantial improvement over first-generation NTR systems, exhibiting ∼100-fold enhanced MTZ sensitivity in zebrafish^16^, it remains fundamentally limited by prodrug toxicity and delivery constraints in large animals. Previous studies in zebrafish documented that first-generation NTR systems require confoundingly toxic MTZ concentrations (10 mM) for effective ablation^15^, with some cell types remaining resistant^51^. The enhanced NTR 2.0 variant achieves ablation at 0.1-1 mM MTZ in zebrafish larvae and can ablate previously resistant cell types^16^. However, use of NTR 2.0 in adult zebrafish typically requires 2.5 mM MTZ for 3 days to achieve skin-cell ablation, or 5 mM for 2 days for β-cell ablation, a concentration that is nearly toxic to the organism over that exposure period, and adult applications remain predominantly limited to superficial tissues^51,17^. Concordantly, our findings demonstrate that in adult axolotls, even the use of NTR 2.0 presents challenges. The poor aqueous solubility of MTZ necessitates high DMSO concentrations incompatible with systemic delivery. In addition, effective MTZ immersion protocols (20 mM acute exposure) for adult axolotls nearly reach toxicity thresholds (**Fig. S9**). Further, injectable formulations of MTZ at 845 mg/kg produced off-target systemic effects in adult axolotls, including lethargy and anemia. While NTR is effective in embryonic and larval zebrafish via water-based delivery, its efficacy decreases in larger, more complex animals due to limited tissue penetration and the "outside-in" mode of action when delivered by immersion. Alternative prodrugs to MTZ such as RON or NFP have shown increased efficacy and safety in zebrafish^28,52^. While RON did show enhanced ablation relative to MTZ in the axolotl, NFP did not. NFP was previously found to have off target effects by inhibiting axon regeneration at 4 µM in animals with no outward signs of toxicity^28^. The fundamental limitation with NTR 2.0 is that MTZ and related prodrugs cannot be delivered systemically at concentrations required for deep-tissue ablation in large animals without causing toxicity.

In contrast, the ihCasp9 system overcomes these limitations through several key advantages (1) the B/B homodimerizer drug (AP20187/AP1903) is soluble and non-toxic at effective concentrations; (2) systemic delivery (*i.v.*, *i.p.*, oral gavage) enables "inside-out" ablation via vascular networks; (3) effective concentrations (1-10 nM *in vitro*, 10 mg/kg *in vivo*) are orders of magnitude below toxic thresholds; (4) rapid kinetics allow ablation within 24-72 hours; and (5) efficacy is maintained across all developmental stages, from larvae to large adults. The ihCasp9 system has been extensively validated in mammalian T-cell therapies, where a single dose of a dimerizer drug induces apoptosis in >99% of transduced cells^20, 21^.

Our grafting and blastema-ablation experiments demonstrate that ihCasp9 effectively ablates non-renewing tissues (skin grafts) and rapidly-renewing populations (blastema cells, macrophages) in ∼10 cm adult animals—applications that have proven challenging or impossible with NTR systems. The new transgenic CD68:ihCasp9 macrophage-ablation axolotl lines we developed demonstrate an efficiency from larval to adult stages that was previously unattainable. Macrophages are notoriously difficult to deplete pharmacologically, and genetic-ablation systems often fail to maintain depletion against continuous hematopoietic repopulation^23,24^. Our sustained macrophage depletion protocol (repeated B/B dosing via *i.p.* injection every 2-3 days) effectively overcomes this challenge, enabling discovery of macrophage-dependent regenerative processes that would be masked by incomplete or transient depletion.

These validated ablation technologies will enable the axolotl research community to perform rigorous cell-type-specific functional studies throughout development and regeneration^53^. This will ultimately advance our understanding of the mechanisms underlying scarless tissue repair^3^.

### Macrophage control of axon regeneration: integration with mammalian and teleost models

Our finding that macrophages are essential for spinal-cord axon regeneration in axolotls aligns with emerging evidence across vertebrate models but also reveals important context-dependent nuances. In mammals, macrophages exhibit paradoxical roles: they can promote axon regeneration through inflammatory modulation and debris clearance, yet can also cause neurotoxicity and axonal retraction^1,54,55,56^. Studies of rodent spinal-cord injury demonstrate that classically activated (M1) macrophages are neurotoxic and cause axonal dieback, whereas alternatively activated (M2) macrophages promote regenerative growth without toxicity^57^. Activated macrophages in rodent model organisms enhance dorsal root ganglion axon regeneration in the injured spinal cord, but can simultaneously induce extensive axonal retraction through direct physical interactions^55, 58^ and control the severity of the glial scar^59^. Our macrophage-depleted axolotls exhibited persistent axon degeneration without recovery, suggesting that in regeneration-competent species, macrophages maintain a predominantly pro- regenerative phenotype that is absolutely required for CNS repair. Our results in macrophage- depleted axolotls show a lack of ependymal tube extension beyond the amputation plane and a loss of distal GFAP^+^ cell migration inside the neural tube associated with a lack of axonal growth. GFAP^+^ cells outside the neural tube are found distal to the amputation plane independent of functional macrophages. Further work is required to assess whether these cells share any similarities with reactive (scar-inducing) glia in mammals. However, in contrast to regenerative failure with excessive scarring in axolotl hearts and limbs lacking macrophages^7,27^, we did not observe any excessive collagen deposition in failed tail regeneration lacking macrophages.

In zebrafish, peripheral macrophages—but not neutrophils or microglia—are necessary for spinal-cord axon regeneration^60^. Macrophage-deficient *irf8* mutant zebrafish show that prolonged inflammation with elevated TNF-α and IL-1β fail to regenerate spinal axons. Critically, while TNF-α promotes regeneration, IL-1β must be dynamically controlled by macrophages to enable functional tissue repair in zebrafish^60^. This parallels our observation that axolotl macrophage depletion prevents axon regeneration, although the specific cytokine profiles remain to be determined. The absolute requirement for macrophages in both zebrafish and axolotl CNS regeneration contrasts sharply with mammals, where macrophages often exacerbate injury, suggesting that an evolutionary divergence in macrophage polarization states underlies regenerative capacity.

Recent work in a rat model demonstrates that regenerative macrophages accumulate in dorsal root ganglia following “conditioning injury” to peripheral nerves and are essential for enhancing central axon regeneration^61^. Self-renewing macrophage populations in dorsal root ganglia contribute to nerve regeneration^62^, and M2 macrophage-based scaffolds promote spinal-cord repair by modulating inflammation and supporting axonal growth^63, 64^. Our findings extend these observations to axolotls, demonstrating that in highly regenerative vertebrates, macrophages not only support spinal axon regeneration, but are also absolutely required for successful axonal extension.

### HNK1^+^ neural crest cells and GFAP^+^ cells: overlapping populations in tail regeneration

Our observation that macrophage depletion causes disorganization of HNK1^+^ neural crest-derived cells, coupled with partial overlap between GFAP^+^ and HNK1^+^ populations, suggests complex interactions between glial and neural-crest lineages during regeneration. HNK1 (CD57, B3GAT1) is a well-established marker for migratory neural-crest cells that labels subpopulations involved in formation of peripheral nervous system structures, including Schwann cells, sensory neurons, and enteric ganglia^46,65^. During axolotl development, HNK1^+^ cells migrate along defined pathways and contribute to diverse derivatives including cartilage, bone, smooth muscle, and peripheral glia ^66,67^. In axolotls, GFAP is primarily expressed in CNS astrocytes, but is also found in non-myelinating Schwann cells, enteric glia, and neural stem/progenitor cells^47^. Importantly, GFAP^+^ progenitors possess broader potential than classically appreciated: in mice they contribute to vascular smooth muscle cells in both neural crest- and non-neural crest-derived vessels^68^, and GFAP^+^ radial glia in the zebrafish spinal cord give rise to multiple neural lineages^69,70,71^. The overlap we observe between GFAP^+^ and HNK1^+^ cells in regenerating axolotl tails likely represents neural crest-derived boundary cap cells, radial glia-like progenitors, or Schwann-cell precursors that express both markers transiently during migration and differentiation.

The disorganization of HNK1^+^ cells that we observed following macrophage depletion suggests that macrophages provide an essential function for neural-crest-cell migration and organization, such as the necessary guidance cues, trophic support, or extracellular-matrix remodeling. In mammalian peripheral nerve injury, macrophages regulate Schwann-cell responses and axonal guidance. Our findings indicate that in regenerating axolotl tails, macrophages orchestrate the spatial organization of neural crest-derived cells critical for peripheral nervous system reconstitution, linking immune regulation to neural patterning.

### Macrophage regulation of ependymal tube-coordinated tail regeneration in axolotl larvae

In urodele tail regeneration, the ependymal tube serves as the primary organizing structure that spatially and temporally coordinates regeneration of all axial tissues. Following tail amputation in axolotls, the ependymal tube extends well beyond the amputation plane in the first week post-amputation through enhanced proliferation of ependymal cells by reducing cell cycle length^19,72^, establishing the distal-most extent of the regenerate before significant outgrowth of the notochord or axons occurs. Our finding that macrophage ablation inhibits ependymal tube extension and dramatically reduces GFAP^+^ radial glial cells within the neural tube positions macrophages as essential upstream regulators of this pioneering ependymal response. This macrophage-dependent establishment of the ependymal tube is critical, as it provides both a) structural scaffolding through radial glial processes that guide regenerating axons—a function directly compromised by macrophage depletion as evidenced by reduced axonal extension in our ablation experiments—and b) inductive signals through floor plate-derived Sonic hedgehog (Shh) signaling that patterns the surrounding blastema tissue and drives ventral cartilage formation^73,74,75^.

The regenerated axolotl tail cartilage arises through two distinct cellular mechanisms that depend on proper ependymal tube formation: first, individual ependymal glial cells migrate out of the regenerating spinal cord into surrounding blastema tissue and transdifferentiate into cartilage and muscle cells, representing ectoderm-to-mesoderm lineage switching^76^; second, and quantitatively predominant, cartilage forms from mesodermal blastema cells that respond to Shh signaling secreted by the ventral floor plate of the ependymal tube, as demonstrated by the complete prevention of Sox9 expression and cartilage formation following cyclopamine-mediated Hedgehog inhibition^75^. Our observation that macrophage ablation results in failure to initiate new cartilage formation reveals that macrophages are essential for both mechanisms of cartilage generation: the structural defect in ependymal tube extension resulting from macrophage ablation would prevent both the emigration of transdifferentiating ependymal cells and the establishment of the Shh-secreting floor-plate domain required to induce cartilage from blastema cells. This dual-origin mechanism contrasts with axolotl limb regeneration, where lineage tracing revealed that dermis can produce cartilage and tendons, but that cartilage cells rarely cross between mesodermal tissue types, suggesting that tail regeneration exhibits greater lineage promiscuity than limb regeneration^77^. The finding that a single upstream immune-cell type—macrophages— are required for establishment of the ependymal organizing center that subsequently coordinates multiple downstream regenerative programs (radial glia scaffolding, axon guidance, dual-mechanism cartilage formation) demonstrates that immune-neural interactions represent a critical regulatory node controlling urodele regenerative capacity. The coexistence of both lineage-restricted (Shh-responsive blastema cells) and lineage-switching (transdifferentiating ependymal cells) mechanisms within a single regenerative context reveals that cellular plasticity during urodele tail regeneration exists along a continuum rather than as a binary property, and our data suggest that macrophage-mediated establishment of the ependymal organizing center may be a prerequisite for accessing the full spectrum of this regenerative plasticity.

The complete absence of cartilage regeneration following macrophage depletion in our axolotl tail regeneration model represents a previously unrecognized macrophage function in vertebrate skeletogenesis. In mammals, macrophages regulate bone remodeling and fracture healing, but their role in cartilage formation during development or regeneration is less defined. Our finding that axolotl macrophages are absolutely required for *de novo* cartilage formation suggests they may secrete factors that induce or maintain chondrogenic progenitors, regulate matrix deposition, or control the differentiation state of cartilage precursors—potential mechanisms that merit further investigation.

### Conclusions and Future Directions

We have established two cell-ablation platforms in axolotls, i.e., the NTR 2.0 system and the ihCasp9 system, and demonstrated that the ihCasp9 system provides superior efficacy, flexibility, and safety for adult-stage applications, while the NTR 2.0 system remains functionally limited by prodrug delivery constraints despite its engineering improvements. Using validated CD68:ihCasp9 transgenic axolotls, we discovered that macrophages are master regulators of multiple regenerative processes: they are absolutely required for spinal-cord axon regeneration, organization of neural-crest-derived peripheral nervous system elements, and *de novo* cartilage formation. These findings position macrophages at the nexus of neural, skeletal, and immune-system interactions during regeneration.

The mechanistic basis for macrophage control of these diverse processes remains to be elucidated. Future work should focus on identifying the specific macrophage-derived signals (cytokines, growth factors, extracellular matrix components) that regulate neural and skeletal progenitors, determining whether distinct macrophage subpopulations control different aspects of regeneration, and investigating whether the spatial relationship between macrophages and regenerating tissues (particularly the leading apical ampulla) is essential for coordinated outgrowth. The tools we have validated here—particularly the CD68:ihCasp9 system, which we demonstrated to be functional across life stages—will enable these mechanistic investigations in adult axolotls, bringing regenerative medicine research closer to understanding how highly regenerative vertebrates achieve scarless, functional tissue replacement.

## Methods

These studies are reported in accordance with ARRIVE guidelines.

### Ethics Statement

All scientific procedures involving animals were undertaken in line with Animal Ethics Committee guidelines for MDI Biological laboratory IACUC.

### Animal maintenance

Ambystoma mexicanum (Mexican axolotl) animals (d/d leucistic “white hosts” originally obtained from the Ambystoma Genetic Stock Centre (AGSC), Lexington, KY, United States, and captive bred. *Ambystoma mexicanum* (Mexican axolotl) were individually housed in 20% Holtfreter’s solution on a 12-hour light, 12-hour dark cycle at 18⁰C. larval/juvenile axolotls less than 4 cm were fed artemia while animals larger than this consumed commercial trout pellets (Rangan). Prior to animal surgeries (blood collection, amputation, and live imaging), animals were anaesthetized using 0.1% ethyl-3-aminobenzoate methanesulfonate salt (MS-222; Sigma-Aldrich) until there was no response to external stimulus. Animals were imaged on a 150 x 15mm petridish coated with 1% agarose and submerged in a low level of 20% Holtfreters solution covered with a damp Kimwipe.

### Drug Solubilization

Metronidazole (MTZ) (Millipore Sigma cat. M3761) and ronidazole (RON) (Cayman Chemical cat. 33071) prodrugs were primarily solubilized in dimethyl sulfoxide (DMSO) (Sigma-Aldrich cat. D8418) then placed in the Eppendorf Thermomixer R at 40⁰C, 1400rpm for 5 minutes. Nifurpirinol (NFP) (MedChemExpress cat. HY-135470) was solubilized in DMSO to make a 20 mg/ml (81.4 mM) stock solution stored in 10 ul aliquots at -80⁰C. For waterborne immersion, MTZ and RON were diluted to their respective dosages in 20% Holtfreter’s solution with a 1% final DMSO concentration. For *i.v.* and *i.p.* injection, MTZ was cooled to RT then directly administered.

An alternative method for solubilizing prodrugs was tested using 10% niacinamide (Sigma-Aldrich cat. N06360100G) and 5% DMSO diluted in 0.1 M sodium chloride (Sigma-Aldrich cat. S5886) made in Cell and Molecular grade UltraPure Sterile Water (Cytiva cat. SH30538.FS) then placed in the Eppendorf Thermomixer R at 40⁰C, 1400rpm for 5-10 minutes. For *i.v.*, *i.p.*, and oral gavage, MTZ and NFP were cooled to RT then directly administered.

B/B Homodimerizer, alternatively known as AP20187 (MedChemExpress cat. HY-13992), was solubilized in DMSO to make a 74.14 mg/ml (50 mM) stock solution stored in 10 ul aliquots at - 80⁰C. *In vivo* injection mix administered via *i.v.*, *i.p.*, or oral gavage contains a final concentration of <0.001% Fast Green, 5% DMSO, 2% Tween 80 (Sigma-Aldrich cat. P4780), and 10% PEG300 (MedChemExpress cat. HY-Y0873) diluted with 0.1 M sodium chloride made in Cell and Molecular grade UltraPure Sterile Water. B/B Homodimerizer was used at a final *in vivo* concentration of 10 mg/kg across all experiments.

### Cloning and transgenesis

#### Ubiquitous NTR 2.0 expression plasmids

To make the CMV: NTR 2.0-P2A-tdTomato plasmid (JG232) the enhanced nitro reductase (NTR 2.0) coding sequence was synthesized (IDT) corresponding to Addgene plasmid # 158653 provided as gift from Jeff Mumm (Addgene plasmid # 158653) as an axolotl codon optimized gBlock (IDT) fused to a P2A peptide and in frame tdTomato coding sequence (obtained from FPbase.org). Using Gibson assembly (NEB), this gBlock was inserted into the PmeI/ BsrG1 linearized (transgenic compatible) destination vector containing I-SceI/Tol2 sites pBSII-SK-mTol2-MCS, received as a gift from Elly Tanaka (Addgene plasmid # 51817). A ubiquitous tdTomato control plasmid CMV:tdTomato (JG319) was constructed by removal of the NTR 2.0 XhoI-XhoI fragment from CMV: NTR 2.0-P2A-tdTomato plasmid (JG232 and vector re-ligation with T4 ligase (NEB). Ligated constructs were transformed into DH5alpha competent cells (Zymo Research). Positive clones were screened by complete sanger sequencing for all final constructs.

#### Ubiquitous iCasp9 expression plasmids

To make the *CMV:ihCasp9-T2A-mCherry plasmid (JG228)* a synthetic gBlock purchased form IDT containing the 110aa DmrA dimerization domain fused to the 286aa human Caspase 9 (FKBP12 F36 to V Truncated Caspase9 (135-416; lacking pro-CARD domain) with a 6aa linker and a 9 aa HA tag based on the Addgene plasmid # 15567 a kind gift from David Spencer^20^. To improve the brightness of the fluorescence and allow direct comparisons in vitro, mCherry was later replaced with gBlock containing the tdTomato sequence direct Gibson assembly (NEB) to make CMV:ihCasp9-T2A-tdTomato (JG361). To make an axolotl specific iAxCasp9 expression plasmid, CMV: iaxCasp9-T2A-tdTomato (JG383), a gBlock predicted to have similar function to the hiCasp9 was synthesized (IDT) with the 110aa DmrA dimerization domain fused to the 284aa Axolotl Caspase 9 axolotl orthologue (Axo Casp9_379aa_ AMEX60DD301051603.3 caspase-9 isoform X1 (CASP9) with CARD domain deleted in silico) with a 6aa linker and a 9 aa HA tag appended. These gBlocks were ligated via a direct Gibson assembly (NEB) into SceI meganuclease containing parent vector pBSII-SK-mTol2-MCS received as a gift from Elly Tanaka (Addgene plasmid # 51817) modified with a gBlock to include a CMV promoter/enhancer driving expression of T2A peptide in frame with mCherry. Ligated constructs were transformed into DH5alpha competent cells (Zymo Research). Positive clones were screened by complete sanger sequencing for all final constructs.

#### Isolation and cloning of the axolotl CD68 promoter

The axolotl CD68 promoter was isolated by targeting 3400bp upstream of the Axolotl coding sequence plus 49bp of exon1+276bp of first intron predicted to contain macrophage-specific transcription factor binding motifs (http://homer.ucsd.edu/homer/motif/). Primers were designed using the UCSC - Axolotl genome browser gene model CD68:AMEX60DD301012740^78^ ^79^. Axolotl genomic DNA was isolated using the DNeasy Blood and tissue kit from Qiagen as per manufacturer’s instructions. Genomic PCR was performed using Takara PrimeStar in 50ul with approx. 500ng genomic DNA for 40 cycles using a touchdown PCR protocol with a primary PCR followed by a nested PCR (AxoCD68genFWDPrimer1: CATAAAGAAATTATTGCCATTACAGGTC, AxCd68genREVprimer1: GTAACTTTTGGAGATGGGAGATAGG, AxCD68genFWDnested2: ACTAGTGACTTTTCTGGTCTCTGTCTTT, AxCD68genREVnested 2: ACAGTGTGGAGTAGCCAGAGAGAT) Candidate bands were cloned into pCR4Blunt-TOPO (Invitrogen) with inserts sequenced. To make the CD68:NTR 2.0-P2A-tdTomato construct (JG346), the CD68 promoter SpeI/Not fragment was used to replace the CMV promoter/ enhancer from the CMV: NTR 2.0-P2A-tdTomato plasmid (JG232). To construct the *CD68:mScarlet-IRES-* ihCasp9 (JG341) plasmid, Gibson assembly was used to insert an axolotl codon optimized gBlock of mScarlet (FPbase.com) with a stop codon upstream of an IRES sequence from encephalomyocarditis virus (EMCV) and the ihCasp9 sequence matching the sequence from Addgene plasmid # 15567 (David Spencer) downstream of the IRES. To increase the stoichiometric ratio of iCasp9 relative to the fluorophore, the SpeI/NotI fragment containing the CD68 promoter from *CD68:mScarlet-IRES-*ihCasp9 (JG341) was subcloned by replacing the CMV promoter in CMV:ihCasp9-T2A-tdTomato (JG361) to make *CD68:ihCasp9-T2A-tdTomato* (JG391). To test the in vitro sensitivity of iCasp9 expressed using a T2A or via an IRES sequence we constructed a CMV:ihCasp9-T2A-mScarlet (JG322) using a 3 fragment Gibson cloning of overlapping gBlocks into the CMV: NTR 2.0-P2A-tdTomato plasmid (JG232) backbone cut with MluI/EcoRV. Ligated constructs were transformed into DH5alpha competent cells (Zymo Research). Positive clones were screened by complete sanger sequencing for all final constructs. All ablation plasmids include I-SceI mega nuclease restriction sites that facilitate reliable axolotl transgenesis. Plasmids were used to generate F0 mosaic and eventually F1 germline transgenic axolotls. Axolotl egg injection was performed according to previously published protocols ^80^. Briefly, freshly laid single-cell-stage white d/d embryos were placed in a 2% agar mold filled with 1X MMR/10% Ficoll/Pen-strep solution and injected with 5nl of 500–1,000 pg plasmid mix containing I-SceI using the Nanoliter2020 microinjection system (World Precision Instruments, NANOLITER2020). After 2 h, embryos were carefully transferred to 0.1X MMR/5% Ficoll/Pen-strep solution for 24 h then moved to 24-well plates containing 0.1X MMR/Pen-strep solution for 7-14 days. Axolotl larvae were screened between 7-14 days and positively identified animals were individually housed in 4-ounce plastic cups with 20% Holtfreter’s solution.

### Multi species cytotoxicity assays

#### Transfection

To generate transiently transfected human embryonic kidney (HEK293), murine fibroblast (L929), and axolotl limb dermal fibroblast (AL1) cell lines, we used GenJet Plus (SignaGen Laboratories cat. SL100499) to perform reverse transfections as per the manufacturer’s instructions. Briefly, cells were harvested from 80-90% confluent T-75 flasks (Thermofisher Scientific cat. 156499) using TrypLE Express (Thermofisher Scientific cat. 12605010) to detach the cells for approximately 5 minutes. Complete culture medium for the respective mammalian (10% fetal bovine serum, 1X penicillin-streptomycin, 1X GlutaMAX, 1X DMEM) or axolotl (5% fetal bovine serum, 1X penicillin-streptomycin, 1X GlutaMAX, 1X Insulin-Transferrin-Selenium, 1X MEM diluted to 0.7X with sterile cell culture grade water) cell line was added to halt trypsinization and an aliquot was taken to determine cell density using 0.4% trypan blue. The trypan blue stained cell aliquot was dispensed onto a hemocytometer to count the number of viable cells within a 5x5 square grid by excluding cells within the grid that are trypan blue^+^, which is used as an indicator for nonviable cells. To calculate cell density, the viable cell count was then multiplied by 10^4^, the dilution factor, and the cell suspension volume. The cell suspension was centrifuged at 25°C, 350 rcf for 5 minutes and the supernatant was disposed of with an aspirator. The remaining cell pellet was resuspended in serum-free culture medium without additives and dispensed into 1.5 ml low-bind tubes (Thermofisher Scientific cat. 90410) at the appropriate cell density for reverse transfection. Mammalian (HEK293 and L929) cell lines were at a density of ∼200,000 cells per 1.5 ml tube while the axolotl AL1 cell line had a density of ∼240,000 cells per 1.5 ml tube. The cell containing 1.5 ml tubes were centrifuged at 25°C, 350rcf for 3 minutes and the supernatant was completely disposed of immediately prior to incubation with transfection complex.

The transfection complexes were prepared simultaneously. The following plasmid constructs were diluted in serum-free culture medium without additives to the appropriate concentration depending on the respective cell line: CMV:tdTomato, CMV:NTR 2.0-P2A-tdTomato, CMV:ihCasp9-T2A-tdTomato, CMV:mScarlet-IRES-ihCasp9 (HEK293 only), and CMV:iaxCasp9-T2A-tdTomato (AL1 only). GenJet Plus was diluted in a separate tube with serum-free culture medium without additives and immediately added dropwise to each diluted plasmid containing tube. The tube was vortexed then incubated at RT for 15-20 minutes to allow the transfection complex to form. Both HEK293 and L929 cell lines had a GenJet Plus ratio of 4 ul (GenJet):1 ug (plasmid), whereas the AL1 cell line required a 2:1 ratio. Once the complexes were formed, the appropriate cell containing tube was resuspended in the corresponding transfection complex for 20 minutes. During this incubation period, transfection complex tubes containing mammalian cell lines were placed in a 37⁰C incubator with 5% CO2 while tubes containing axolotl cells were placed in a 25⁰C incubator with 5% CO2. Following incubation with the transfection complex, cells were resuspended in the respective complete cell culture medium, transferred to a 50ml reservoir (Corning cat. 4870), and dispensed into 96-well plates (Greiner cat. 655090) at the appropriate seeding density with a 12-channel pipette. Transfected cells were seeded at either ∼25,000-30,000 (HEK293), ∼15,000-20,000 (L929), or ∼20,000-25,000 (AL1) cells per 96-well. At 10-14 hours post-plating, the transfection complex containing medium was carefully removed and replaced with complete culture medium.

#### *In vitro* drug exposure

The transfected cell lines were monitored until peak expression was detected at either 24 h (HEK293∼80% transfected), 48 h (L929∼40% transfected), or 96 h (AL1∼40% transfected) post-transfection. At this point, 2X master mixes of either metronidazole (MTZ), ronidazole (RON), or AP20187 (B/B) were individually prepared at various concentrations in the complete culture medium associated with the cell line being dosed. NTR 2.0 sensitized cells were exposed to final (1X) concentrations of either MTZ or RON at 8, 40, 200, 1000, 2000, and 5000 µM with a matched DMSO concentration of 0.1%. iCasp9 sensitized cells were exposed to final (1X) B/B concentrations at 0.01, 0.1, 1, 10, and 100 nM with a matched DMSO concentration of 0.1%. A dose-response curve was generated for both NTR 2.0 and iCasp9 sensitized cells relative to the 0.1% DMSO vehicle. The corresponding CMV:tdTomato non-sensitized control cells were exposed to the highest tested concentration of each inducible drug, which was compared to the 0.1% DMSO vehicle condition to assess drug toxicity. Each treatment group included N=3 technical replicates. HEK293 cells were exposed to the inducible drugs for 24 hours. L929 and AL1 cell lines were exposed to the inducible drugs for either 18 or 48 hours. Ablation assays conducted after 48 hours of drug exposure had the media replaced at 24 hours with freshly prepared drug containing media.

#### Dual staining

To determine the state of activation for apoptotic induced cell death, cells were live stained with the commercially available NucView 488 Caspase-3 Substrate and Annexin V CF640R dual apoptosis assay kit (Biotium cat. 30073) as described by the manufacturer. Briefly, the complete culture medium was removed, and cells were washed twice with 1X binding buffer. The cells were then stained with 5 µM NucView 488 Caspase-3 Substrate and 5 µM Annexin V CF640R made up in 1X binding buffer at RT for 20 minutes protected from light. Cells were then washed twice with 1X binding buffer then fixed with 2% PFA containing 1.25 mM CaCl2 diluted with 1X PBS, pH 7.4, for approximately 10 minutes. Cells were then washed twice with 1X PBS containing 1.25 mM CaCl2 and placed in 1X binding buffer for immediate imaging. The 96-well plates were then placed in the dark at 4⁰C for long-term storage.

#### Widefield Zeiss Axio Observer.Z1 Inverted Microscopy

Images of *in vitro* assays (Figure 1B-G, S2-7) were acquired using a widefield Zeiss Axio Observer.Z1 inverted microscope (ref: 431007-9902-000, Carl Zeiss Microscopy, Germany) equipped with an EC Plan-Neofluar 10x/0.30 Air (dry) (ref: 420340-9901-000) objective lens.

Samples were illuminated with a <u>Solid-State Light Source Colibri 7, Type R[G/Y]CBV-UV 7-</u> <u>channel fluorescence light source</u> (ref: 423052-9741-000). DAPI, tdTomato, NucView 488, and CF640R fluorescence was excited and collected by using Zeiss DAPI filter cube set 49 (ref: 488049-9901-000), Zeiss EGFP filter cube set 38 HE (ref: 89038-9901-000), Zeiss mCherry filter cube set 63 HE (ref: 489063-0000-000), and Zeiss Cy5 filter cube set 50 (ref: 488050-9901-000) respectively. Images were acquired with a Zeiss Axiocam 305 color (Ref: 426560-9030-000) controlled with <u>Zen Pro 3.1</u> (Carl Zeiss Microscopy, Germany) software, 3x3 binning, at a resolution of 4378x4488 pixels, in 12 bit and saved in CZI file format. Tiling (N=38) was performed to capture 100% of the well. Images were then stitched using Zen software to fuse the tiles with 5% minimal overlap and 30% maximal shift.

#### Nikon Ti-E YokoGawa CSU-W1 / Nikon C2+ Microscopy

Images of *in vitro* dual staining (Fig. S1) were acquired using a Spinning-disk confocal unit (CSU-W1, Yokogawa, Japan) on a Nikon inverted Ti-Eclipse microscope stand (Nikon Instruments Inc., Japan), equipped with a Plan-Apochromat λ **D 10X/0.45** Air (dry) (ref: MRD 70170) and a Plan-Apochromat λ **20X/0.75** Air (dry) (ref: MRD 00205) objective lens. All images were acquired in 2048*2048 pixels, binning 1x1, in 16bit with a Scientific CMOS Zyla 4.2 (Andor Technology, United Kingdom) controlled with NIS AR 5.41.02 (build 1711, Nikon Instruments Inc., Japan) software and saved in Nd2 file format. *In vitro* dual staining samples (Fig. S1) were placed into a Stagetop Incubator System (Okolab srl, Italy), warmed at 25°C for using the Heated Chamber for Nikon Motorized Stage ( ref: H301-NIKON-Ti-S-ER) and maintain in 5% CO2 using the all in One Controller with C2 Air Mixer (ref: UNO-T-H-CO). Tiling (N=25) was performed to capture 100% of the well in MW-OIL insert. Images were then stitched using NIS software. Fluorescence of *in vitro* dual staining (DAPI, Nucview 488, tdTomato, CF640R), excited with. 405nm, 100% 488nm, 100% 561nm, and 100% 640nm, from a LUNF-XL laser combiner (reff: 77098033) and collected using Nikon DM 445/514/594 (ref: 99226) with DAPI BP 426-446nm (ref: ET436/20x), EGFP BP 500-550nm (ref: FF01-525/50), AF568 BP 579-631nm (ref: ET605/52m), AF647 BP 669-741nm (ref: ET705/72m) emission filter respectively.

Images were exported as tiffs in ZEN Lite then opened as 8-bit tiffs in FIJI for quantification. Masks of the NucView 488 Caspase-3 Substrate and Annexin V CF640R dual apoptosis staining were generated then overlayed with the tdTomato transfected cell mask to determine if cells were healthy or undergoing cell death based on calculated mean grey value. tdTomato^+^ transfected cells were identified as healthy (Caspase-3/Annexin V^-^) if the calculated overlayed mean grey value was less than ∼125. tdTomato^+^ transfected cells were classified as undergoing cell death (Caspase-3^+^, Annexin V^+^, or Caspase-3/Annexin V^+^) if the calculated overlayed mean grey value was greater than ∼125.

A pie chart was made by averaging N=3 wells for each condition for each cell type and transfection construct. The survival curves were made by graphing the average number of total tdTomato^+^ transfected cells and the average number of healthy tdTomato^+^ transfected cells (Caspase-3/Annexin V^-^)

#### Skin grafting ablation assays

Germline (F1) adult CMV:NTR 2.0-P2A-tdTomato (N=1) or CMV:ihCasp9-T2A-mCherry (N=1) transgenic skin donors were placed on a surgical tray ventral side down and kept under anesthetic for the duration of the grafting procedure using a 0.1% tricaine soaked Kimwipe drape (10- to 12-month-old, 13-15 cm length). The donor’s left hindlimb was left uncovered and positioned perpendicular to the central body axis. A total of three rectangular strips of donor skin were sequentially dissected from the distal, central, and proximal regions of the mid-zeugopod. Surgical tweezers were used to gently lift the donor skin while simultaneously using a pair of surgical scissors to cut an approximately 5x1 mm horizontal strip of skin from the mid-zeugopod. The corner of the dissected donor skin was grasped with a pair of surgical tweezers so that the dermis could be carefully peeled or separated from the underlying subcutaneous tissue with minimal vascular disruption. The harvested donor skin was then placed in a 35 mm glass bottom petri dish (MatTek cat. P35G-1.5-20-C) containing 0.7X PBS and set aside until required. After the skin was harvested, a 0.7X PBS soaked Kimwipe was used to cover the donor’s left hindlimb to prevent the area from drying out until the next rectangular strip was required. A total of five 1x1 mm square pieces can be harvested from a single 5x1 mm strip of donor skin within 30–60 minutes of being dissected from the animal.

Recipient d/d axolotls were placed in a 150x15 mm petri dish ventral side down with a 0.1% tricaine soaked Kimwipe drape, leaving the left hindlimb exposed and extended perpendicular to the central body axis (8- to 10-month-old, ∼10 cm length). An approximately 1x1 mm square piece of d/d skin was excised from the central mid-zeugopod of the recipient’s left hindlimb in the same manner as described above. The square piece of recipient skin was promptly disposed of and immediately replaced with a similar sized 1x1 mm piece of donor skin that was cut from the rectangular strip prior to graft placement. The transgenic donor skin graft was positioned on the d/d recipient in the same orientation to maintain proximal-distal identity and reduce cell migration. After grafting, the petri dish containing the recipient animal was covered with a perforated lid to maintain a relatively humid environment and placed at 4⁰C for 1-2 hours. Recipients were carefully transferred to 20% Holtfreter’s solution and animal handling was limited for 3 days post-grafting to minimize risk of graft displacement. At 7 days post-grafting, successful allograft integration was confirmed by restoration of blood flow.

For prodrug delivery through various administration routes (Fig. 3), viable CMV:NTR 2.0-P2A-tdTomato skin allografts (N=8) were selected and allocated to either *i.v.* injection (N=2), *i.p.* injection (N=2), chronic waterborne (WB) exposure (N=3), or acute WB exposure (N=1). Within the *i.v.* and *i.p.* administration routes, half of the animals received a single dose of 100% DMSO while the other half received a single 845 mg/kg dose of MTZ (N=1 per condition) (theoretical *in vivo* concentration of 0.5% DMSO and ∼ 4.9 mM MTZ). For chronic WB exposure, animals were treated with either 1% DMSO (vehicle), 1 mM MTZ, or 1 mM RON for 7 consecutive days with daily replenishment (N=1 per condition). For acute WB exposure, a single 18 hour treatment with 20 mM MTZ was administered and compared to the chronic WB vehicle (N=1 per condition). CMV:NTR 2.0 skin allografts were imaged prior to drug exposure (untreated) and several days following drug administration (1, 2, 3, and 7 days post-treatment).

For MTZ and NFP prodrug delivery solubilized in 10% niacinamide and 5% DMSO through various administration routes (Figure S12), viable CMV:NTR 2.0-P2A-tdTomato skin allografts (N=18) were selected and allocated to either *i.v.* injection (N=6), *i.p.* injection (N=6), or oral gavage (N=6). Within each administration route, grafts were split into three groups: MTZ with niacinamide (N=2), NFP with niacinamide (N=2), or NFP without niacinamide (N=2). Half of these animals were treated with the appropriate vehicle while the other half were treated with the respective prodrug (N=1 per condition). Animals treated with MTZ in the presence of niacinamide received a single *i.v.* injection at 120 mg/kg (0.05% final DMSO *in vivo*, ∼700 µM MTZ), whereas both *i.p.* injection and oral gavage were delivered at a single 400 mg/kg dose (approx. 0.1% DMSO final, 2.3 mM MTZ). All animals treated with NFP prodrug either in the presence or absence of niacinamide received a single *i.v.*, *i.p.*, or oral gavage dose at 6 mg/kg (∼0.1% DMSO, ∼24.4 µM NFP). CMV:NTR 2.0 skin allografts were imaged prior to drug exposure (untreated) and several days following drug administration (1, 2, and 3 days post-treatment).

For B/B delivery through various administration routes (Fig. 3), viable CMV:ihCasp9-T2A-mCherry grafts (N=6) were selected and allocated to either *i.v.* injection (N=2), *i.p.* injection (N=2), or oral gavage (N=2). Within each administration route, half of the animals received a single dose of vehicle (∼0.05% DMSO in vivo) while the other half received a single 10 mg/kg (∼6.74 µM in vivo) dose of B/B (N=1 per condition). CMV:ihCasp9 skin allografts were imaged prior to drug exposure (untreated) and several days following drug administration (1, 2, and 3 days post-treatment).

#### Zeiss Axio Zoom.V16 Microscopy

Images of *in vivo* mosaic larval/juvenile limb ablation (Fig. 2), *in vivo* skin allografting (Fig. 3, S8, S10-14), *in vivo* blastemal grafting (Fig. 4, S15), and *in vivo* CD68 macrophage (Fig. 6, S18-23, S26, S27) were acquired using a Zeiss Axio Zoom.V16 (ref: 435080-9031-000, Carl Zeiss Microscopy, Germany) equipped with a Zeiss Plan Z 1.0x/0.25 objective lens (ref: 435282-9100-000) and a mechanical stage 150*100 Mot (ref: 435465-9000-000) with an insert plate S, glass 237x157x3 mm (ref:435465-9053-000). tdTomato, mCherry, and mScarlet fluorescence were excited with a 120W Metal halide lamp (HXP 120C) and collected by using Zeiss mCherry filter cube set 63 HE (ref: 489063-0000-000). eGFP fluorescence (S15) was excited with a 120W Metal halide lamp (HXP 120C) and collected by using Zeiss eGFP filter cube set 38 HE (ref: 489038-9901). Unilateral dark field images were captured with a transillumination base 300 (ref: 435533-9500-000) equipped with a transillumination top 450 mot (ref: 435500-9000-000). Z-stacks were processed using the Zen module extended depth of focus contrast method with default settings (No Alignment, 7 contrast length scale, 11 smoothing, 0.15 reconstruction). Imaging of *in vivo* skin allografts was performed on a Zeiss Axio Zoom.V16 with a Z-stack image step size of 25 microns (Fig. 3, S8, S10-14).

For short time-lapse video of *in vivo* skin graft vascularization (Fig. S8), images were collected using an exposure time of 200 ms for 6 seconds. All images were acquired using a Zeiss Axiocam 506 color camera (ref: 426556-0000-000) controlled with <u>Zen 3.1 with a 1x1 binning</u> at a resolution of 2752x2208 pixels and saved in CZI format. Images of *in vivo* mosaic larval/juvenile limb ablation were captured with a zoom of 32x (Fig. 2C) or 25x (Fig. 2D) in 14 bit. Images of *<u>in vivo</u>* skin allografting (Figure 3, S8, S10-14) were captured at zoom 100x in 42 bit. Images of *in vivo* blastemal grafting (Fig. 4, S15) were captured at zoom 16x in 14 bit. Images of *in vivo* CD68 macrophage (Fig. 6, S18-23, S26, S27) were captured at zoom 63x in 14 bit.

Quantification of CMV:NTR 2.0-P2A-tdTomato or CMV:ihCasp9-T2A-mCherry skin grafts was performed by individually aligning the graft vascularization at the same position across all captured timepoints. Relative mean fluorescence intensity (MFI) was calculated by drawing a border that encapsulates the entire tdTomato^+^ or mCherry^+^ graft prior to drug exposure (untreated) and applying the same region of interest across every timepoint for each graft. The mean pixel intensity within the defined region was measured in Fiji (white dotted line in Figure 3d or e). The MFI generated for each timepoint was then divided by the MFI of the associated untreated condition and multiplied by 100 to generate the relative MFI as a percentage for each treatment group.

#### Blastema grafting ablation assays

The method for blastemal grafting was adapted and modified from previously published protocols^81^. Briefly, CMV:ihCasp9-T2A-mCherry donor forelimbs were bilaterally amputated in the mid-stylopod region and the bone was trimmed to the equivalent position of the contracted soft tissue to facilitate resorption. Donors were allowed to regenerate until either the early- or mid-bud blastemal stage. Unilateral recipient forelimb amputations were performed as described in donor animals then transferred to a 150x15 mm petri dish ventral side up. The recipient stumps were positioned perpendicular to the body axis using a Kimwipe brace under each forelimb to enhance graft stability and exposed regions of the body were draped with a 0.1% tricaine soaked Kimwipe. Donor blastemal tissue was amputated along with ∼2 mm of stump tissue (shown to contribute to the regenerate by Wells et al.^82^) and placed in the corresponding location of freshly amputated recipients under semi-dry conditions while maintaining proximal-distal identity. Immediately following blastemal grafting, the petri dish containing the recipient animal was covered with a perforated lid to maintain a relatively humid environment and placed at 4⁰C for 1-2 hours. Recipients were carefully transferred to 20% Holtfreter’s solution and animal handling was limited for 3 days post-grafting to minimize risk of graft displacement.

For CMV:ihCasp9 blastemal graft ablation at various regenerative stages (Fig. 4), early-bud blastemal grafts were harvested from bilaterally amputated CMV:ihCasp9-T2A-mCherry donors (N=6 animals, N=2 blastemal grafts harvested per animal, 8-month-old, ∼10 cm length) at 7 DPA and placed onto the freshly amputated stumps of d/d recipients (N=12 animals, N=1 blastemal graft per animal, 8- to 10-month-old, ∼10 cm length). At 24 days post-amputation of the d/d recipient, CMV:ihCasp9 blastemal grafts were imaged prior to drug exposure and classified into one of three regenerative stages: late bud, paddle, or digit (N=4 grafts per regenerative stage). The d/d recipients were equally distributed into either the vehicle (N=2 grafts per regenerative stage, N=6 grafts total) or 15 mg/kg B/B treatment group (N=2 grafts per regenerative stage, N=6 grafts total). Injections were administered via *i.p.* injection every other day from 24 to 28 days post-amputation for a total of three doses. CMV:ihCasp9 graft ablation was captured at 24 (untreated), 28, 30, and 32 days post-amputation.

For CMV:ihCasp9 blastemal graft ablation following a single *i.p.* injection (Fig. S15), mid-bud blastemal grafts were harvested from bilaterally amputated CMV:ihCasp9-T2A-mCherry donors (N=5 animals, N=2 blastemal grafts harvested per animal, 9-month-old, ∼10 cm length) at 13 DPA and placed onto the freshly amputated stumps of CAGGS:GFP recipients (N=10 animals, N=1 blastemal graft per animal, 10-month-old, ∼10 cm length). At 18 days post-amputation of the d/d recipient, CMV:ihCasp9 blastemal grafts were imaged prior to drug exposure then administered a single *i.p.* injection of either the vehicle or 10 mg/kg B/B (N=5 per treatment). CMV:ihCasp9 graft ablation was captured at 18 (untreated), 20, and 22 days post-amputation.

Imaging of *in vivo* blastemal grafts was performed on a Zeiss Axio Zoom.V16 with a Z-stack step size of 100 microns (Fig. 4, S15).

Quantification of CMV:ihCasp9 blastemal grafts was executed by individually aligning the mCherry^+^ stumps at the same position across all captured timepoints. Relative mean fluorescence intensity (MFI) was calculated by drawing a border that encapsulates the entire mCherry^+^ graft (stump as well as regenerate tissue) at every timepoint and measuring the mean pixel intensity within the defined region in Fiji (white dotted line in Fig. 4E). The MFI generated for each timepoint was then divided by the MFI of the associated 24 dpa graft and multiplied by 100 to generate an MFI percentage that is relative to the untreated condition. To calculate the length and surface area of the regenerating CMV:ihCasp9 blastemal graft, the position where the center of the regenerate extends from the mCherry^+^ stump was identified and marked across all timepoints. Regenerate length was then measured by drawing a straight line from this marked position to the most distal end of the regenerate in Fiji. Regenerate surface area was measured by drawing a border that encompasses the tissue that extends beyond the marked position in Fiji.

### Immunofluorescent tissue staining and imaging

Limb tissue samples were fixed in 4% PFA diluted with 1X PBS, pH 7.4, overnight at 4⁰C. The following day, tissues were washed with 1X PBS for 3 x 15 minutes then transferred to 250 mM ethylenediaminetetraacetic acid (EDTA) and stored at 4⁰C for 7 days. Samples were cryopreserved using a 15% sucrose, 15% fish gelatin solution at RT until they were no longer floating then embedded in 15x15x5 mm molds (Fisher Scientific cat. 22363553) containing Tissue Plus O.C.T. compound (Fisher Scientific cat. 23-730-571) through partial submersion in liquid nitrogen. Cryosections of 10µm thickness were made using the Leica CM 1860 UV cryostat at - 20⁰C and collected on Superfrost Plus microscope slides (Fisher Scientific cat. 1255015) and stored at -80⁰C until required.

For immunostaining, sections were rehydrated with 1X PBS and separated using ImmEdge Hydrophobic Barrier (PAP) Pen (Vector Laboratories cat. H-4000). Tissue permeabilization was performed by submerging slides in 0.1% Triton-X (Fisher Scientific cat. AC215680025) for 10 minutes and washed with deionized water for 4 x 5 minutes. Sections were blocked in blocking buffer (10% goat serum, 1% BSA, 0.1% fish gelatin in 1X PBS) for 1 h then incubated with primary antibody in a 1X PBS humidified staining tray overnight at 4⁰C. Primary antibodies used for staining combo include mouse anti-acetylated tubulin (1:200, Sigma cat. t6793-100), rabbit anti-RFP (1:200, Rockland cat. 600-401-379) and chicken anti-IBA1 (1:50, Antibodies inc. cat. IBA1-0020). The next day, sections were washed with 1X PBS for 3 x 5 minutes then incubated with secondary antibody at RT for 2 h. Secondary antibodies include Alexa Fluor 488 anti-mouse IgG2b (1:500, Invitrogen cat. A21141), Alexa Fluor 568 anti-rabbit IgG (1:500, Invitrogen cat. A11011), and Alexa Fluor 647-PLUS anti-chicken IgY (1:500, Invitrogen cat. A32933), respectively. Following secondary incubation, slides were washed with 1X PBS for 3 x 5 minutes and TrueBlack Plus Lipofuscin Autofluorescence Quencher (Biotium cat. 23014) was used as described by the manufacturer. After the last 1X PBS wash, sections were stained with DAPI (1:1000) for 10 minutes, mounted in VECTASHIELD Vibrance Antifade Mounting Medium (Vector Laboratories cat. H-1700-10), and covered with a 22x50 mm – No. 1.5 Microscope Cover Glass (Fisher Scientific cat. 12541024) for imaging.

Tail tissue wholemount samples (∼1 mm) were fixed in 4% PFA diluted with 1X PBS, pH 7.4, overnight at 4⁰C. The following day, tissues were washed with 1X PBS for 3 x 15 minutes then stored in 1X PBS at 4⁰C until required. For wholemount immunostaining, a protocol was adapted from Sabin et al.^74^. Samples were blocked in blocking buffer (10% goat serum, 1% BSA, 0.1% fish gelatin, 0.1% Triton-X, 0.1% Tween-20, 1X Azide in 1X PBS) on the shaker (150rpm, covered) at RT for 24 h then incubated with primary antibody for 4 days under the same conditions. Primary antibodies used in Figure 7 include mouse anti-acetylated tubulin paired with mouse anti-HNK-1 (1:200, Sigma cat. C6680). Primary antibodies used in Supplemental Figure S24 include mouse anti-GFAP (1:200, Sigma cat. MAB360) paired with mouse anti-HNK-1. Primary antibodies used in Supplemental Figure S25 include mouse anti-acetylated tubulin paired with mouse anti-GFAP. After primary antibody incubation, samples were washed with 1X PBST for 4 x 30 minutes then incubated with secondary antibody on the shaker (150rpm, covered) at RT overnight. Secondary antibodies used in Figure 7 include Alexa Fluor 488 anti-mouse IgG2b (1:500) and Alexa Fluor 647 anti-mouse IgM (1:500, Invitrogen cat. A21238), respectively. Secondary antibodies used in Supplemental Figure S24 include Alexa Fluor 488 anti-mouse IgG1 (1:500, Invitrogen cat. A21121) and Alexa Fluor 647 anti-mouse IgM, respectively. Secondary antibodies used in Supplemental Figure S25 include Alexa Fluor 488 anti-mouse IgG2b (1:500) and Alexa Fluor 647 anti-mouse IgG1 (1:500, Invitrogen cat. A21240), respectively. The next day, samples were washed with 1X PBS for 3 x 30 minutes then stained with DAPI (1:1000) for 15 minutes. After staining, samples were washed with 1X PBS for 3 x 30 minutes then mounted in VECTASHIELD PLUS Antifade Mounting Medium (Vector Laboratories cat. H-1900-10). FUN-TAK (Loctite cat. 1087306) placed under a 22x50 mm – No. 1.5 Microscope Cover Glass for imaging.

Victoria blue staining protocol was adapted from Vieira al.^83^ using 10% neutral buffered formalin fixation. Briefly, tail tissue samples were dehydrated for 1 hour in 50% EtOH then 2 hours in acid alcohol (1% HCl diluted in 100% EtOH). Samples were stained with 1% Victoria blue solubilized in acid alcohol for 45 minutes and destained in 70% EtOH for 20 minutes. Tails were subjected to further dehydration by sequential 20-minute incubations with increasing concentrations of EtOH: 90%, 95%, 100%, and 100% EtOH, respectively. Tail tissue samples were then cleared using methyl salicylate (Sigma-aldrich cat. M6752) for 1 day before being transferred onto a glass slide and mounted in methyl salicylate with a 22x50 mm – No. 1.5 Microscope Cover Glass for imaging.

### Nikon Ti-E YokoGawa CSU-W1 / Nikon C2+ Microscopy

Images of 7dpa limb IF staining (Fig. 5, S17), and 21dpa tail whole mount staining (Fig. 7, S24, S25) were acquired using a Spinning-disk confocal unit (CSU-W1, Yokogawa, Japan) on a Nikon inverted Ti-Eclipse microscope stand (Nikon Instruments Inc., Japan), equipped with a Plan-Apochromat λ **D 10X/0.45** Air (dry) (ref: MRD 70170) and a Plan-Apochromat λ **20X/0.75** Air (dry) (ref: MRD 00205) objective lens. All images were acquired in 2048*2048 pixels, binning 1x1, in 16bit with a Scientific CMOS Zyla 4.2 (Andor Technology, United Kingdom) controlled with NIS AR 5.41.02 (build 1711, Nikon Instruments Inc., Japan) software and saved in Nd2 file format. 7dpa limb IF staining (DAPI, Alexa Flour 488, Alexa Flour 568, Alexa Flour 647), and 21dpa tail whole mount staining (DAPI, Alexa Flour 488, Alexa Flour 568, Alexa Flour 647) were excited with. 405nm, 100% 488nm, 100% 561nm, and 100% 640nm, from a LUNF-XL laser combiner (reff: 77098033) and collected using Nikon DM 445/514/594 (ref: 99226) with DAPI BP 426-446nm (ref: ET436/20x), EGFP BP 500-550nm (ref: FF01-525/50), AF568 BP 579-631nm (ref: ET605/52m), AF647 BP 669-741nm (ref: ET705/72m) emission filter respectively. Z-stack images of 7dpa limb IF staining (Fig. 5, S17) and 21dpa tail whole mount staining (Fig. 7, S24, S25) were collected with a step size of either 3 µm (10X) or 0.9 µm (20X) with the Scanning stage (ref: Ti-S-ER) in the GS35-M plate insert.

Widefield Zeiss Axio Observer.Z1 Inverted Microscopy

Images of Victoria blue staining (Figure 6I) were acquired using a <u>widefield</u> <u>Zeiss Axio</u> <u>Observer.Z1 inverted microscope</u> (ref: 431007-9902-000, Carl Zeiss Microscopy, Germany) equipped with a Plan-Apochromat 20x/0.80 Air (dry) (ref: 420650-9901-000) objective lens. Samples were illuminated with a Visible LED (ref: 423053-9030-000), through a Polarizer D (ref: 000000-1121-813), a LD condenser (ref: 424244-0000-000) equipped with DIC slider EC PN 20x/0.50 II (ref: ref: 426940-0000-000) and the Analyzer module Pol ACR P&C for transmitted light (ref: 424937-0000-000). Images were acquired with a Zeiss Axiocam 305 color (Ref: 426560-9030-000) controlled with <u>Zen Pro 3.1</u> (Carl Zeiss Microscopy, Germany) software. Samples had a 1x1 binning at a resolution of 2056x2464 pixels, in 36 bit and saved in CZI file format. Z-stack images were collected with a step size of 1 microns, using the <u>Z-drive mot. Basic</u> (ref: 431007-9902-000).

### CD68 macrophage-specific genetic cell ablation during larval tail regeneration

Tail amputations were performed under a stereoscope (Leica Stereozoom S9i) using a #11 surgical blade (SteriSharps cat. 77-1110) on a #3 handle to remove ∼0.5mm from the distal end of the notochord. Penicillin-Streptomycin (Sigma-Aldrich cat. P4333) was added to the water at a final concentration of 0.01 mg/ml to mitigate the loss of immunocompromised animals due to macrophage-specific genetic cell ablation. Amputated tissues were fixed in 4% PFA diluted with 1X PBS, pH 7.4, overnight at 4⁰C. The following day, tissues were washed with 1X PBS for 3 x 15 minutes then stored in 1X PBS at 4⁰C until required. Animals that had an amputated tail width greater than 135 microns were included in the assay. This was determined by removing an approximately 100-200 µm sliver from the amputation plane of the removed tail tissue and positioning it in a cross-sectional orientation submerged in 0.7X PBS. This view was captured and the central width of the amputated tail tissue was measured for every animal in Fiji.

Germline (F1) CD68:NTR 2.0-P2A-tdTomato and corresponding non-sensitized d/d control animals (Fig. 6, S21-S23, S26) were exposed to either 20% Holtfreter’s (untreated), chronic 1% DMSO (vehicle), 1 mM chronic/10 mM acute MTZ (prodrug), or 1 mM chronic/2 mM acute RON (prodrug) through waterborne (WB) immersion (N=5 animals per treatment, 2-month-old, 2-3 cm length). Chronic WB treatment groups were continuously immersed in specified prodrug from 0 to 14 DPA that was replaced daily. Acute WB treatment groups were subjected to a total of four 18 hour immersions in high concentrations of respective prodrug at 0, 3, 6, and 9 DPA then transferred to 20% Holtfreter’s. Animals were individually housed in 50 ml of respective treatment and placed in a dark tub covered with a perforated lid to minimize light exposure. All animals satisfied the inclusion criteria. Animals were imaged at 0, 2, 7, 14, and 21 DPA.

Germline (F1) CD68:mScarlet-IRES-ihCasp9 animals were distributed into three *i.p.* injection groups: uninjected control (N=4 animals), vehicle (N=5 animals), or 10 mg/kg B/B (N=8 animals) (2-month-old, 2-3 cm length). Within the same experiment, age and size matched non-sensitized d/d control animals were treated with either vehicle or 10 mg/kg B/B (N=5 per treatment) (Fig. 6, S18-S20, S26). Injections were administered every two days from -2 to 20 DPA, alternating between the left and right lower quadrants of the abdominal wall. There were six CD68:mScarlet-IRES-ihCasp9 animals (N=1 uninjected, N=5 B/B treated animals) and four d/d animals (N=2 vehicle, N=2 B/B treated animals) that didn’t meet the inclusion criteria. Therefore, these animals were excluded from our downstream analysis. Animals were imaged at 0, 2, 7, 14, and 21 DPA. Mosaic (F0) CD68:ihCasp9-T2A-tdTomato animals were confirmed to have monocyte expression through epithelial priming prior to amputation (Figure S27). This was conducted by exclusively removing the epithelial tissue from the distal end of the tail at -5 DPA then screening the animals at -3 DPA for CD68:ihCasp9-T2A-tdTomato^+^ recruitment. The best expressors were selected and paired based on the level of mosaicism. Pairs with matched expression were divided into the vehicle or 10 mg/kg B/B treatment group (N=4 animals per treatment, 2-month-old, 2-3 cm length). Tail amputations were performed as described above. Injections were administered every 2-3 days from -2 to 13 dpa, alternating between the left and right lower quadrants of the abdominal wall. Animals were imaged at -5, -3, 0, 2, 7, 10, 13, 17, 24, and 30 DPA.

Imaging of *in vivo* CD68 macrophages in amputated tails was performed on a Zeiss Axio Zoom.V16 Z-stack step size of 9 microns (Fig. 6, S18-23, S26, S27).

Quantification of CD68 tail ablation assays was executed by individually marking and aligning the amputation plane identified at 2 dpa across all captured timepoints for each animal. Regenerate cartilage length was measured by drawing a straight line from the center of the amputation plane to the most distal end of the cartilaginous rod in Fiji. Regenerate spinal cord length was measured by drawing a straight line from the center of the amputation plane to the most distal end of the ependymal tube in Fiji. Regenerate surface area was measured by drawing a border that encompasses the tissue that extends beyond the amputation plane in Fiji.

### Fluorescence-activated cell sorting, cDNA synthesis and qRT-PCR

Limbs stumps harvested at 4 DPA were isolated from CD68: NTR 2.0-P2A-tdTomato or d/d control axolotls (N=3 animals per line, 8- to 10-month-old, ∼10 cm length) and cells dissociated via methods previously described^37^. TdTomato⁺ and non-fluorescent cells were purified by fluorescence-activated cell sorting (FACS) from three animals per experiment. To obtain sufficient material for downstream analysis, cells from all animals were pooled per condition prior to RNA extraction. Cell samples were collected into TRIzol® reagents (Thermo Fisher Scientific). Total RNA was purified using Direct-ZolTM RNA MiniPrep (Zymo Research, Irvine, CA, United States) according to manufacturer’s instructions. RNA quality was assessed by spectrophotometry using a NanoDrop ND-1000 (NanoDrop). Reverse transcription was performed using SuperScript® VILO^TM^ cDNA synthesis kit (Thermo Fisher Scientific). Quantitative real-time quantitative polymerase chain reaction (qPCR) assays were performed using LightCycler® 480 SYBR green (Roche, Indianapolis, IN, United States). Each sample was analyzed in triplicate technical reactions. Ct values were averaged across technical replicates prior to calculation of ΔCt values (target minus reference). Relative expression was calculated using the ΔΔCt method and reported as fold change. This experiment was repeated independently with similar results. Primer sequences used in qPCR assays RbL27-F 5’CATCAGATCAAGCAAGCAGTA , RbL27-R 5’ CCAATGCAGCAGTTTAGATG, Beta Actin-F 5’ TCCATGAAGGCTGCCCAACT, Beta Actin-R 5’ TGGCGCCACATCTGATTGAT, CSF1R-F 5’CTCCAGGATGGGACTGTCAT, CSF1R-R 5’ CCGCTTGGAGGTAGAGTCTG, IGHM-F 5’ CTGAACAGAGGGTGCTCTCC, IGHM-R 5’ TTCTGCAGCTGCTGTGAGTT, CD68-F 5’ AACCAATCGTTGCCTCAGTC, CD68-R 5’ CACCTGGATGGTGAGGTTCT, CSF3R-F 5’ TGACTGTTGAGGAGCGGAAT, CSF3R-R 5’ CTTCTGGTAGAGGCCTCCAC. Primers for B cells were designed against IGHM (IgM heavy chain constant region, GenBank: X68700.1) ^84^

### Single cell RNAseq analsyis

Raw FASTQ read files from the axolotl limb Li et al.^85^ spanning 0 hours post-amputation (hpa) through 33 days post-amputation (dpa) were obtained from the SRA Run Selector (BioProject: PRJNA489484, SRP accession: SRP229913). FASTQ files were analyzed using the nf-core scRNAseq Nextflow pipeline (v4.0.0-ge0ddddb; Nextflow v24.04.4 https://nf-co.re/scrnaseq/4.0.0/) with with the CellRanger option and 10x Genomics Chromium Single Cell 3’ v2 chemistry parameters. Computations were carried out using the Memverge float executor controlling Amazon Web Services batch computing services. Reads were aligned and quantified using the recently published axolotl reference genome GCF_040938575.1_UKY_AmexF1.

Resulting count matrices for each sample were analyzed using a custom Nextflow pipeline, scscape (v4.2.2; Nextflow v24.04.4, https://github.com/mdibl/scscape), which leverages Seurat (see Seurat citations at https://cran.r-project.org/web/packages/Seurat/citation.html) single-cell processing capabilities. Quality control filtering removed cells with fewer than 300 detected genes, mitochondrial content exceeding 10%, and genes expressed in fewer than 3 cells. Cells failing to meet the 10th percentile threshold for either UMI count or gene count were additionally filtered. All other Seurat parameters were left as default. Downstream processing included doublet detection and removal, log normalization, feature scaling, principal component analysis (PCA), k-nearest neighbor graph construction, Louvain clustering, and dimensional reduction via UMAP and t-SNE. The resulting Seurat object was converted to a Loupe Browser (https://www.10xgenomics.com/support/software/loupe-browser/latest) object for further analysis.

Gene names for the GCF_040938575 genome assembly were updated by the MDIBL Comparative Genomic Core to more human readable versions using the following procedure. All genes with text descriptions but a gene name of the form LOCn (where n is string of integers) were extracted and passed to Anthropic Claude Sonnet, version 3.5, with the following prompt: “*You are an expert bioinformatician, trying to take a list of axolotl gene descriptions and map them to Human Genes with the same or almost the same descriptions. If you are unable to find the gene to a high degree of certainty in Humans, you should look using the same strategy in Zebrafish, then Mouse, and then finally check in other common Model organisms. You should only return the most accurate gene for each description, and with it, report in a separate column from which organism that gene was pulled, in cases of identical match in multiple organisms prioritizing in order of Human, Zebrafish, Mouse, and everything else. I will give you a list of 1000 gene descriptions, for which you should report in bunches of 100 your output in format of 1. Gene Description, 2. Matched Gene Name, 3. Organism from which the Gene Name was pulled, and 4. Degree of confidence you have in a match. If your confidence in any given match is less than 50%, you should report back NA for that description.*” For Genes that weren’t found to have a match by Claude, the UniProt (https://uniprot.org/, citation https://doi.org/10.1093/nar/gkae1010) was manually searched using the text gene descriptions. If an organism had a gene with a close match of the gene description, the gene name was transferred, with confidence level "manual". If no organism had a similar gene, the gene symbol was left as unknown.

## Statistical Analysis

Statistical analyses were performed using the GraphPad Prism Software v10 (GraphPad Software, San Diego, CA, United States). The data is always shown as mean values ± SD. Analyses of significant differences between means were performed using either unpaired student t-tests to compare two treatment groups or one-way ANOVA with Dunnett’s multiple comparisons test, one-way ANOVA with Tukey’s multiple comparisons test, and two-way ANOVA with Tukey’s multiple comparisons test. Differences between groups were considered significant at four different levels, including: * *p* < 0.05, ** *p* < 0.01, *** *p* < 0.001, **** *p* < 0.0001.

## Acknowledgements

We acknowledge the MDI Biological Laboratory’s (MDIBL’s) Light Microscopy Facility (LMF) and Frederic Bonnet for widefield and confocal imaging assistance. We thank Joel Graber, Ryan Seaman and the Comparative Genomics and Data Science Core for Bioinformatics support. We thank Will Schott, Krystal-Leigh Brown and Danielle Littlefield at JAX’s Flow Cytometry Service for their support and professional cell sorting assistance. We also would like to thank Stephen Sampson, Iain Drummond and Joel Graber for careful reading of the manuscript. We acknowledge the use of BioRender software for assistance in cartoon/figure design.

## Author contributions

G.J. performed transgenesis, live imaging, tissue grafting and *in vivo* cell ablations and image processing and Fiji quantification. G.J. performed tissue staining, spinning disk confocal imaging and image processing. G.J. performed cell transfections and live/dead cell viability assays and widefield imaging. G.J. assisted in data interpretation, data analysis, manuscript preparation, and most of the figure generation. A.H. performed all cloning and RT-PCR assays along with Fiji quantification and graphing of *in vitro* assays. J.H.G Graber and M.S. provided bioinformatic support. J.W.G. and G.J. performed tissue dissociation and FACS cell purification for gene expression analysis. J.W.G. conceived the study, generated the hypothesis, designed experiments, analyzed data, assisted with figure assembly and wrote/edited the manuscript and provided funding.

## Data Availability Statement

The raw data supporting the conclusions of this article will be made available by the authors, without undue reservation.

