## Supplemental Figures S1-S27 for "Two axolotl-adapted cell-ablation platforms reveal macrophage-dependent processes essential for spinal-cord and skeletal regeneration"

**Contents:**
**Supplementary Figures and legends (S1-S27)**

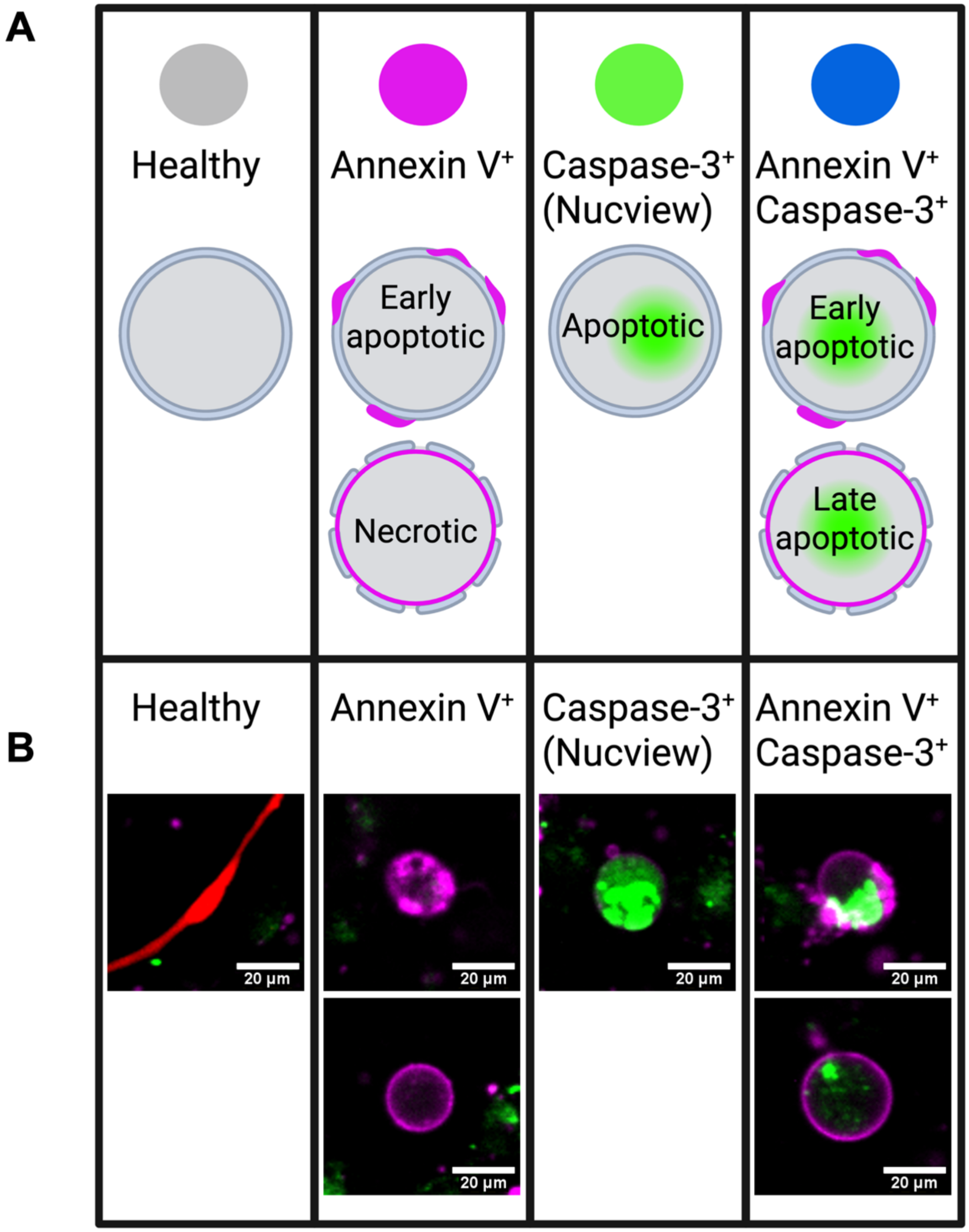


**Figure S1. Dual apoptosis assay distinguishes multiple cell death modalities.**
**(A)** Schematic diagram illustrating expected staining patterns for cell stages defined as healthy, early apoptotic (Annexin V$^{+}$ or Caspase-3/Annexin V$^{+}$ with membrane integrity), necrotic (Annexin V$^{+}$ without membrane integrity), apoptotic (Caspase-3$^{+}$), or late apoptotic (Caspase-3/Annexin V$^{+}$ without membrane integrity).
**(B)** Representative fluorescence images of live dual-apoptosis staining (Caspase-3/Annexin V) in transiently transfected CMV:ihCasp9-T2A-tdTomato (sensitized) axolotl (AL1) cells at 48 hours post-treatment with 0.01 nM AP20187 (B/B; ihCasp9 homodimerizer).
Images demonstrate all detectable cell death phenotypes used for quantifying cell survival after genetically targeted cell ablation. The same patters were obtained with HEK293 and L929 cells. Note: necrotic phenotype was extremely rare but is shown to demonstrate detection capability. Scale Bar: 20 µm.

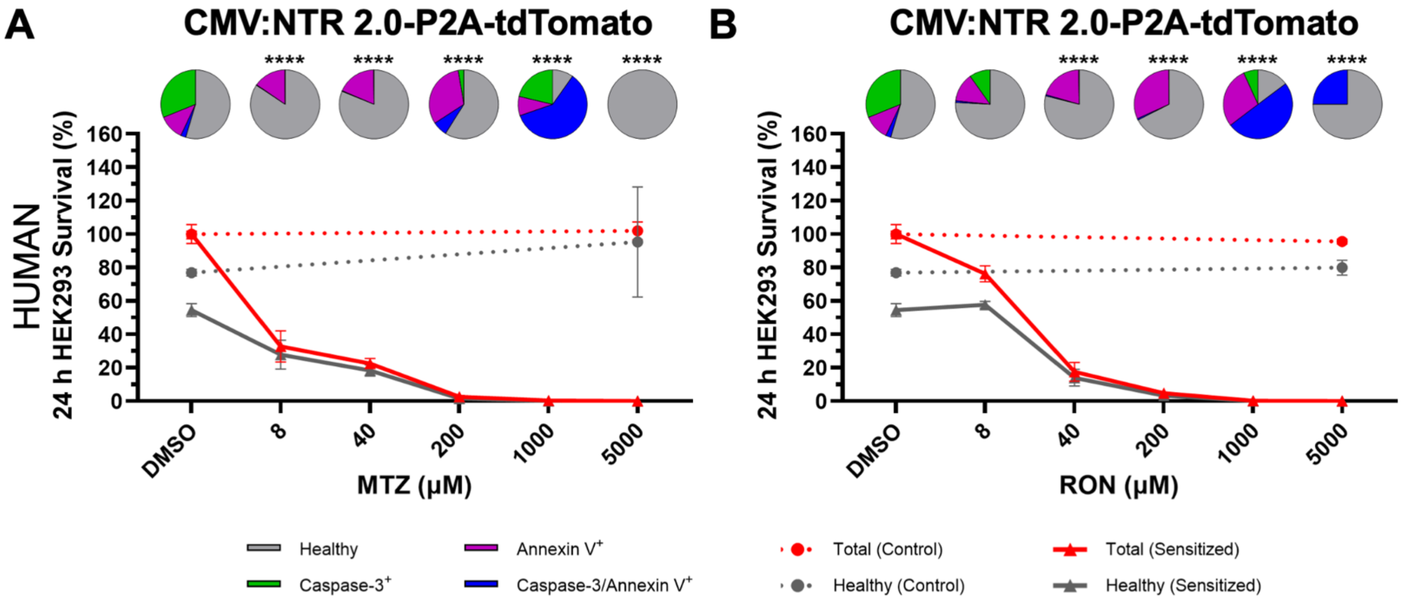


**Figure S2. Comparative prodrug sensitivity: metronidazole versus ronidazole in NTR 2.0-sensitized human HEK293 cells.** Dose-response curves showing survival of CMV:NTR 2.0-P2A-tdTomato sensitized (solid lines) versus CMV:tdTomato control (dotted lines) human HEK293 cells relative to DMSO vehicle. Red lines: total transfected cell counts; gray lines: healthy transfected cell counts (Caspase-3/Annexin V$^-$). Pie charts indicate cell fate distribution: healthy (gray), Caspase-3$^+$ (green), Annexin V$^+$ (magenta), or Caspase-3/Annexin V$^+$ (blue). **(A)** NTR 2.0-sensitized HEK293 cells after 24 h metronidazole (MTZ; NTR 2.0 prodrug) exposure. **(B)** NTR 2.0-sensitized HEK293 cells after 24 h ronidazole (RON; NTR 2.0 prodrug) exposure. Error bars represent mean ± SD (N=3 wells per treatment). No statistically significant difference was observed between the DMSO vehicle and the maximum 5000 µM MTZ or RON concentration in healthy non-sensitized tdTomato control cells (unpaired t-test). Statistical significance of each prodrug concentration relative to the DMSO vehicle was determined in healthy NTR 2.0-sensitized cells by one-way ANOVA with Dunnett's multiple comparisons test (****$p$<0.0001). Representative of 3 independent experiments.

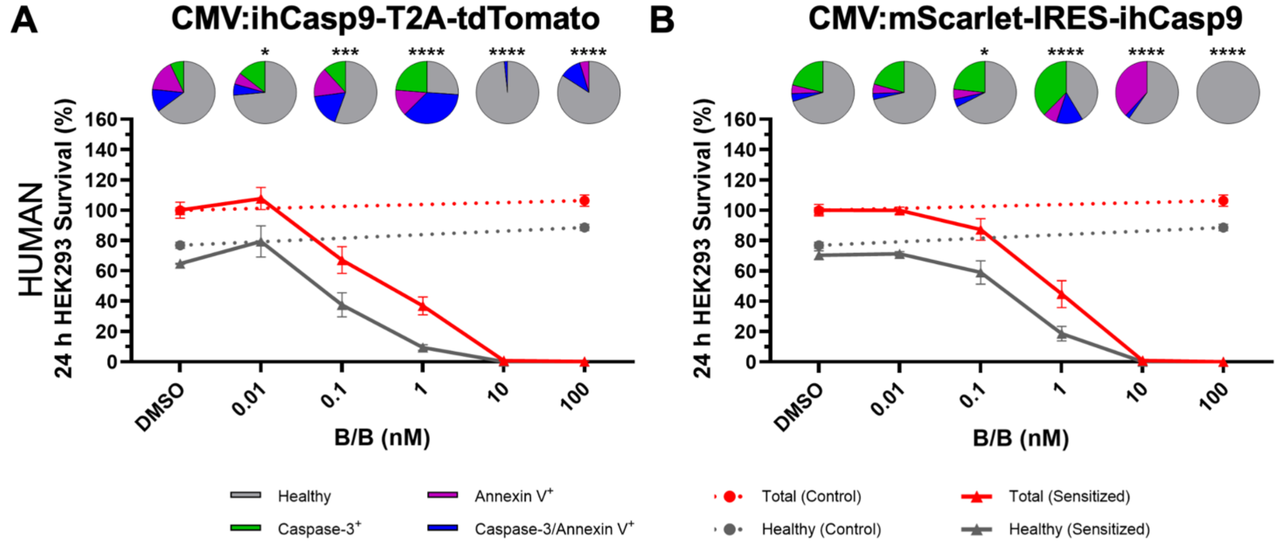


**Figure S3. IRES-ihCasp9 exhibits reduced basal activity compared to T2A-ihCasp9 architecture in human HEK293 cells.** Dose-response curves showing survival of ihCasp9-sensitized (solid lines) versus CMV:tdTomato control (dotted lines) human HEK293 cells relative to DMSO vehicle. Red lines: total transfected cell counts; gray lines: healthy transfected cell counts (Caspase-3/Annexin V$^-$). Pie charts indicate cell fate distribution: healthy (gray), Caspase-3$^+$ (green), Annexin V$^+$ (magenta), or Caspase-3/Annexin V$^+$ (blue). **(A)** CMV:ihCasp9-T2A-tdTomato-sensitized HEK293 cells after 24 h B/B exposure. **(B)** CMV:mScarlet-IRES-ihCasp9-sensitized HEK293 cells after 24 h B/B exposure. Error bars represent mean ± SD (N=3 wells per treatment). No statistically significant difference was observed between the DMSO vehicle and the maximum 100 nM B/B concentration in healthy non-sensitized tdTomato control cells (unpaired t-test). Statistical significance of each B/B concentration relative to the DMSO vehicle was determined in healthy ihCasp9-sensitized cells by one-way ANOVA with Dunnett's multiple comparisons test (*$p$<0.05, ***$p$<0.001, ****$p$<0.0001). Representative of 3 independent experiments. B/B=AP20187 (ihCasp9 homodimerizer).

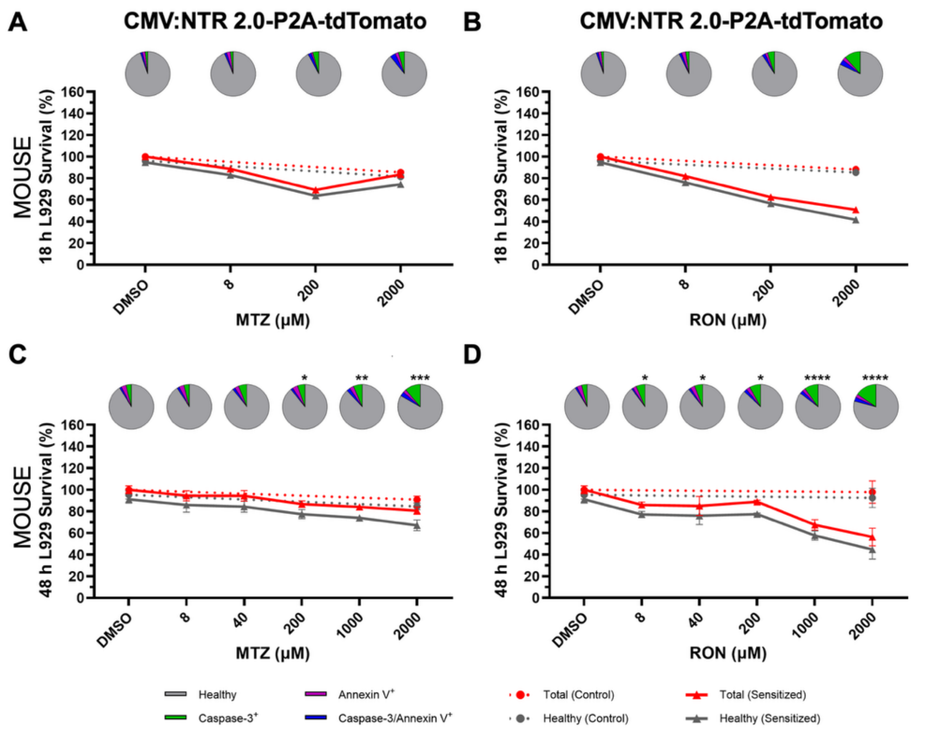


**Figure S4. Ronidazole demonstrates enhanced sensitivity over metronidazole in L929 mouse fibroblasts.**
Dose-response curves showing survival of CMV:NTR 2.0-P2A-tdTomato sensitized (solid lines) versus CMV:tdTomato control (dotted lines) mouse L929 cells relative to DMSO vehicle. Red lines: total transfected cell counts; gray lines: healthy transfected cell counts (Caspase-3/Annexin $V^{-}$). Pie charts indicate cell fate distribution: healthy (gray), Caspase-$3^{+}$ (green), Annexin $V^{+}$ (magenta), or Caspase-3/Annexin $V^{+}$ (blue). **(A, C)** NTR 2.0-sensitized L929 cells after 18 h or 48 h metronidazole (MTZ; NTR 2.0 prodrug) exposure. **(B, D)** NTR 2.0-sensitized L929 cells after 18 h or 48 h ronidazole (RON; NTR 2.0 prodrug) exposure. Data collected at 18 h are presented descriptively without statistical analysis (N=1 well per treatment). Error bars at 48 h represent mean ± SD (N=3 wells per treatment). No statistically significant difference was observed between the DMSO vehicle and the maximum 2000 μM MTZ or RON concentration in healthy non-sensitized tdTomato control cells at 48 h (unpaired t-test). Statistical significance of each prodrug concentration relative to the DMSO vehicle was determined in healthy NTR 2.0-sensitized cells at 48 h by one-way ANOVA with Dunnett's multiple comparisons test (*$p<0.05$, **$p<0.01$, ***$p<0.001$, ****$p<0.0001$). Representative of 3 independent experiments.

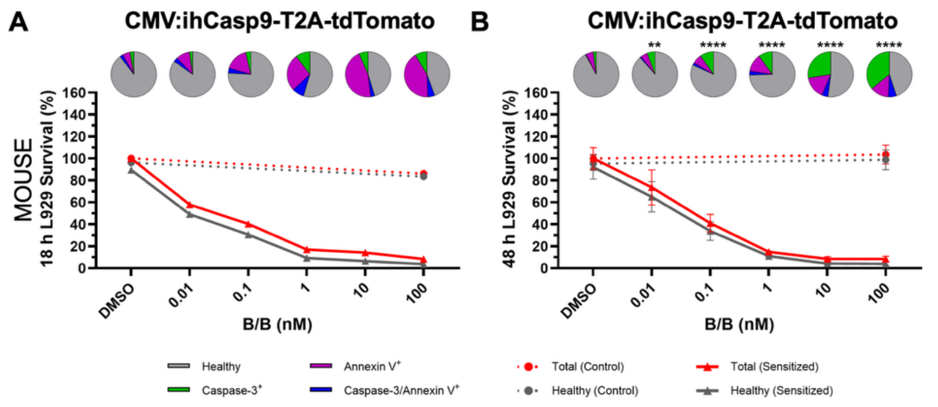


**Figure S5. Temporal dynamics of ihCasp9-mediated ablation in mouse L929 cells.**
Dose-response curves showing survival of ihCasp9-sensitized (solid lines) versus CMV:tdTomato control (dotted lines) mouse L929 cells relative to DMSO vehicle. Red lines: total transfected cell counts; gray lines: healthy transfected cell counts (Caspase-3/Annexin $V^{-}$). Pie charts indicate cell fate distribution: healthy (gray), Caspase-$3^{+}$ (green), Annexin $V^{+}$ (magenta), or Caspase-3/Annexin $V^{+}$ (blue). **(A, B)** CMV:ihCasp9-T2A-tdTomato-sensitized L929 cells after 18 h or 48 h AP20187 (B/B; ihCasp9 homodimerizer) exposure. Data collected at 18 h are presented descriptively without statistical analysis (N=1 well per treatment). Error bars at 48 h represent mean ± SD (N=3 wells per treatment). No statistically significant difference was observed between the DMSO vehicle and the maximum 100 nM B/B concentration in healthy non-sensitized tdTomato control cells at 48 h (unpaired t-test). Statistical significance of each B/B concentration relative to the DMSO vehicle was determined in healthy ihCasp9-sensitized cells at 48 h by one-way ANOVA with Dunnett's multiple comparisons test (**$p<0.01$, ***$p<0.001$, ****$p<0.0001$). Representative of 3 independent experiments.

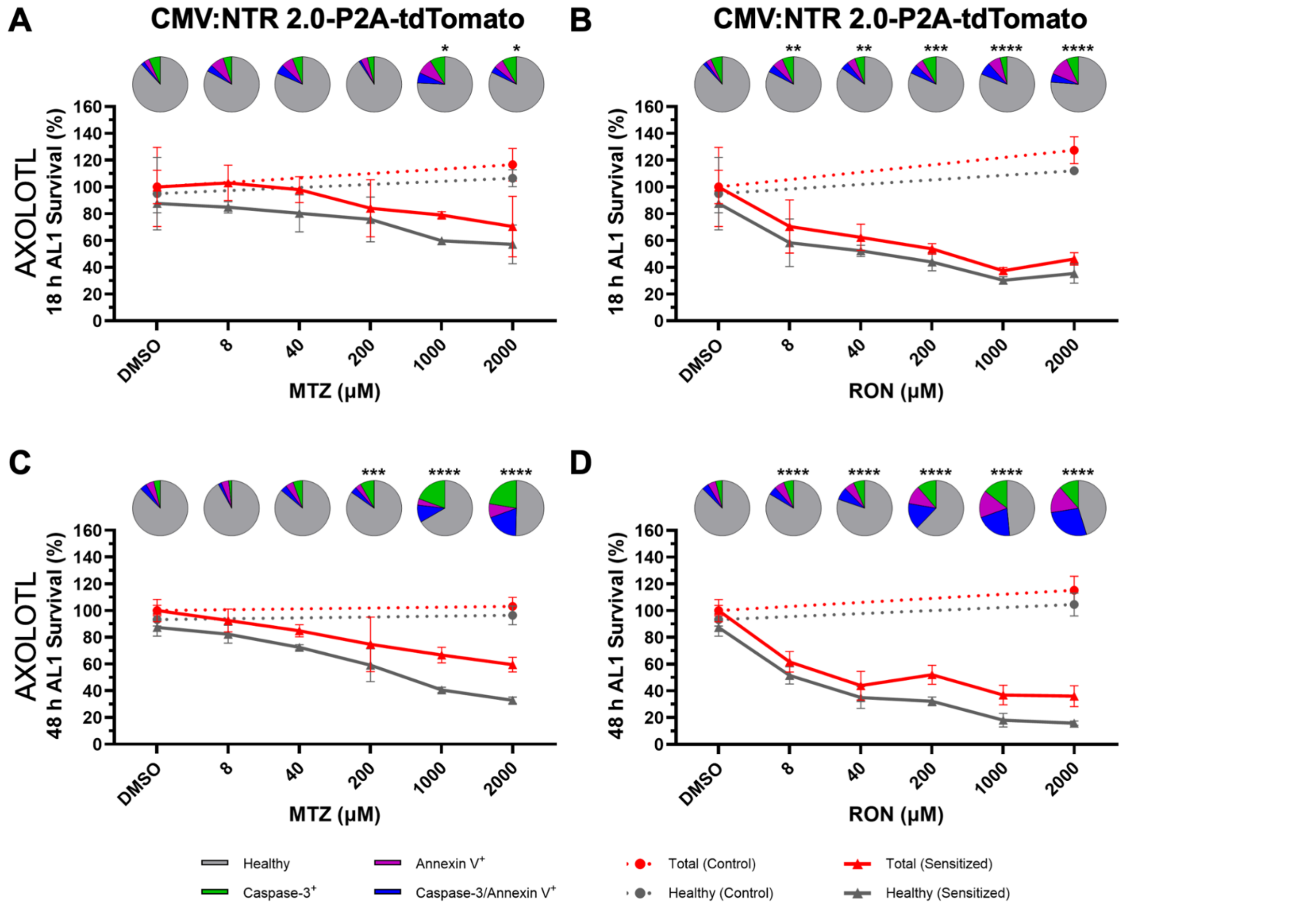


**Figure S6. Ronidazole shows superior efficacy to metronidazole in axolotl AL1 cells.**
Dose-response curves showing survival of CMV:NTR 2.0-P2A-tdTomato-sensitized (solid lines) versus CMV:tdTomato control (dotted lines) axolotl AL1 cells relative to DMSO vehicle. Red lines: total transfected cell counts; gray lines: healthy transfected cell counts (Caspase-3/Annexin V$^-$). Pie charts indicate cell fate distribution: healthy (gray), Caspase-3$^+$ (green), Annexin V$^+$ (magenta), or Caspase-3/Annexin V$^+$ (blue). **(A, C)** NTR 2.0-sensitized AL1 cells after 18 h or 48 h metronidazole (MTZ; NTR 2.0 prodrug) exposure. **(B, D)** NTR 2.0-sensitized AL1 cells after 18 h or 48 h ronidazole (RON; NTR 2.0 prodrug) exposure. Error bars represent mean ± SD (N=3 wells per treatment). No statistically significant difference was observed between the DMSO vehicle and the maximum 2000 µM MTZ or RON concentration in healthy non-sensitized tdTomato control cells (unpaired t-test). Statistical significance of each prodrug concentration relative to the DMSO vehicle was determined in healthy NTR 2.0-sensitized cells by one-way ANOVA with Dunnett's multiple comparisons test (*$p$<0.05, **$p$<0.01, ***$p$<0.001, ****$p$<0.0001). Representative of 3 independent experiments.

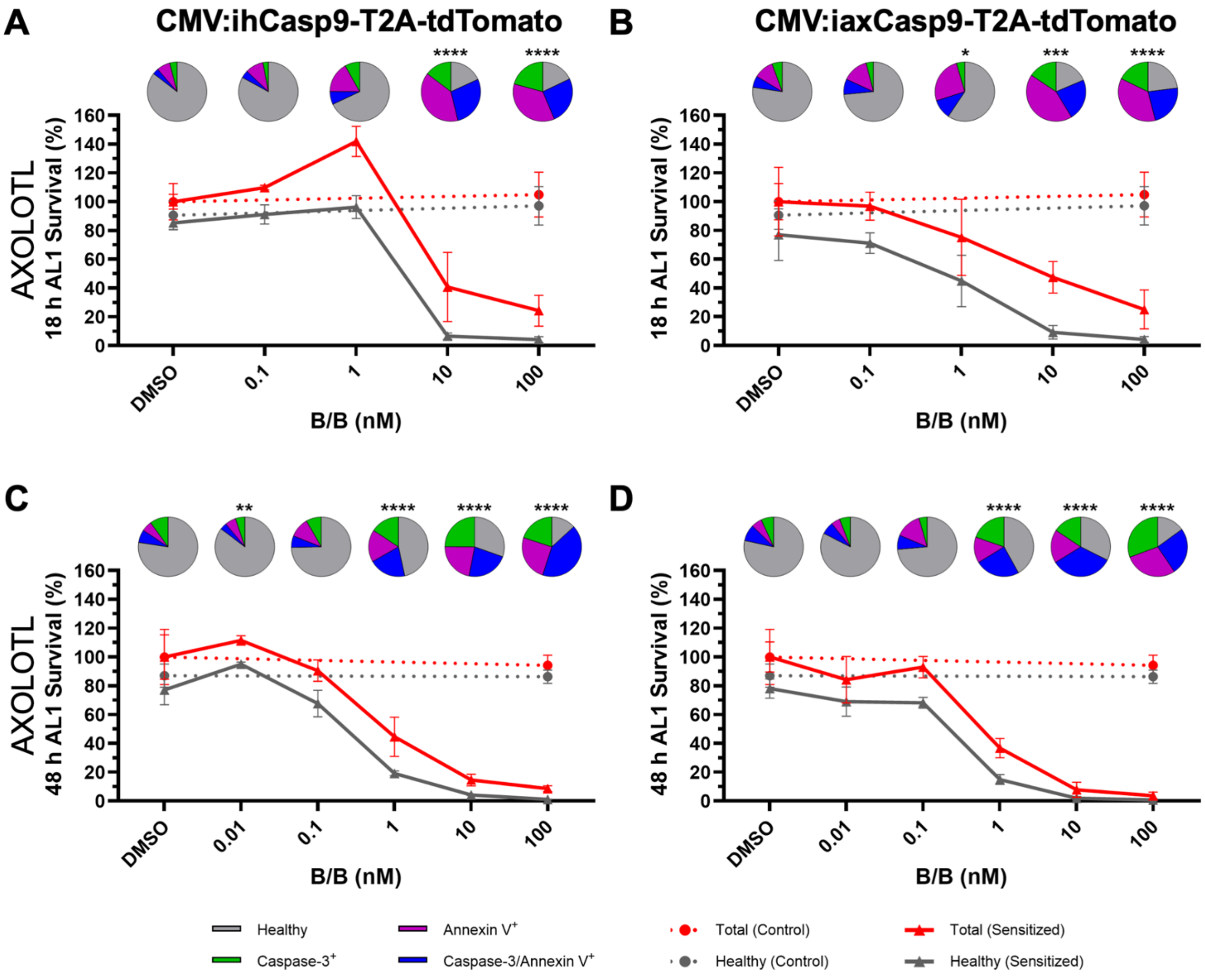


**Figure S7. Human and axolotl-specific iCasp9 variants exhibit comparable ablation efficiency in axolotl AL1 cells.**
Comparative assessment of CMV:ihCasp9-T2A-tdTomato (human Caspase-9 based) and CMV:iaxCasp9-T2A-tdTomato (axolotl Caspase-9 orthologue based) constructs in axolotl AL1 cells *in vitro*. Human construct: 110aa DmrA dimerization domain fused to 286aa human Caspase-9 (FKBP12-F36V-ΔCARDCasp9, residues 135-416) via 6aa linker with C-terminal 9aa HA epitope tag. Axolotl construct: 110aa DmrA fused to 284aa axolotl Caspase-9 (AMEX60DD301051603.3, CARD domain deleted in silico) via 6aa linker with C-terminal 9aa HA tag. Dose-response curves showing survival of ihCasp9- or iaxCasp9-sensitized (solid lines) versus CMV:tdTomato control (dotted lines) axolotl AL1 cells relative to DMSO vehicle. Red lines: total transfected cell counts; gray lines: healthy transfected cell counts (Caspase-3/Annexin $V^-$). Pie charts indicate cell fate distribution: healthy (gray), Caspase-$3^+$ (green), Annexin $V^+$ (magenta), or Caspase-3/Annexin $V^+$ (blue). **(A, C)** ihCasp9-sensitized AL1 cells after 18 h or 48 h B/B exposure. **(B, D)** iaxCasp9-sensitized AL1 axolotl cells after 18 h or 48 h B/B exposure. Error bars represent mean ± SD (N=3 wells per treatment). No statistically significant difference was observed between the DMSO vehicle and the maximum 100 nM B/B concentration in healthy non-sensitized tdTomato control cells (unpaired t-test). Statistical significance of each B/B concentration relative to the DMSO vehicle was determined in healthy ihCasp9- or iaxCasp9-sensitized cells by one-way ANOVA with Dunnett's multiple comparisons test (*$p$<0.05, **$p$<0.01, ***$p$<0.001, ****$p$<0.0001). Representative of 3 independent experiments. B/B=AP20187 (ihCasp9 homodimerizer).

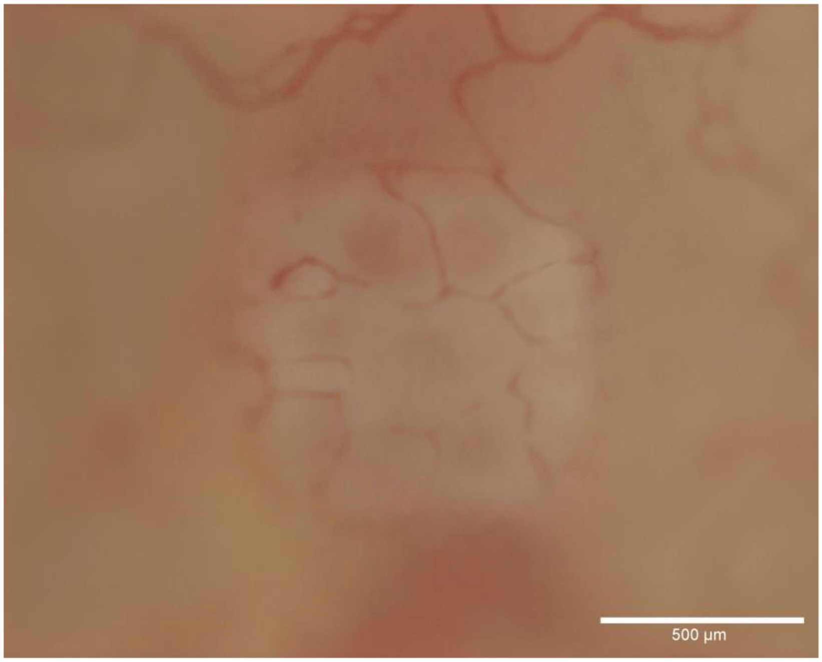


**Figure S8. Validation of skin allograft vascularization prior to drug administration.**
Representative video of blood flow in transgenic skin allografts 7 days post-transplantation, demonstrating complete functional vascularization prior to inducible drug delivery (N=1 animal shown, ~10-month-old, ~10 cm length). Scale Bar: 500 µm.
Video available as supplemental movie.

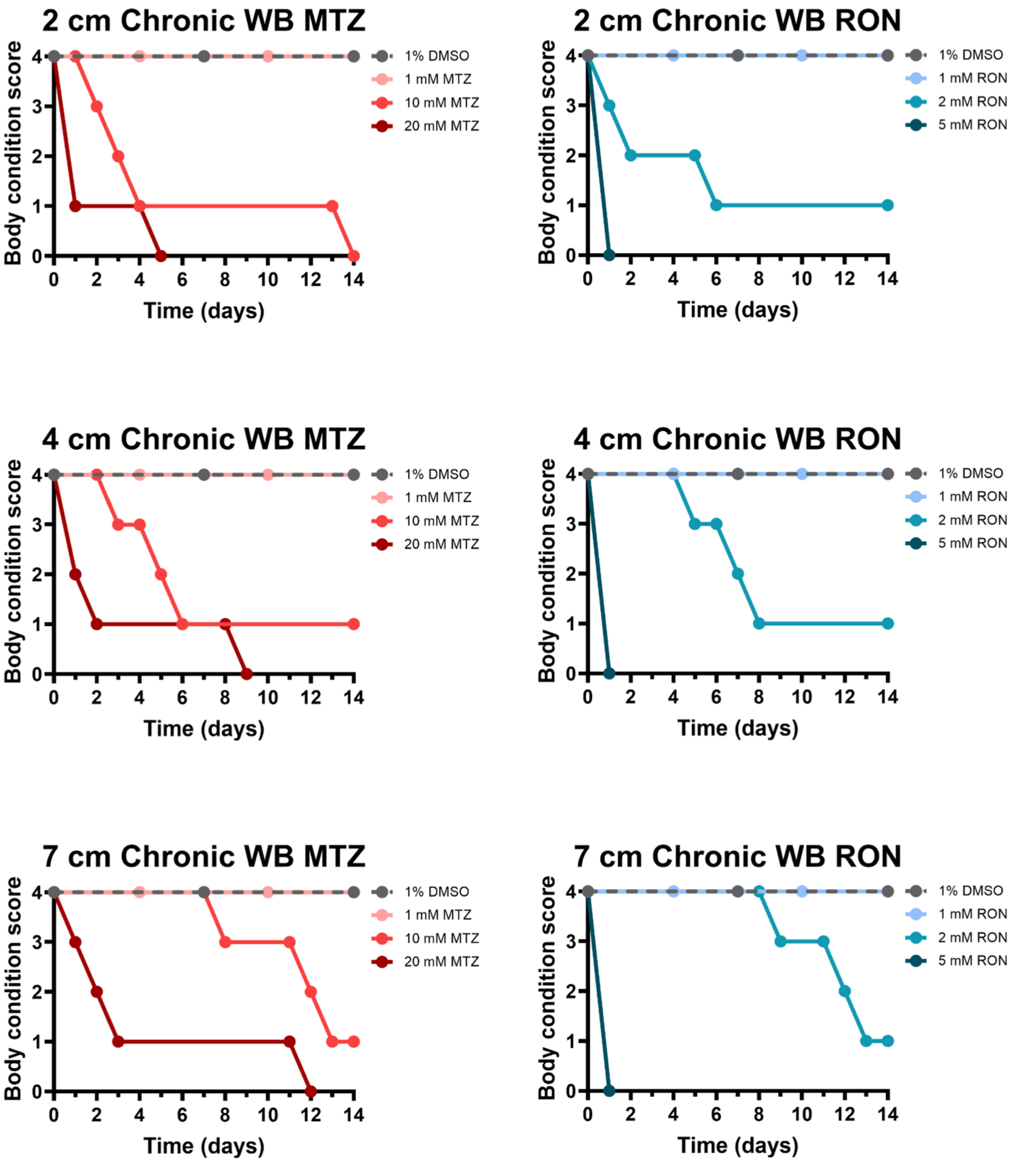


**Figure S9. Size-dependent drug toxicity establishes safe dosing parameters across developmental stages.**
Assessment of metronidazole (MTZ; NTR 2.0 prodrug) and ronidazole (RON; NTR 2.0 prodrug) toxicity through waterborne immersion (WB) to establish a safe dosing rage across three developmental stages of control d/d (non-sensitized) animals as defined by animal snout-to-tail length : 2 cm (~2-month-old larval), 4 cm (~4-month-old juvenile), and 7 cm (~7-month-old adult) (N=3 animals per group). Animals were continuously immersed for 14 days in freshly prepared 1% DMSO vehicle, MTZ (1, 10, and 20 mM), or RON (1, 2, and 5 mM). Animals were scored prior to drug replacement daily using a standardized health assessment: 4=healthy, 3=anemia, 2=malnutrition/food refusal, 1=edema, 0=euthanized for humane reasons. Only the 1 mM concentration for both MTZ and RON were deemed safe for all animal sizes. Note: no inter-animal variation was observed within treatment groups; variation occurred only between treatment groups.

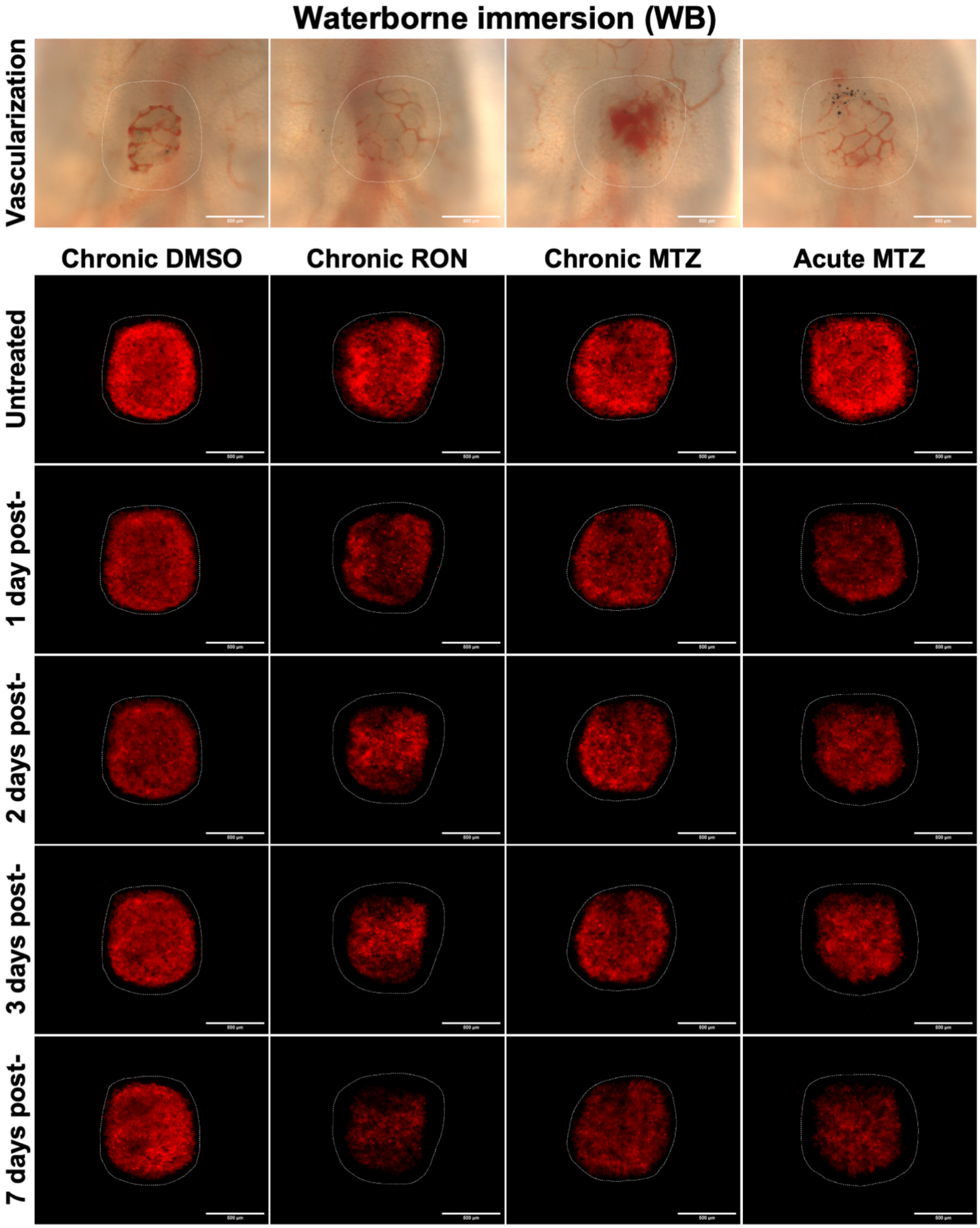


**Figure S10. CMV:NTR 2.0 skin allograft ablation via chronic versus acute prodrug immersion.**
Representative extended depth of focus projections associated with Figure 3B relative mean fluorescence intensity (MFI) quantification data shows time course of skin allograft vascularization for CMV:NTR 2.0-P2A-tdTomato fluorescence (red) at pre-treatment (day 0) compared to 7 days of chronic (1% DMSO, 1 mM RON, 1 mM MTZ) or a single 18 hour acute (20 mM MTZ) waterborne immersion (N=1 per treatment, representative of 3 independent experiments, 8- to 10-month-old, ~10 cm length). Measured MFI within the transgenic skin graft boundary (white dotted line) prior to treatment (day 0) and post-treatment (1, 2, 3, and 7 days) in Fiji. Scale Bar: 500 μm. MTZ=Metronidazole (NTR 2.0 prodrug), RON=Ronidazole (NTR 2.0 prodrug)

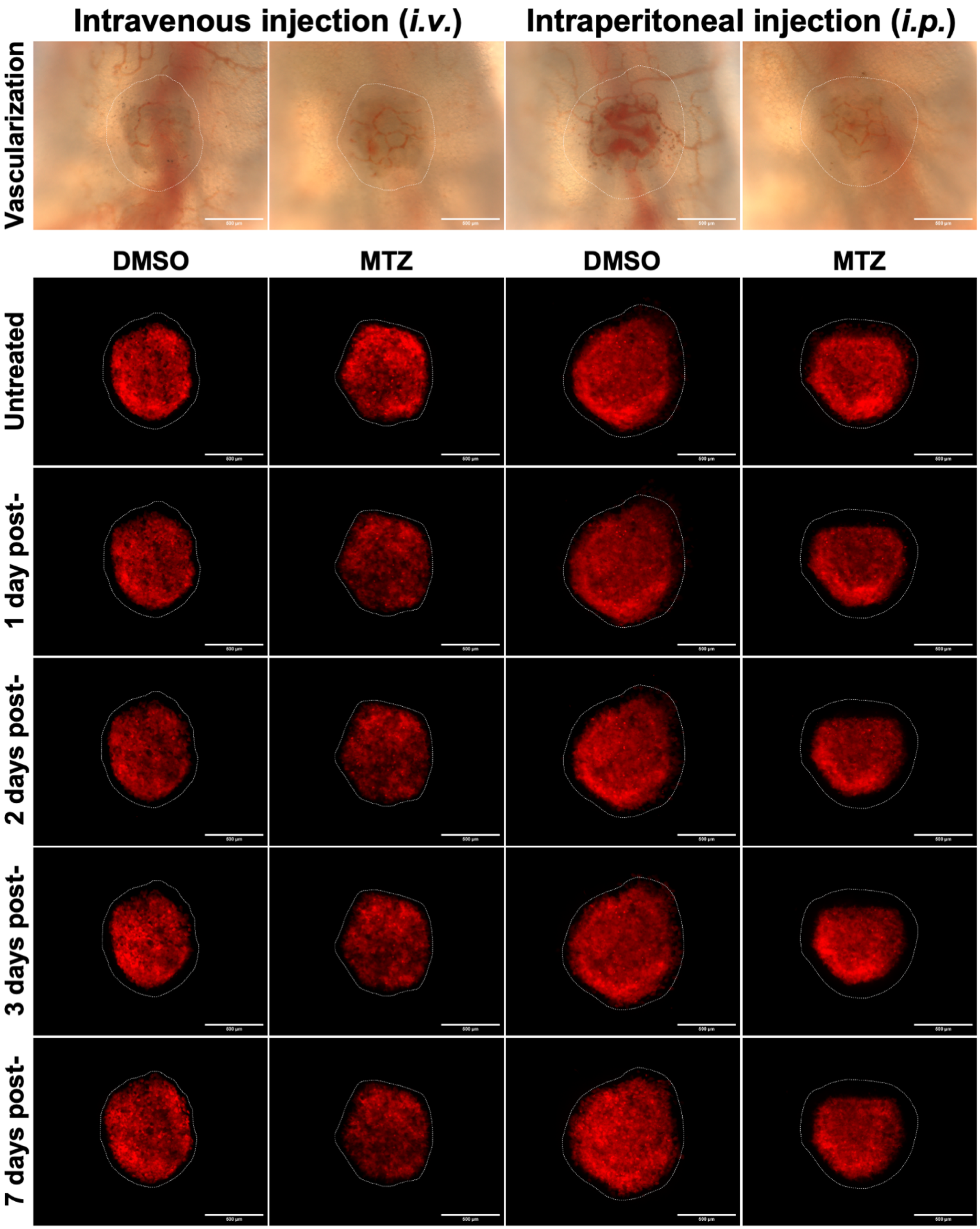


**Figure S11. Systemic MTZ delivery via *i.v.* and *i.p.* injection shows limited efficacy in NTR 2.0 skin allografts.** Representative extended depth of focus projections associated with Figure 3B relative mean fluorescence intensity (MFI) quantification data shows time course of skin allograft vascularization for CMV:NTR 2.0-P2A-tdTomato fluorescence (red) prior to treatment (day 0) compared to subsequent administration of a single DMSO vehicle or 845 mg/kg metronidazole (MTZ; NTR 2.0 prodrug) dose via intravenous (*i.v.*) versus intraperitoneal (*i.p.*) injection (N=1 animal per treatment, 8- to 10-month-old, ~10 cm length). Measured MFI within the transgenic skin graft boundary (white dotted line) prior to treatment (day 0) and post-treatment (1, 2, 3, and 7 days) in Fiji. Scale Bar: 500 µm.

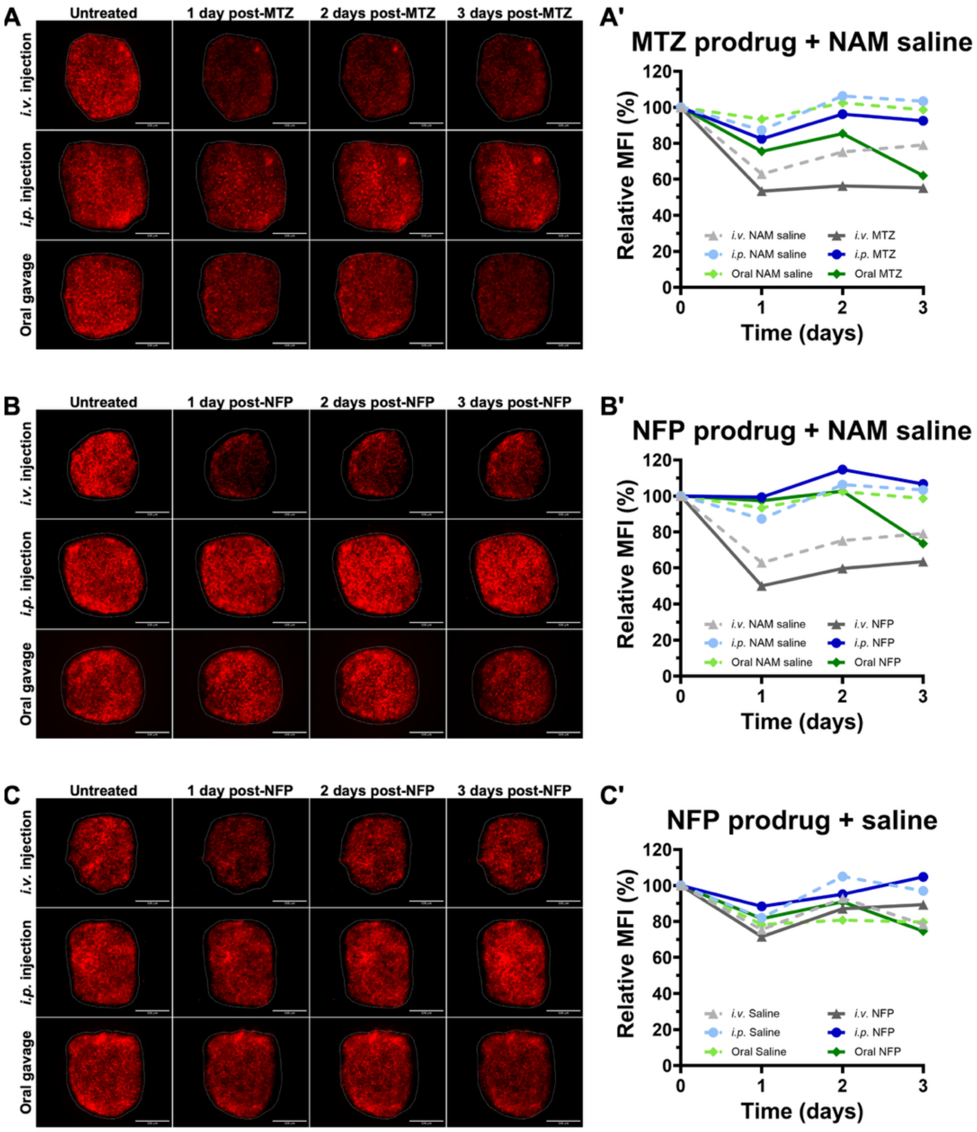


**Figure S12. Niacinamide saline vehicle enhances aqueous MTZ solubility but raises specificity concerns *in vivo*.**
Evaluation of metronidazole (MTZ) and alternative NTR 2.0 prodrug nifurpirinol (NFP) with niacinamide (NAM)-supplemented saline vehicle (5% DMSO + 10% Niacinamide in 0.7x saline) to improve aqueous solubility using CMV:NTR 2.0-P2A-tdTomato (NTR 2.0) donor skin grafts. Assessed ablation efficiency after administering a single dose of respective vehicle or prodrug via intravenous injection (*i.v.*), intraperitoneal injection (*i.p.*), and oral gavage.
**(A)** Representative extended depth of focus projections used to measure (A**'**) relative mean fluorescence intensity (MFI) shows time course of skin allograft vascularization for NTR 2.0 fluorescence (red) prior to treatment (day 0) compared to subsequent administration of MTZ in NAM saline via *i.v.* (120 mg/kg), *i.p.* (400 mg/kg), and oral gavage (400 mg/kg)(N=1 animal per treatment, representative of 3 independent experiments, 8- to 10-month-old, ~10 cm length). Measured MFI within the transgenic skin graft boundary (white dotted line) prior to treatment (day 0) and post-treatment (1, 2, and 3 days) in Fiji. Scale Bar: 500 µm. **(B)** Representative extended depth of focus projections used to measured (B**'**) relative MFI shows time course of skin allograft vascularization for NTR 2.0 fluorescence (red) prior to treatment (day 0) compared to subsequent administration of 6 mg/kg NFP in NAM saline via *i.v.*, *i.p.*, and oral gavage (N=1 animal per treatment, 8- to 10-month-old, ~10 cm length). Measured MFI within the transgenic skin graft boundary (white dotted line) prior to treatment (day 0) and post-treatment (1, 2, and 3 days) in Fiji. Scale Bar: 500 µm.
**(C)** Representative extended depth of focus projections used to measure (C**'**) relative MFI shows time course of skin allograft vascularization for NTR 2.0 fluorescence (red) prior to treatment (day 0) compared to subsequent administration of 6 mg/kg NFP in saline only via *i.v.*, *i.p.*, and oral gavage (N=1 animal per treatment, 8- to 10-month-old, ~10 cm length). Measured MFI within the transgenic skin graft boundary (white dotted line) prior to treatment (day 0) and post-treatment (1, 2, and 3 days) in Fiji. Scale Bar: 500 µm. **(A')** Quantification of MFI in NTR 2.0 skin grafts treated with NAM saline vehicle (dotted lines) or MTZ (solid lines) relative to untreated control following *i.v.* (120 mg/kg), *i.p.* (400 mg/kg), and oral gavage (400 mg/kg) (N=1 animal per treatment, 8- to 10-month-old, ~10 cm length). Measured pre-treatment (day 0) and post-treatment (1, 2, and 3 days) MFI in Fiji. Data are presented descriptively without statistical analysis.
**(B')** Quantification of MFI in NTR 2.0 skin grafts treated with NAM saline vehicle (dotted lines) or 6 mg/kg NFP (solid lines) relative to untreated control via *i.v.*, *i.p.*, and oral gavage (N=1 animal per treatment, 8- to 10-month-old, ~10 cm length). Measured pre-treatment (day 0) and post-treatment (1, 2, and 3 days) MFI in Fiji. Data are presented descriptively without statistical analysis.
**(C')** Quantification of MFI in NTR 2.0 skin grafts treated with saline only vehicle (dotted lines) or 6 mg/kg NFP (solid lines) relative to untreated control via *i.v.*, *i.p.*, and oral gavage (N=1 animal per treatment, 8- to 10-month-old, ~10 cm length). Measured pre-treatment (day 0) and post-treatment (1, 2, and 3 days) MFI in Fiji. Data are presented descriptively without statistical analysis.

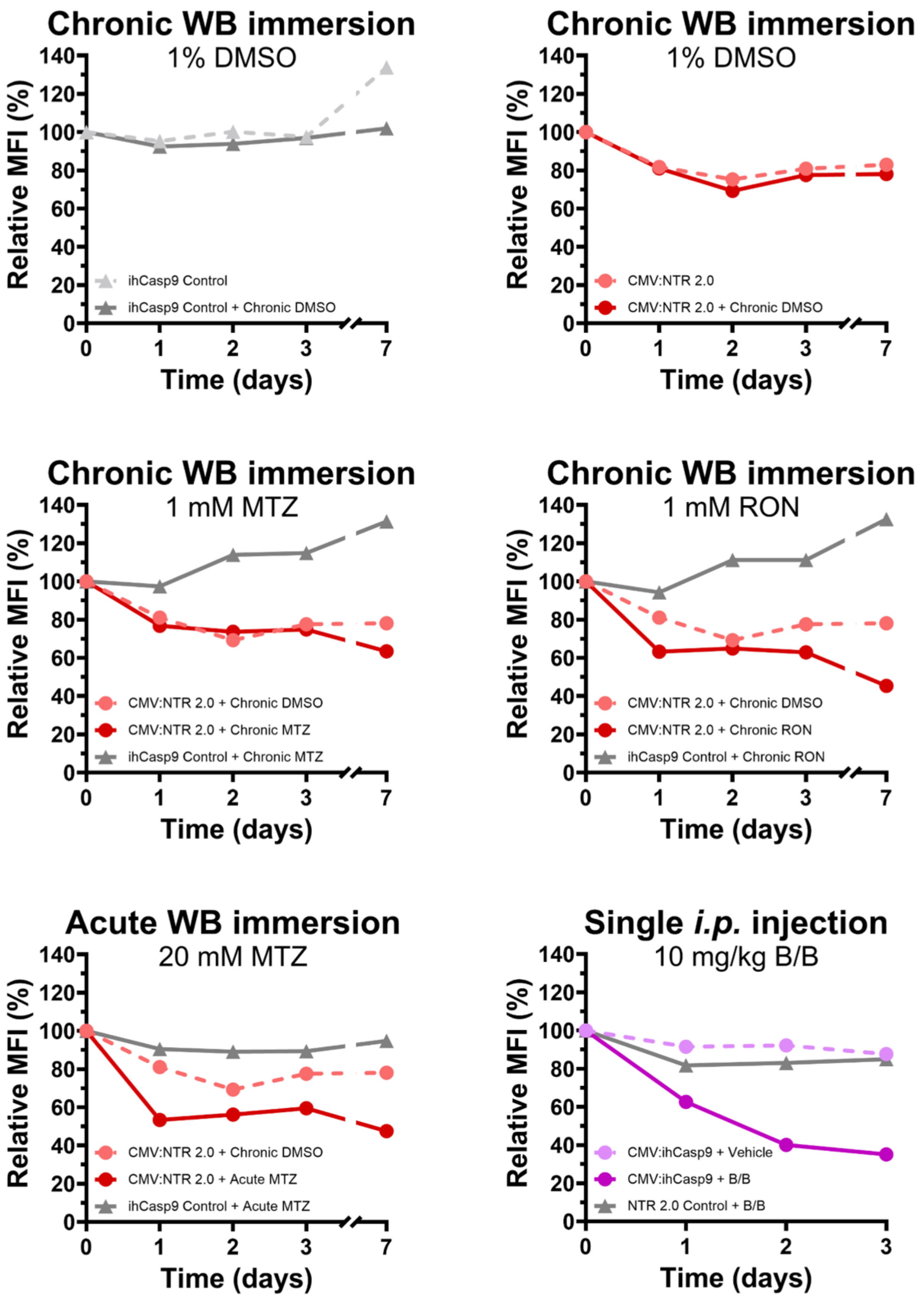


**Figure S13. Cross-system specificity controls confirm selective ablation in sensitized grafts.**
Reciprocal skin allografting experiment demonstrating inducible drug specificity: CMV:ihCasp9-T2A-mCherry (CMV:ihCasp9) grafts were used as controls to show no response to chronic (7 days) or acute (18 hours) waterborne immersion (WB) in MTZ/RON, whereas a single intraperitoneal injection (*i.p.*) had no inherent effect on the CMV:NTR 2.0-P2A-tdTomato (CMV:NTR 2.0) control graft (N=1 animal per treatment, 8- to 10-month-old, ~10 cm length). Full-thickness transgenic skin grafts were transplanted onto wild-type d/d recipients and allowed to vascularize for 7 days prior to drug administration. Treatment conditions tested: chronic WB 1% DMSO, acute WB MTZ (20 mM), chronic WB MTZ (1 mM), chronic WB RON (1 mM), and *i.p.* B/B (10 mg/kg). Mean fluorescence intensity was measured at pre-treatment (day 0) and post-treatment (1, 2, 3, and 7 days) in Fiji. Data are presented descriptively without statistical analysis. MTZ=Metronidazole (NTR 2.0 prodrug), RON=Ronidazole (NTR 2.0 prodrug), B/B=AP20187 (ihCasp9 homodimerizer)

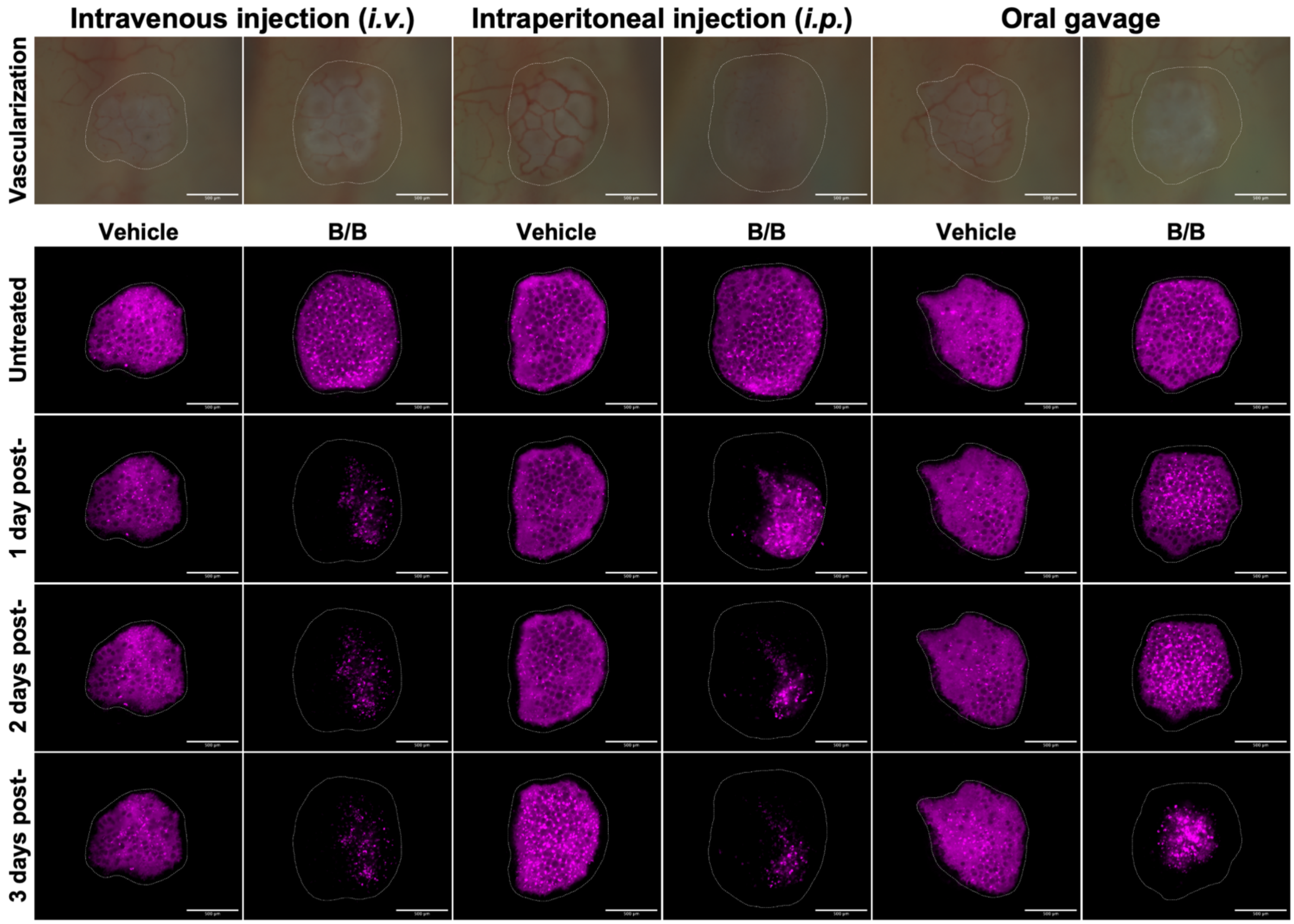


**Figure S14. Multiple administration routes enable effective ihCasp9-mediated ablation in adult skin allografts.** Representative extended depth of focus projections associated with Figure 3C relative mean fluorescence intensity (MFI) quantification data shows time course of skin allograft vascularization for CMV:ihCasp9-T2A-mCherry fluorescence (magenta) prior to treatment (day 0) compared to subsequent administration of a single vehicle or 10 mg/kg B/B dose via intravenous injection, intraperitoneal injection, and oral gavage (N=1 animal per treatment, representative of 3 independent experiments, 8- to 10-month-old, ~10 cm length). Measured MFI within the transgenic skin graft boundary (white dotted line) prior to treatment (day 0) and post-treatment (1, 2, and 3 days) in Fiji. Scale Bar: 500 µm. B/B=AP20187 (ihCasp9 homodimerizer)

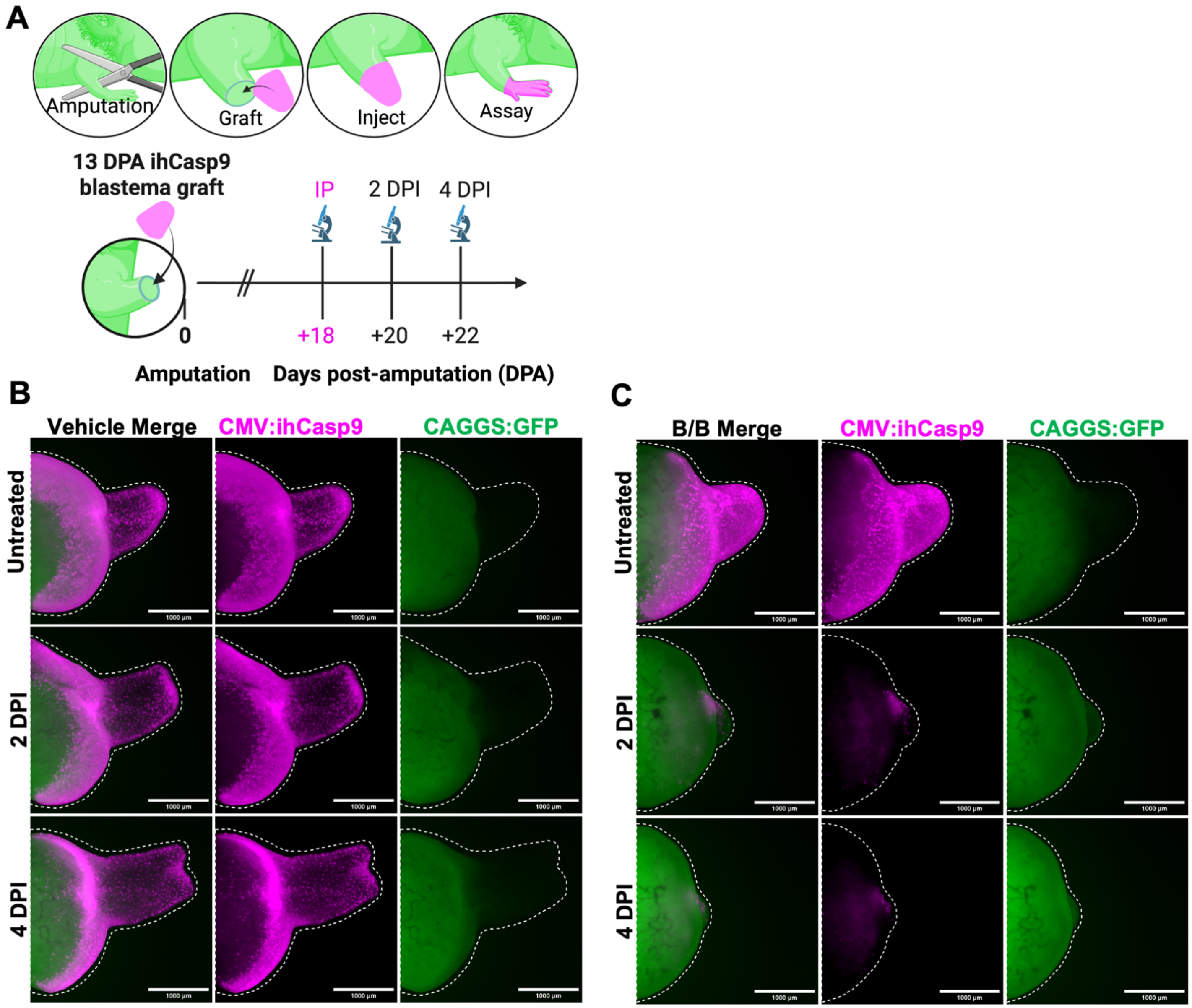


**Figure S15. Single-dose B/B via *i.p.* injection ablates CMV:ihCasp9 blastema grafts with recipient cell replacement.**
CMV:ihCasp9-T2A-mCherry (CMV:ihCasp9) blastemal tissue was harvested at 13 days post-amputation (DPA) and grafted onto CAGGS:GFP transgenic recipients to demonstrate complete ablation within 2 days post-injection (DPI) of a single *i.p.* 10 mg/kg AP20187 (B/B; ihCasp9 homodimerizer) dose, with subsequent replacement by host GFP$^{+}$ cells (N=5 animals per treatment, 9- to 10-month-old, ~10 cm length).
**(A)** Experimental schematic depicting blastema grafting timeline and imaging schedule. **(B, C)** Representative extended depth of focus projections showing mCherry$^{+}$ ablation time-course and GFP$^{+}$ cell infiltration within the CMV:ihCasp9 allograft limb boundary when treated with **(B)** vehicle or **(C)** B/B (white dotted line). Scale Bar: 1000 μm.

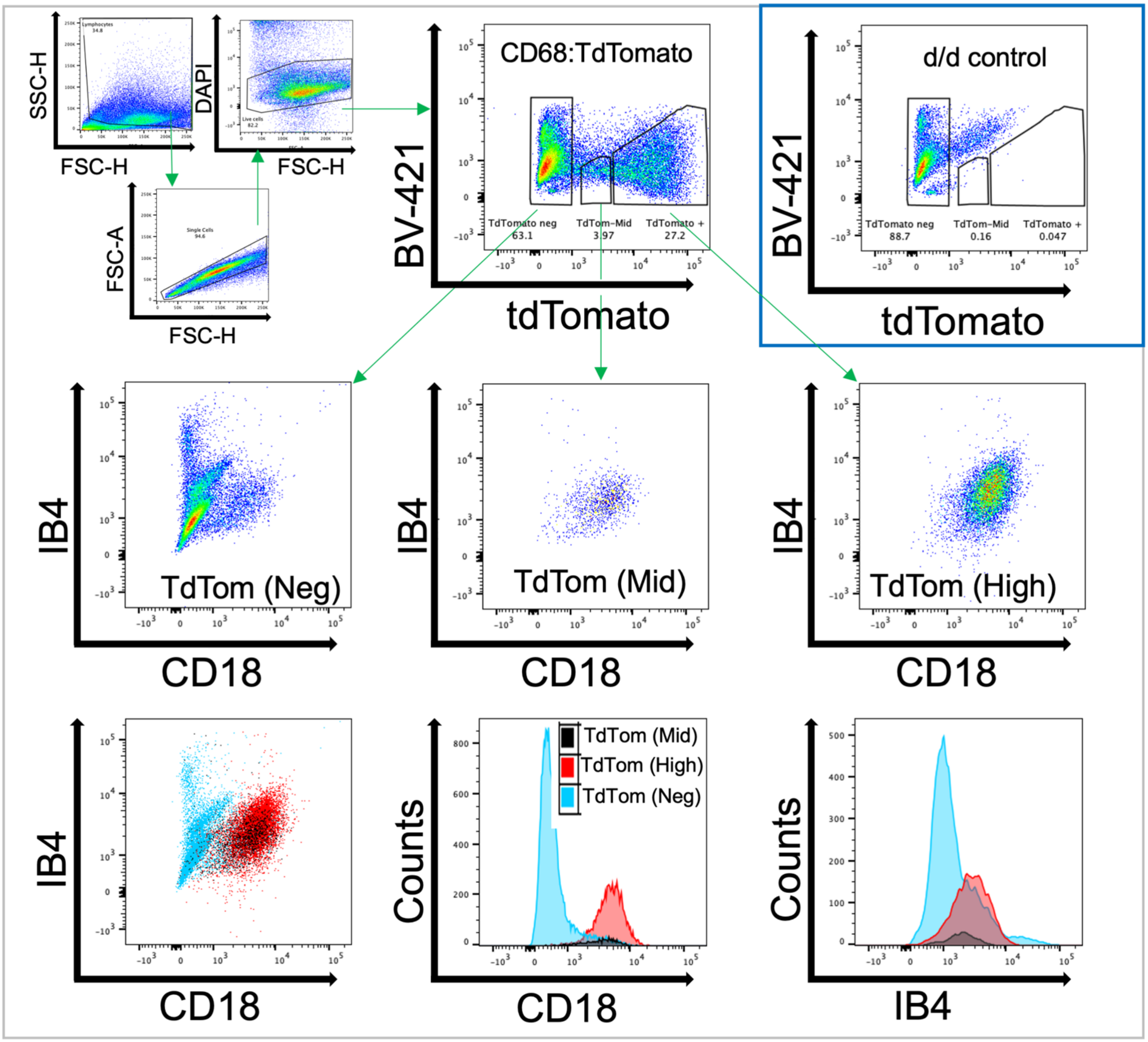


**Figure S16. CD68:tdTomato⁺ cells exhibit CD18$^{high}$IB4$^{high}$ surface phenotype consistent with axolotl macrophages.**
Flow cytometric characterization of FACS-sorted CD68:tdTomato$^{+}$ cells from pooled CD68:NTR 2.0-P2A-tdTomato transgenic limbs at 4 days post-amputation using established axolotl macrophage isolation technique by dual staining with anti-IB4 lectin and anti-CD18 myeloid markers (representative of limbs from N=3 separate animals, 8- to 10-month-old, ~10 cm length). Gating strategy shown on dot plots for debris removal (SSC-H versus FSC-H), doublet discrimination (FSC-A versus FSC-H), live cell inclusion (DAPI/BV421versus FSC-H), and identification of CD68:tdTomato populations (BV-421 versus tdTomato) classified by tdTomato expression level: negative, mid, and high. Control d/d (non-transgenic) pooled control sample processed under the same conditions demonstrates the specificity of the tdTomato fluorescent signal (Boxed in blue). Comparison of CD68:tdTomato$^{+}$ populations dual stained with anti-IB4 lectin and anti-CD18 shows macrophage specificity compared to CD68:tdTomato$^{neg}$ population. Antibodies: Fluorescein conjugated Isolectin B4 (IB4) (1:50, Vector labs cat. FK-1201), APC conjugated Mouse IgG1 anti-human CD18 (1:50, Biolegend cat. 302114).

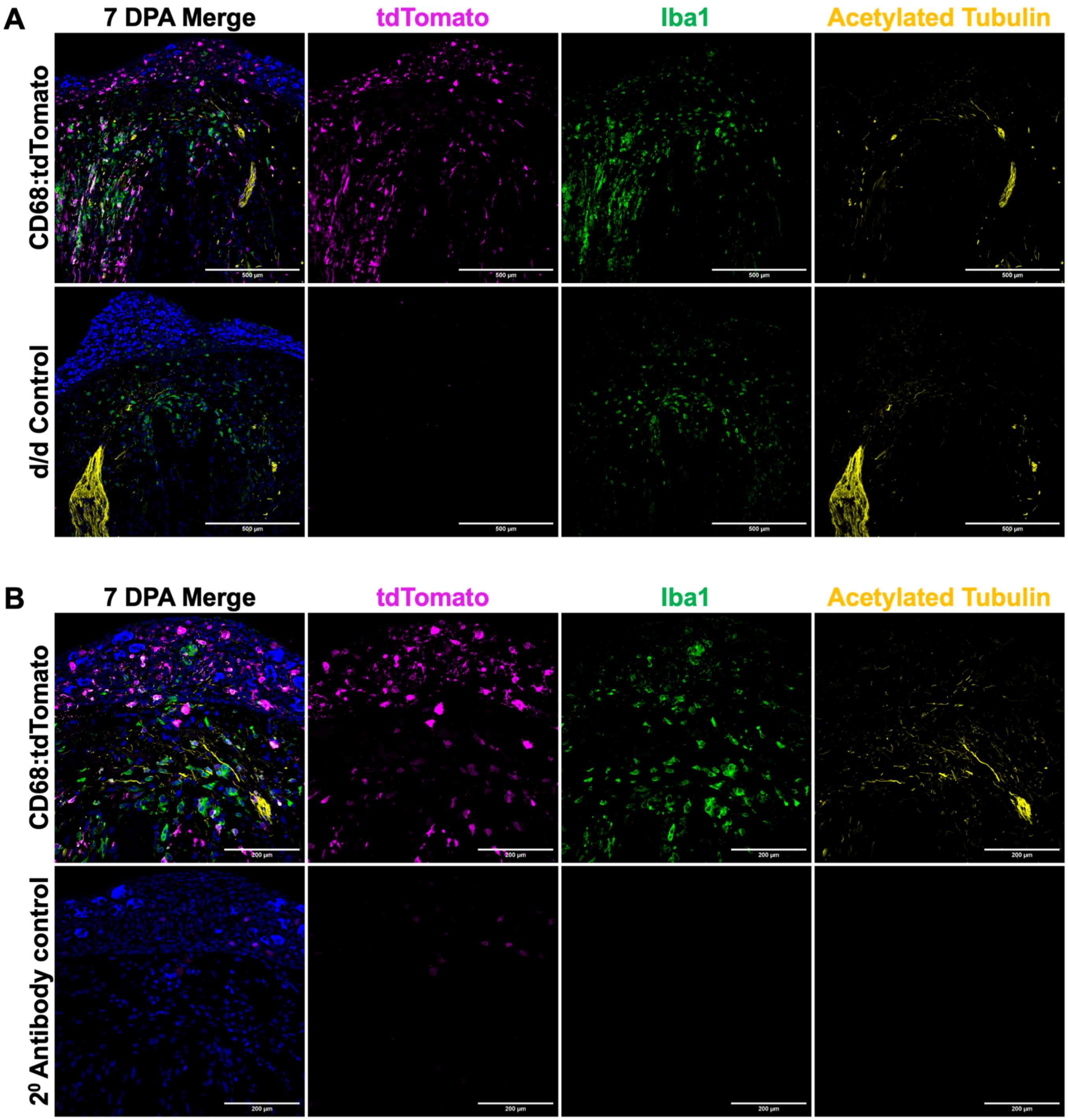


**Figure S17. Validation of CD68 transgene specificity in limbs at 7 days post-amputation via multi-marker immunofluorescence.**
**(A)** Confocal maximum intensity projections comparing transgenic CD68:NTR 2.0-P2A-tdTomato and d/d (non-transgenic) control limbs at 7 DPA. CD68:tdTomato$^{+}$ cells stained with anti-tdTomato antibody (magenta). Macrophages stained with anti-Iba1/AIF1 (green). Axons stained with Acetylated tubulin (yellow). Scale Bar: 500 µm.
**(B)** High-magnification confocal maximum intensity projections of 7 DPA CD68:NTR 2.0-P2A-tdTomato limbs stained with the complete antibody panel compared to the secondary antibody-only control demonstrates specificity of anti-tdTomato antibody (magenta) staining, co-expression with Iba1/AIF1 (green), and spatial association with acetylated tubulin$^{+}$ axons (yellow). Note that endogenous tdTomato is high in tissue resident macrophages but weak in monocytes requiring antibody amplification. Scale Bar: 200 µm. DPA=Days post-amputation. Antibodies: Rabbit anti-RFP (1:200, Rockland cat. 600-401-379); Chicken anti-Iba1 (1:50, Antibodies Inc. cat. IBA1-0020); Mouse IgG2b anti-acetylated tubulin (1:200, Sigma cat. T6793-100).

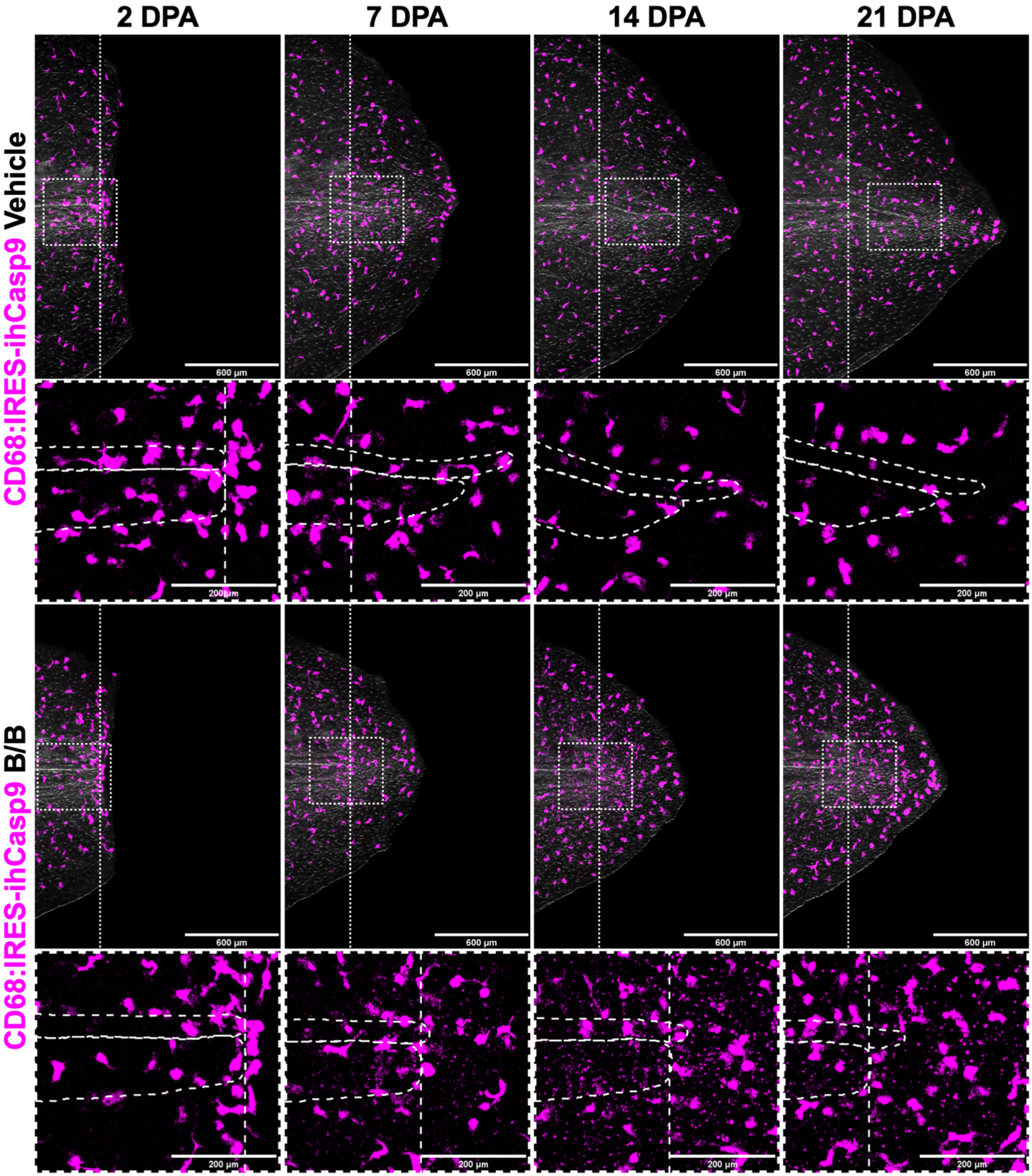


**Figure S18. Quantitative analysis of ihCasp9-mediated macrophage depletion effects on larval tail regeneration.** Representative extended depth of focus projections comparing vehicle- versus B/B-treated CD68:mScarlet-IRES-ihCasp9 larval tail regeneration at 2, 7, 14 and 21 DPA (N=3-5 animals per treatment, 2-month-old, 2-3 cm length). Amputation plane marked by white dotted line and white boxed regions mark inset locations. Insets of vehicle-treated tails show CD68$^{+}$ macrophage recruitment to the amputation plane (2 DPA) that appear to lead (are caudal to) spinal cord and cartilage regeneration (7, 14, and 21 DPA) in all cases examined. Insets of B/B-treated tails show decreased CD68$^{+}$ macrophage recruitment to the amputation plane (2 DPA) with an accumulation of fluorescent puncta indicative of apoptotic debris surrounding the regenerative zone (7, 14, and 21 DPA) relative to the vehicle control. Thin white dotted lines outline the spinal cord (dorsal) and cartilage (ventral). Scale Bar: 600 µm. Inset Scale Bar: 200 µm. B/B=AP20187 (ihCasp9 homodimerizer), DPA=Days post-amputation. Companion quantification shown in **Fig. 6**.

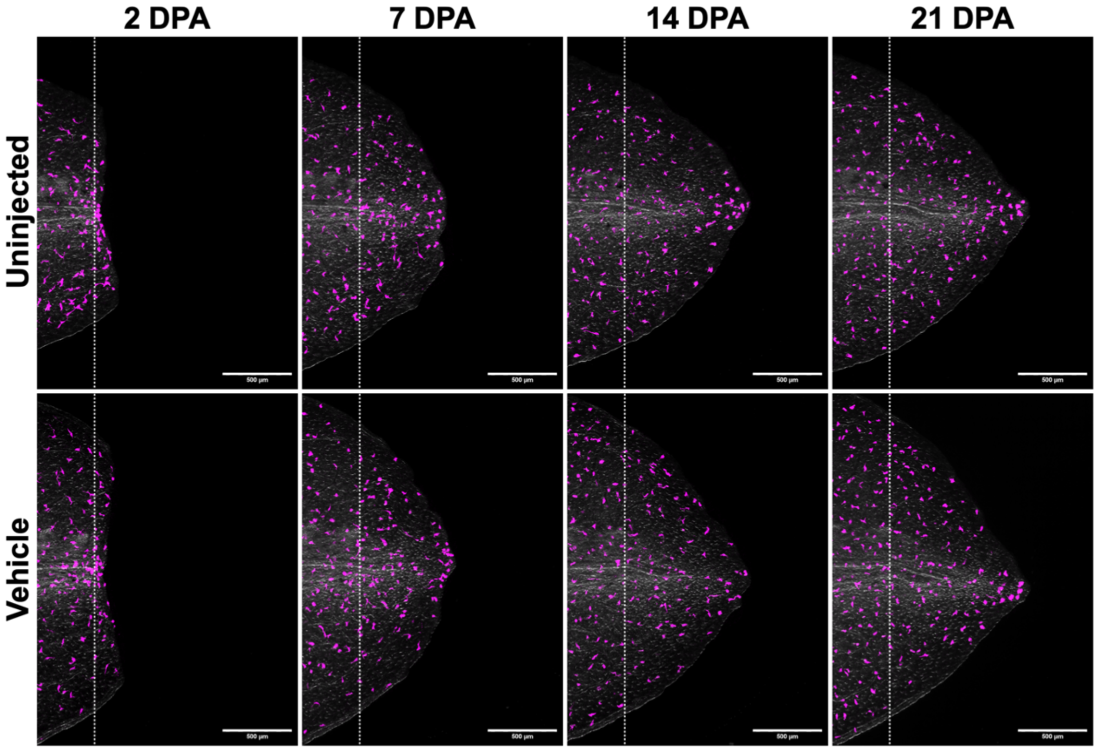


**Figure S19. Vehicle components alone do not affect tail regeneration in CD68:IRES-ihCasp9 larvae.**
Companion quantification for **Fig. 6.** Representative extended depth of focus projections demonstrate that vehicle administration (without B/B homodimerizer) does not impair cartilage, spinal cord, or epithelial outgrowth during CD68:mScarlet-IRES-ihCasp9 transgenic larval tail regeneration (N=3-5 animals per treatment, 2-month-old, 2-3 cm length). Comparison of uninjected and vehicle treatment groups at several post-amputation timepoints (2, 7, 14, and 21 days). Amputation plane marked by white dotted line. Scale Bar: 500 µm. DPA=Days post-amputation.

---

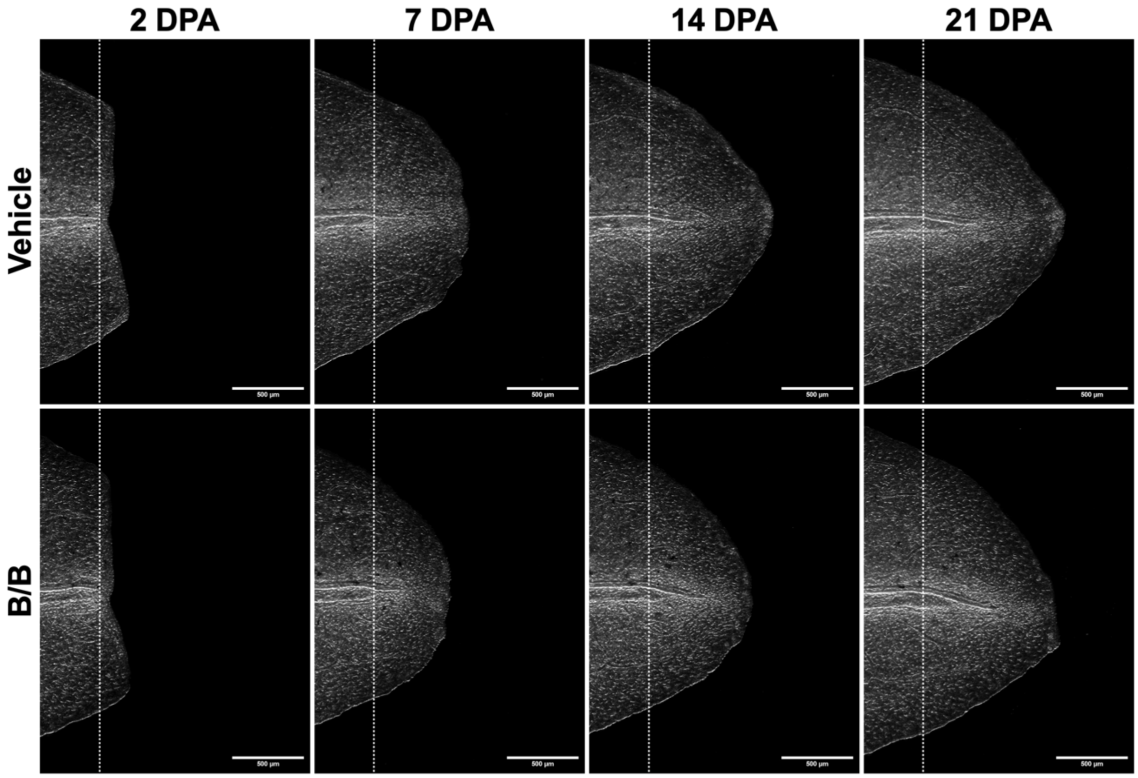


**Figure S20. B/B homodimerizer shows no off-target effects in non-sensitized control animals.**
Companion quantification for Figure 6. Representative extended depth of focus projections show comparable progression of d/d (non-sensitized) control larval tail regeneration at 2, 7, 14, and 21 DPA when treated with vehicle or 10 mg/kg AP20187 (B/B; ihCasp9 homodimerizer) via intraperitoneal injection (N=3 animals per treatment, 2-month-old, 2-3 cm length). Amputation plane marked by white dotted line. Scale Bar: 500 µm. DPA=Days post-amputation.

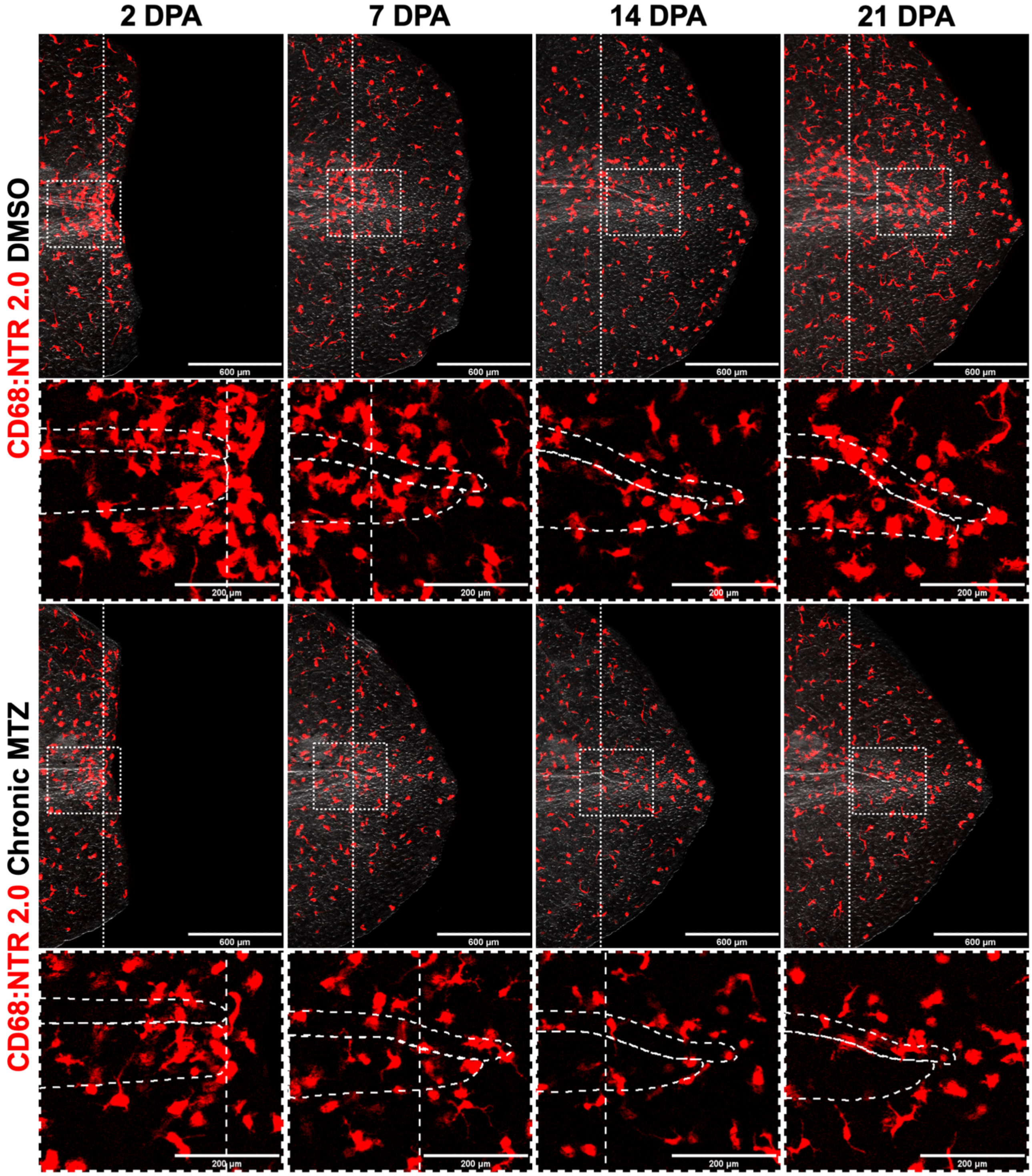


**Figure S21. NTR 2.0-mediated macrophage depletion via chronic MTZ impairs larval tail regeneration.**
Representative extended depth of focus projections comparing 1% DMSO vehicle versus chronic 1 mM MTZ-treated CD68:NTR 2.0-P2A-tdTomato larval tail regeneration at 2, 7, 14 and 21 DPA (N=5 animals per treatment, 2-month-old, 2-3 cm length). Amputation plane marked by white dotted line and white boxed regions mark inset locations. Insets of DMSO-treated tails show CD68$^{+}$ macrophage recruitment to the amputation plane (2 DPA) to caudal end of spinal cord (7, 14, and 21 DPA). Insets of MTZ-treated tails show decreased CD68$^{+}$ macrophage recruitment to the amputation plane (2 DPA) with progressively rounding CD68$^{+}$ morphology (7 and 14 DPA) that starts to recover after a 7-day washout period (21 DPA) relative to the vehicle control. Thin white dotted lines outline the spinal cord (dorsal) and cartilage (ventral). Scale Bar: 600 µm. Inset Scale Bar: 200 µm. MTZ=Metronidazole (NTR 2.0 prodrug), WB=Waterborne, DPA=Days post-amputation. Companion quantification shown in **Fig. 6**.

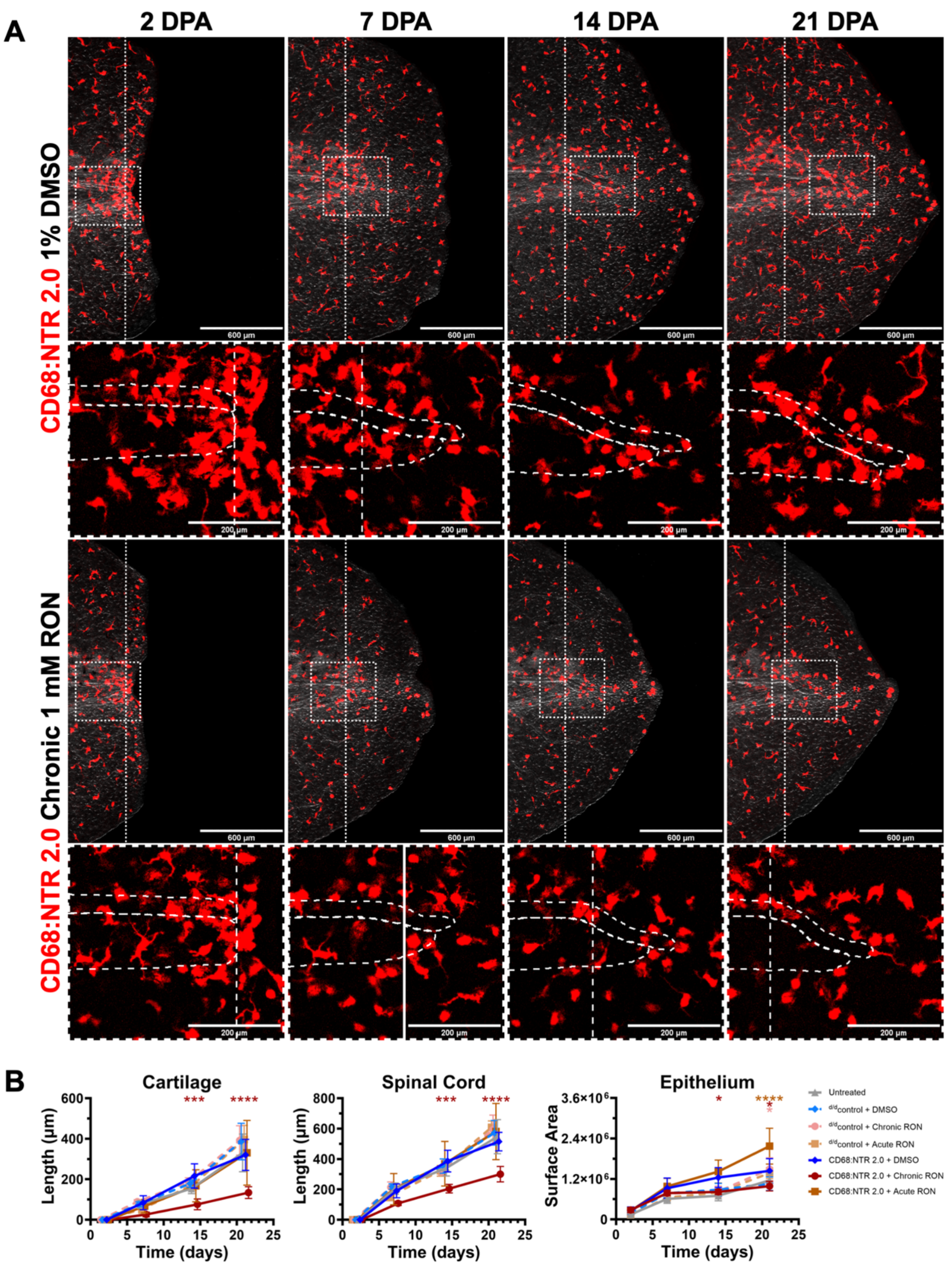


**Figure S22. Ronidazole-mediated NTR 2.0 ablation inhibits larval tail regeneration.**
Companion data for **Fig. 6**. Note that the same experimental design was used as figure 6 using RON in place of MTZ.
**(A)** Representative extended depth of focus projections comparing 1% DMSO vehicle versus chronic 1 mM RON-treated CD68:NTR 2.0-P2A-tdTomato larval tail regeneration at 2, 7, 14 and 21 DPA (N=5 animals per treatment, 2-month-old, 2-3 cm length). Amputation plane marked by white dotted line and white boxed regions mark the inset location. Insets of DMSO-treated tails show $CD68^{+}$ macrophage recruitment to the amputation plane (2 DPA) that appear to lead (are caudal to) spinal cord and cartilage regeneration (7, 14, and 21 DPA). Insets of RON-treated tails show decreased $CD68^{+}$ macrophage recruitment to the amputation plane (2 DPA) with progressively rounding $CD68^{+}$ morphology (7 and 14 DPA) that starts to recover after a 7-day washout period (21 DPA) relative to the vehicle control. Thin white dotted lines outline the spinal cord (dorsal) and cartilage (ventral). Scale Bar: 600 µm. Inset Scale Bar: 200 µm. RON=Ronidazole (NTR 2.0 prodrug), WB=Waterborne, DPA=Days post-amputation.
**(B)** Quantitative morphometric analysis of tail regenerates in CD68:NTR 2.0-P2A-tdTomato transgenics (solid lines) and non-sensitized d/d controls (dashed lines) treated with WB RON immersion. Measurements of cartilage length, spinal cord length, and epithelial surface area that extend beyond the amputation plane were collected at 2, 7, 14, and 21 DPA in Fiji (N=5 animals per treatment, 2-month-old, 2-3 cm length). Error bars represent mean ± SD. Statistical significance for both RON treatments relative to the DMSO vehicle at each regenerative timepoint was determined by two-way ANOVA with Tukey's multiple comparisons test (*$p$<0.05, ***$p$<0.001, ****$p$<0.0001).

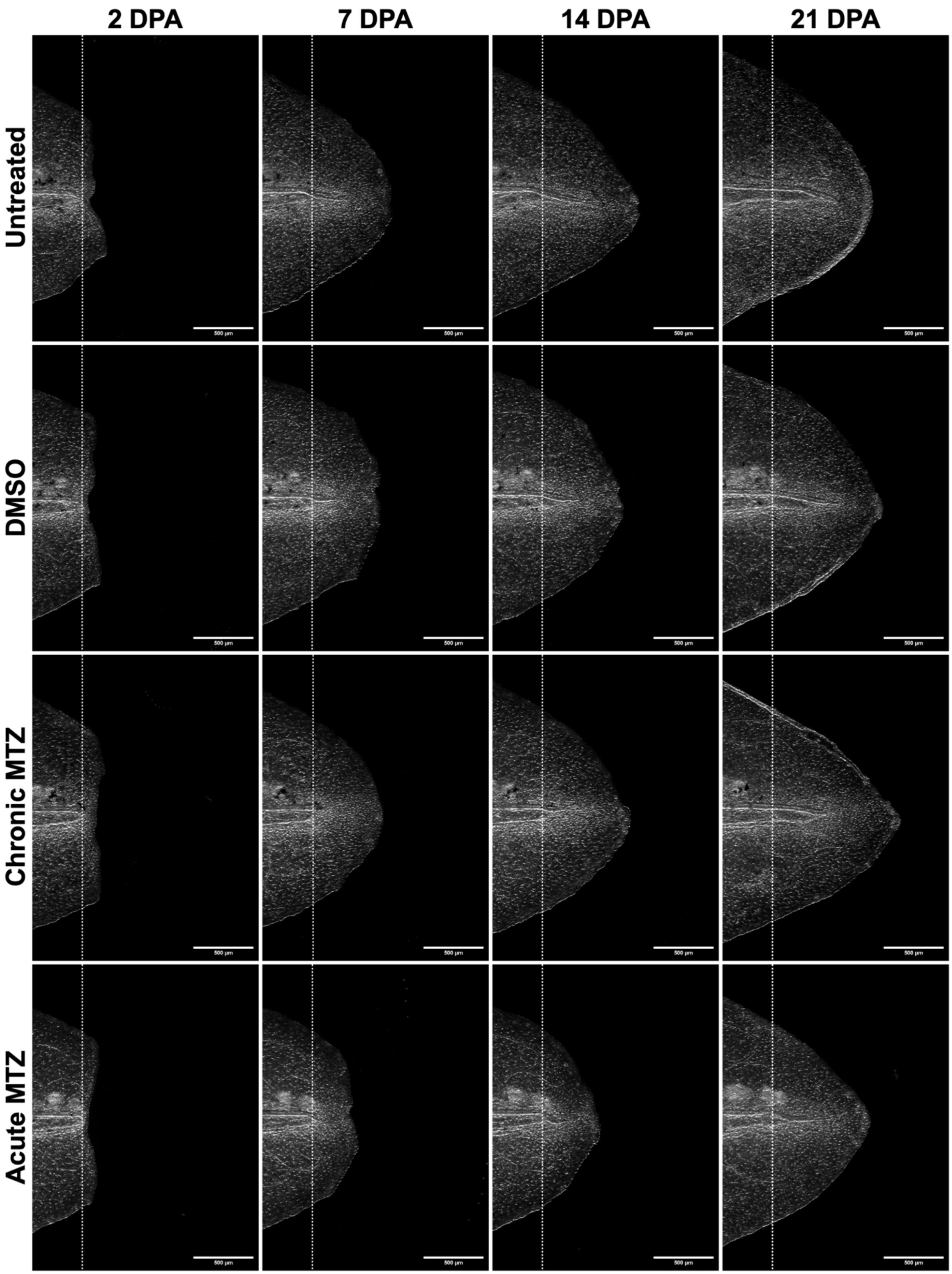


**Figure S23. Acute high-dose MTZ causes off-target regeneration defects in non-sensitized control larvae.**
Companion data for **Fig. 6**. Representative extended depth of focus projections of non-sensitized d/d control animals at 2, 7, 14, and 21 DPA demonstrate that 1% DMSO and chronic 1 mM MTZ were well-tolerated compared to the untreated animals, whereas four acute 10 mM MTZ WB immersions delays regeneration, indicating off-target MTZ prodrug toxicity at high concentrations in larval stage animals (N=5 animals per treatment, 2-month-old, 2-3 cm length). Scale Bar: 500 µm. MTZ=Metronidazole (NTR 2.0 prodrug), DPA=Days post-amputation.

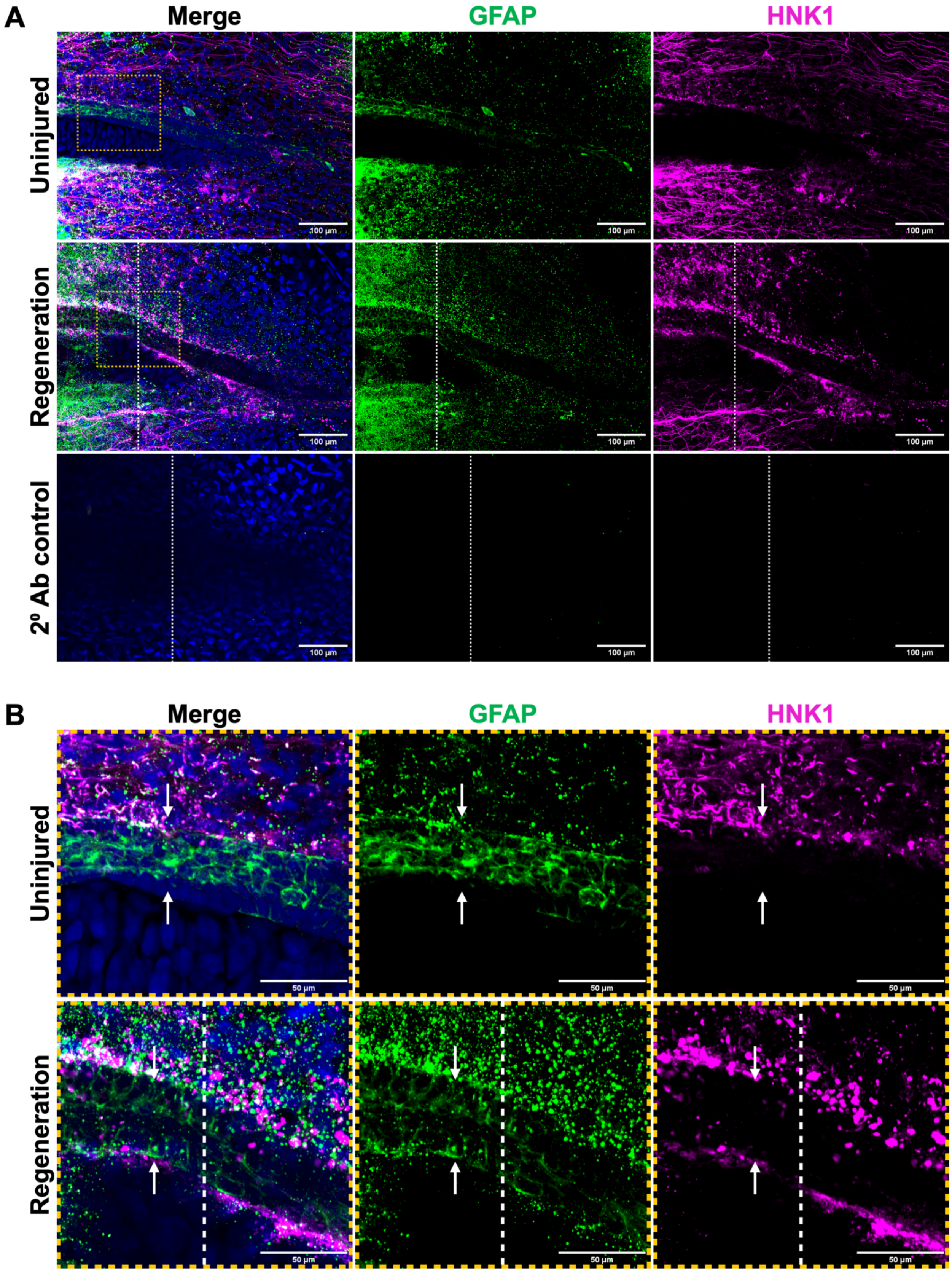


**Figure S24. HNK1⁺ and GFAP⁺ populations show partial overlap in regenerating tail CNS.**
**(A)** Representative confocal maximum intensity projections of whole-mounted uninjured and 21 DPA wild-type d/d tails shows the spatial relationship between GFAP⁺ glial/ependymal cells (green) and HNK1⁺ neural crest-derived cells (magenta) with specificity compared to the secondary (2º) antibody control (N=2 animals per timepoint, 2-month-old, 2-3 cm length). Amputation plane marked by white dotted line and orange boxed regions mark the inset location. **(B)** Insets (orange boxes) of the spinal cord in uninjured versus 21 DPA wild-type d/d tails show an increase in HNK1⁺ and decrease in GFAP⁺ expression inside the ependymal tube (dorsal/ventral border of tube shown with white arrows) during regeneration. Scale Bar: 100 μm. Inset scale Bar: 50 μm. DPA=Days post-amputation. Antibodies: Mouse IgG1 anti-GFAP (1:200, Sigma cat. MAB360), Mouse IgM anti-HNK1 (1:200, Sigma cat.C6680).

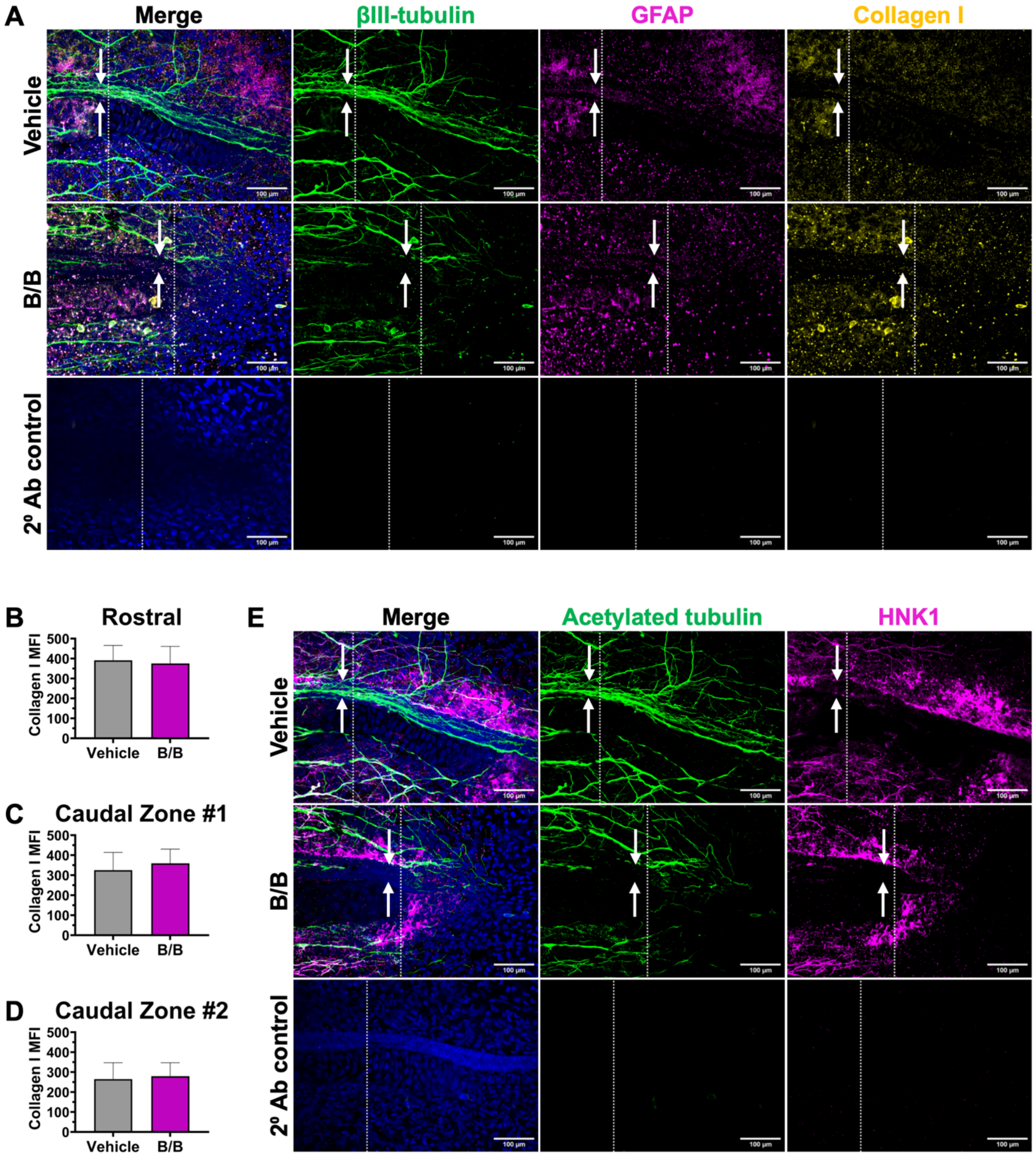


**Figure S25. Regenerative failure in macrophage-depleted tails is not associated with excessive collagen deposition and features a reduction in GFAP$^+$ cells inside the ependymal tube but normal numbers outside.**

**(A)** Representative confocal maximum intensity projections of whole-mounted CD68:mScarlet-IRES-ihCasp9 tails at 21 DPA (vehicle-versus B/B-treated) stained for βIII-tubulin$^+$ axons (green), GFAP$^+$ cells (magenta), and collagen I (yellow) compared to secondary (2º) antibody control (N=3 animals per treatment, 2-month-old, 2-3 cm length). Amputation plane marked by white dotted line. Dorsal ventral boundary of ependymal tube marked with white arrows. Scale Bar: 100 µm.

**(B-D)** Quantification of MFI for collagen I staining in 21 DPA whole-mounted CD68:tdTomato-IRES-ihCasp9 tails (vehicle- versus B/B-treated) within three defined 200 µm regions relative to the amputation plane: rostral zone (adjacent to the amputation plane, stump)**(B)**, caudal zone #1 (adjacent to the amputation plane, proximal regenerate) **(C)**, or caudal zone #2 (distal regenerate) **(D)** (N=3 animals per treatment, 2-month-old, 2-3 cm length). Error bars represent mean ± SD. No statistically significant difference was observed in collagen I MFI between vehicle and B/B-treated animals across all defined regions (unpaired t-test, significance threshold of $p<0.05$).

**(E)** Representative confocal maximum intensity projections of whole-mounted CD68:tdTomato-IRES-ihCasp9 tails at 21 DPA (vehicle versus B/B-treated) co-stained with acetylated tubulin$^+$ axons (green) and HNK1$^+$ cells (magenta) compared to secondary (2º) antibody control (N=3 animals per treatment, 2-month-old, 2-3 cm length). Amputation plane marked by white dotted line. Dorsal ventral boundary of ependymal tube marked with white arrows. Scale Bar: 100 µm.

B/B=AP20187 (ihCasp9 homodimerizer), DPA=Days post-amputation. Antibodies: Mouse IgG2a anti-tubulin βIII (TUBB3) (1:200, Biolegend cat. 801202), Mouse IgG1 anti-GFAP (1:200, Sigma cat. MAB360), Rabbit anti-collagen type I (1:200, Millipore cat. AB755P).

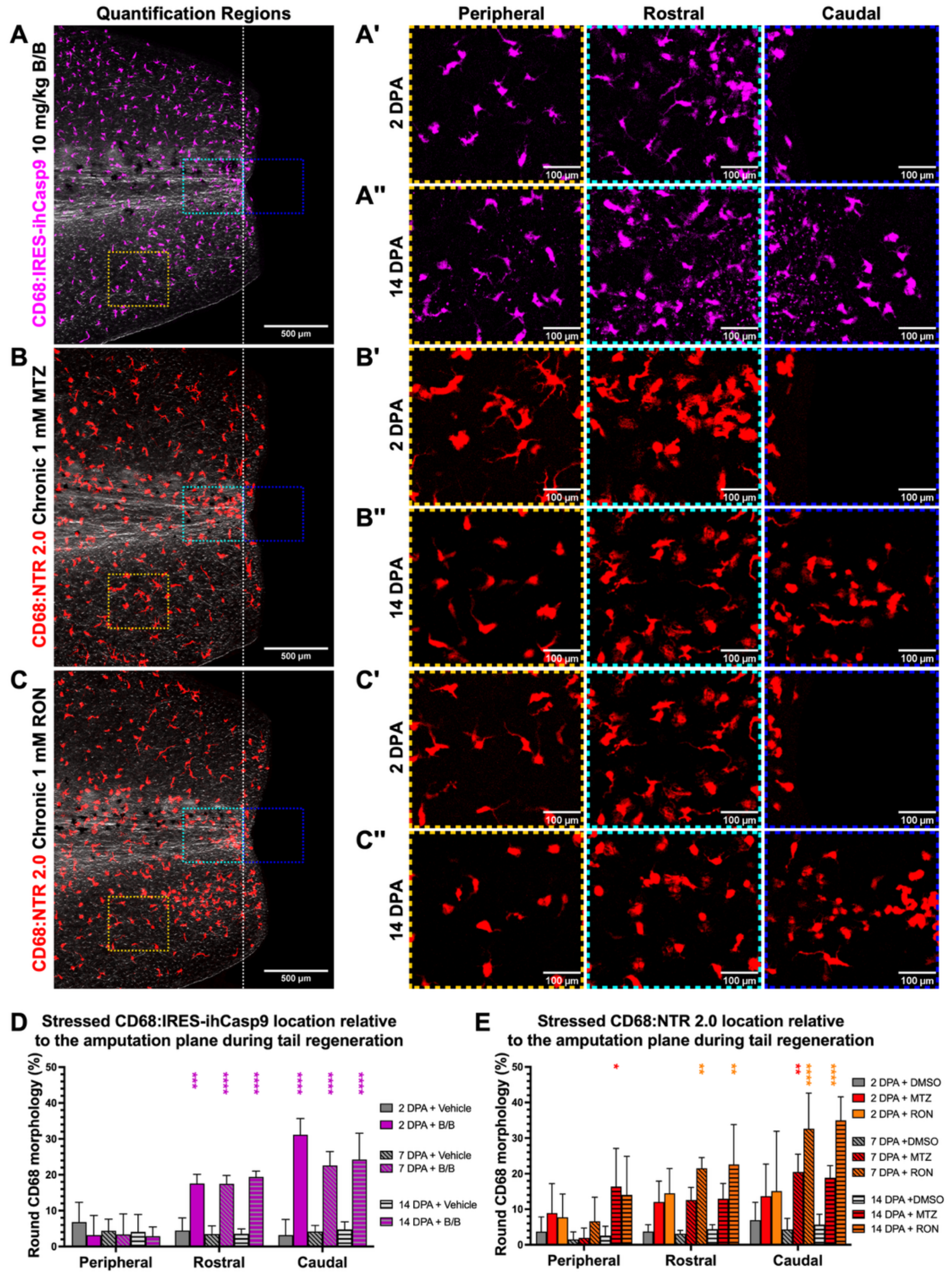


**Figure S26. Spatial analysis reveals route-dependent macrophage ablation patterns.**

**(A-C)** Representative extended depth of focus projections at 2 DPA show tail quantification regions for CD68$^{+}$ morphology in **(A)** B/B-treated CD68:mScarlet-IRES-ihCasp9 transgenics and CD68:NTR 2.0-tdTomato transgenics treated with either **(B)** 1 mM MTZ or **(C)** 1 mM RON via chronic WB immersion (N=3-5 animals per treatment, 2-month-old, 2-3 cm length). Quantification regions were defined as the rostral zone (stump, adjacent to amputation plane), caudal zone (regenerate, beyond the amputation plane), and peripheral zone (skin-associated, remote location from amputation site). **(A'-C')** Insets show predominantly dendritic CD68$^{+}$ morphology in the rostral (cyan box), caudal (blue box), and peripheral (orange box) zones at 2 DPA. **(A''-C'')** Insets show more rounded CD68$^{+}$ morphology at 14 DPA in the rostral (cyan box), caudal (blue box), and peripheral (orange box) zones associated with prolonged inducible drug exposure relative to 2 DPA timepoint. Amputation plane marked by white dotted line.

**(D)** Quantification of stressed/rounded macrophage morphology (indicative of drug-induced apoptosis) in CD68:mScarlet-IRES-ihCasp9 animals following *i.p.* B/B delivery at 2, 7, and 14 DPA (N=3-5 animals per treatment, 2-month-old, 2-3 cm length). Bar graphs show region-specific differences where stressed macrophages were seen in the highly vascularized rostral and caudal zones at all measured timepoints. Error bars represent mean ± SD. Statistical significance of B/B treatment relative to vehicle-treated animals within each quantification region was determined by two-way ANOVA with Tukey's multiple comparisons test (***$p$<0.001, ****$p$<0.0001).

**(E)** Quantification of stressed/rounded macrophage morphology in CD68:NTR 2.0-P2A-tdTomato animals following WB immersion-based chronic 1 mM MTZ/RON delivery shows contrasting spatial patterns with greater peripheral ablation at 2, 7, and 14 DPA (N=5 animals per treatment, 2-month-old, 2-3 cm length). Error bars represent mean ± SD. Statistical significance of treatment with MTZ or RON relative to DMSO-treated animals within each quantification region was determined by two-way ANOVA with Tukey's multiple comparisons test (*$p$<0.05, **$p$<0.01, ****$p$<0.0001).

Scale Bar: 500 µm. Inset Scale Bar: 100 µm. B/B=AP20187 (ihCasp9 homodimerizer), *i.p.*=Intraperitoneal, MTZ=Metronidazole (NTR 2.0 prodrug), RON=Ronidazole (NTR 2.0 prodrug), WB=Waterborne, DPA=Days post-amputation.

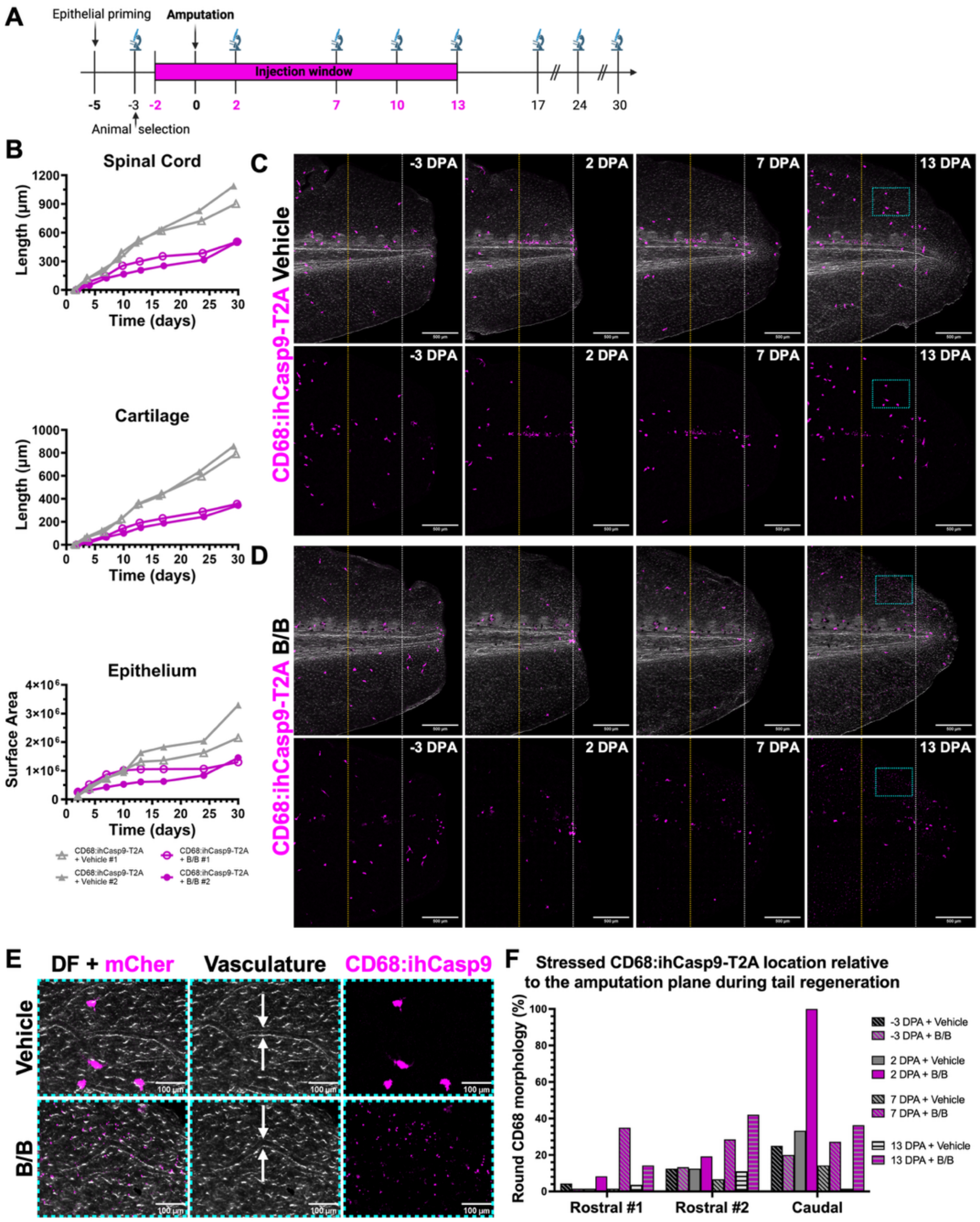


**Figure S27. Mosaic F0 CD68:ihCasp9-T2A-tdTomato animals demonstrate enhanced ablation potency.**

**(A)** Experimental timeline for macrophage ablation in mosaic CD68:ihCasp9-T2A-tdTomato F0 transgenic axolotls. The timepoints with *i.p.* drug delivery of B/B shown in magenta. Imaging schedule shown with microscope icon and time relative to amputation.
generation, epithelial priming, tail amputation, B/B *i.p.* injection regimen, and imaging schedule.

**(B)** Quantification of individual cartilage length, spinal cord length, and epithelial surface area in high-expressing F0 mosaic CD68:ihCasp9-T2A-tdTomato pairs treated with either vehicle (solid gray lines) or B/B (solid magenta lines) demonstrate that partial macrophage ablation (≤50% of population) is sufficient to impede tail regeneration (N=2 per treatment, paired based on the percentage of fluorescent macrophage labeling, 2-month-old, 2-3 cm length).

**(C, D)** Representative extended depth of focus projections comparing vehicle-treated **(C)** versus B/B-treated **(D)** mosaic CD68:ihCasp9-T2A-tdTomato animals at -3, 2, 7, and 13 DPA (N=2 per treatment, paired based on the percentage of fluorescent macrophage labeling, 2-month-old, 2-3 cm length). Amputation plane indicated by white dotted line and cyan boxed regions mark the inset locations. Orange dotted line divides the rostral region into zone #1 (furthest from the amputation plane) and zone #2 (adjacent to the amputation plane) for quantification of CD68$^{+}$ morphology (Figure S27D). Scale Bar: 500 µm. Cyan dashed box shows inset location for E.

**(E)** Representative extended depth of focus projections showing insets of B/B-induced macrophage destruction near vasculature (what arrows show example vessel) with characteristic fluorescent puncta (apoptotic debris) absent in vehicle controls at 13 DPA (N=3 per treatment, paired based on the percentage of fluorescent macrophage labeling, 2-month-old, 2-3 cm length Inset Scale Bar: 100 µm.

**(F)** Spatial quantification of stressed macrophages in mosaic F0 CD68:ihCasp9-T2A-tdTomato transgenic animals showing enhanced peripheral ablation compared to germline F1 CD68:IRES-ihCasp9 transgenic line (Figure S26), consistent with improved 1:1 stoichiometry of T2A architecture (N=1 per treatment, 2-month-old, 2-3 cm length). Quantification was performed at -3, 2, 7, and 13 DPA within defined 750 µm zones: Rostral zone #1 (furthest from the amputation plane), Rostral zone #2 (immediately adjacent to the amputation plane), and caudal zone (beyond the amputation plane). Note: Majority of B/B-affected macrophages had completed apoptosis by measurement timepoints and were not included in morphological analysis, evidenced by abundant puncta expression. Data are presented descriptively without statistical analysis. B/B=AP20187 (ihCasp9 homodimerizer), *i.p.*=Intraperitoneal, DPA=Days post-amputation.
